# Brain rhythms during Instagram Reels viewing and subsequent performance on sustained attention and working memory: an EEG study

**DOI:** 10.64898/2026.11.26.769379

**Authors:** Zahra Ramezani, Amin Mohammad Mohammadi, Zoha Hajiha, Behrooz Dolatshahi

## Abstract

What does even a short scrolling ‘snack’ do to the brain? Short-form video feeds such as Instagram Reels have been rapidly becoming a near-universal phenomenon, used widely by people around the world, especially young adults. However, this is a very recent development, and yet their immediate effects on brain activity, attention, and working memory remain unclear. In a counterbalanced, within-subject experiment, we examined whether 10 minutes of Instagram Reels altered electroencephalographic (EEG) activity, cognitive performance, and subjective fatigue. Thirty university students completed three testing blocks separated by two counterbalanced interventions selected from viewing Reels on their own accounts, viewing a silent sequence of neutral images on their phones, or resting with eyes closed. EEG was recorded using 65 electrodes during interventions and subsequent sustained-attention (SART) and working-memory (N-back) tasks. Compared with neutral images, Reels viewing was associated with higher posterior relative theta power and a lower posterior alpha-to-theta ratio. During the subsequent N-back, frontal relative alpha was higher after Reels than after neutral images. Fatigue increased more after Reels than after rest. One of four planned contrasts in the cleaned sample was significant. Brief Reels viewing was associated with differences in EEG activity compared with viewing neutral images, while no significant differences in attention or working-memory performance were detected after correction for multiple comparisons.

## Introduction

Social media are used by most of the world’s population. At the start of October 2026, 5.81 billion social media user identities were reported, equivalent to 69.9% of the global population, and the typical internet user spent about 19 hours a week, or more than 2.5 hours a day, on social media and video platforms [1]. Instagram is among the most widely used of these platforms: 56.6% of internet users aged 16 years and over reported having used it in the past 30 days [1]. Short-form video, in which users scroll through a continuous feed of brief clips, has become a central format on such platforms, with TikTok, Instagram Reels, and YouTube Shorts as the main examples [2,3].

Social media bring clear benefits: they help people keep in touch with friends and family, give access to information and peer support, including for people living with mental illness [4], and are used for teaching and learning in higher education [5]. At the same time, their rapid growth, and especially the spread of short-form video, has raised concerns about their effects on mental health and cognition.

Most research on these concerns has examined habitual use rather than brief, controlled exposure. For well-being, the associations found so far are small. In three large adolescent datasets (*n* = 355,358), digital technology use explained at most 0.4% of the variation in well-being [6], and a review of more than 80 reviews and meta-analyses concluded that most studies are cross-sectional and of low quality and that the average association is negative but very small, with its direction still unclear [7]. For cognition, heavy smartphone and internet use has been linked with weaker sustained attention, greater distractibility, and poorer working memory [8,9], and problematic internet use with poorer inhibitory control and working memory [10]. For short-form video specifically, a meta-analysis of 71 studies with 98,299 participants found that heavier use was associated with poorer attention (*r* = −.38) and poorer inhibitory control (*r* = −.41) [3], and a systematic review of 42 studies in young people reported similar associations with inattention, impulsivity, and working memory [11]. However, nearly all of this evidence comes from studies that measure habitual use and cognition at the same point in time. Such studies cannot show which comes first: people with weaker attentional control may be drawn to fast-changing content, rather than the reverse, or a third factor, such as poor sleep or low mood, may affect both. Nor can they show whether a single viewing session changes brain activity or performance in the minutes that follow, which is the question an experiment can answer.

Experiments on the immediate effects of a single, brief exposure are fewer, and their results differ (Table 1). In two experiments, 45 min of free smartphone use slowed responses on a vigilance task and increased errors on No-Go trials compared with watching a documentary for the same length of time; both activities increased subjective mental fatigue, but only smartphone use impaired performance [12]. Fifteen minutes of scrolling one’s own Instagram feed lowered n-back and Go/No-Go accuracy and altered prefrontal activation measured with functional near-infrared spectroscopy, compared with scrolling a feed of black- and-white nature images [13]. In another experiment, participants performed a prospective-memory task, which requires remembering to carry out a planned action, before and after a 10-min break. Spending the break on TikTok impaired prospective memory relative to resting, whereas spending it on Twitter or YouTube did not [14]. When participants could switch between short videos as often as they liked during a 10-min break, prospective memory worsened afterwards, whereas when the number of switches was limited, it improved [15]. After 30 min of TikTok-style videos, participants no longer benefited from longer preparation intervals in task switching [16], and learning from several short videos led to poorer recall than learning from one long video of the same duration and content [17]. In children, fast-paced or fast-edited video impaired executive function, increased shifts of attention, and altered the N2 and P3 components during response inhibition [18–20]. Other experiments found no effect of a short social media session on working memory [21] or on attention, working memory, reasoning, and inhibition [22]. These studies differ in participants’ age, exposure duration (about 4-45 min), content, comparison condition, and outcome, so they cannot yet be combined into a single estimate of the short-term effect.

**Table 1.** Experiments on the Short-Term Cognitive Effects of Social Media, Short-Form Video, and Fast-Paced Audiovisual Content.

| Study | Participants | Exposure (duration) | Comparison | Outcome measures | Main finding |
|---|---|---|---|---|---|
| Lillard & Peterson (2011) [18] | 60 children aged 4 years | Fast-paced cartoon (9 min) | Educational cartoon; drawing | Executive-function tasks (e.g., delay of gratification, Tower of Hanoi) | Poorer executive function after the fast-paced cartoon |
| Kostyrka-Allchorne, Cooper, Gossmann, et al. (2017) [19] | 70 children aged 2-4.5 years | Film with fast editing (4 min) | Same story with slow editing | Free play (attention shifts between toys) | More attention shifts after the fast-edited film |
| Kostyrka-Allchorne et al. (2019) [20] | 40 children aged 7 years | Fast-paced video (4 min) | Slow-paced video | Go/No-Go; N2 and P3 event-related potentials | More No-Go errors and altered N2/P3 timing after the fast-paced video |
| Almarzouki et al. (2022) [21] | 118 university students | Own social media (6-min block) | Online painting (within person) | CANTAB Spatial Working Memory | No overall difference; more errors after social media in participants with at least moderate depression |
| Chiossi et al. (2023) [14] | 60 adults (15 per condition) | TikTok, Twitter, or YouTube (10 min) | Rest without screens | Prospective memory within a lexical decision task | Prospective memory impaired after TikTok only |
| Jacquet et al. (2023) [12] | 2 experiments, 40 per group | Free smartphone use (45 min) | Documentary | Psychomotor Vigilance Task; Go/No-Go; fatigue and drowsiness | Slower vigilance responses and more No-Go errors after smartphone use; fatigue rose in both groups |
| Aitken et al. (2025) [13] | 20 college students | Own Instagram feed (15 min) | Feed of black-and-white nature images | n-back; Go/No-Go; fNIRS | Lower task accuracy and altered prefrontal activation after Instagram |
| Luo et al. (2025) [16] | 72 adults (3 groups) | TikTok-style short videos (30 min) | Documentary; no video | Cued task switching with varied preparation intervals | No preparation benefit after short videos; benefit preserved after documentary or no video |
| Barton & Smyth (2025) [15] | 45 adults | Short-form videos with unlimited or limited (10) switches (10 min) | No video (10-min wait) | Prospective memory within a lexical decision task | Worse prospective memory with unlimited switching, better with limited switching |
| Mendonça et al. (2025) [22] | 66 adolescents aged 13-15 years | Social media use (30 min) | Face-to-face conversation | d2 attention; Corsi blocks; abstract reasoning; Go/No-Go | No differences in any domain |
| Wei et al. (2026) [17] | 57 participants | Several short videos (10 min) | One long video matched for duration and content | Recall; fMRI during retrieval | Poorer recall and reduced claustrum, caudate, and middle temporal activation after short videos |

*Note.* fMRI, functional magnetic resonance imaging; fNIRS, functional near-infrared spectroscopy.

Several features of this literature may explain the disagreement. First, many studies treat social media use as a single exposure, although text, images, long videos, and short videos place different demands on the user. When platforms were compared directly, only TikTok impaired prospective memory [14], and how often participants could switch between videos determined whether prospective memory worsened or improved [15]. On Instagram, like TikTok, the Reels feed delivers a continuous, algorithmically personalized sequence of short videos that the user advances by scrolling. Personalized short videos activate reward-related and default-mode regions more strongly than non-personalized videos [23], and preferred videos that are watched to the end deactivate the dorsal anterior cingulate and dorsolateral prefrontal cortex [24]. Studying one platform and one format makes it possible to define the exposure precisely and to choose a control condition that matches its non-social features, such as holding a phone and looking at its screen. Second, most studies use between-group designs, in which large differences between people in cognitive ability, habitual use, and resting EEG can hide small effects; within-subject designs remove these stable differences from the comparison [25].

Third, behavioral measures such as accuracy and reaction time can be close to ceiling, and many widely used tasks have low between-person reliability [26]. EEG provides complementary information: it records brain activity continuously, both during exposure and during later task performance, and it can reveal changes in processing when performance is unchanged. EEG signals predict lapses of sustained attention up to 20 s before errors occur [27], and theta, alpha, and beta activity change during mind wandering [28]. Alpha oscillations (8-13 Hz) dominate the posterior EEG when the eyes are closed and decrease when the eyes open [29,30]. Lower alpha over a region is often taken to reflect reduced inhibition and greater processing of incoming input, and higher alpha reflects the gating of information [31–33]. Theta oscillations (4-8 Hz) have been linked to encoding and working memory, and frontal-midline theta to cognitive control [34,35]. Because rising task demands often increase theta and decrease alpha, the alpha/theta ratio is used to describe the balance between the two bands, although it is an indirect measure whose meaning depends on the task and the region. Power above about 30 Hz is particularly susceptible to muscle activity [36,37]. EEG studies of short-form video have so far been correlational, linking heavier use to a smaller P300 [2] and to lower frontal theta during conflict trials [38].

Attention and working memory are plausible targets of short-form video. Inhibition and working memory are core executive functions [39], and the strongest correlates of short-form video use identified so far concern attention and inhibitory control [3]. A feed that changes every few seconds may repeatedly capture attention and displace what is held in mind. The Sustained Attention to Response Task (SART) measures sustained attention and response inhibition [40], and the N-back task measures the maintenance and updating of working memory at several load levels [41].

The present study examined how 10 min of Instagram Reels affects EEG activity during viewing and during subsequent attention and working-memory tasks. Reels were watched on participants’ own accounts so that the feed was personalized by the platform’s algorithm, as in everyday use [23], and were compared with a silent sequence of neutral images on the phone, which controlled for holding a phone and looking at its screen, and with eyes-closed rest, which provided a resting reference. Because there was little prior evidence on how EEG activity changes during or after short-form video viewing, no directional hypotheses were formulated. Instead, four planned contrasts, based on the literature linking frontal-midline theta to cognitive control and posterior alpha to visual processing and attention, were designated as primary tests. During the interventions, frontal-midline theta (C-I1) and occipital alpha (C-I2) were compared between Reels and neutral images. During the subsequent tasks, occipital alpha was compared between Reels and rest (C-T1), and frontal-midline theta between Reels and neutral images (C-T2). The wider spectral profile was examined in exploratory analyses, and the study asked whether N-back and SART performance and subjective fatigue after Reels differed from those after neutral images and after rest.

## Results

Each of 30 participants completed three testing blocks separated by two 10-min interventions and so received two of the three conditions: Instagram Reels (E), neutral images (N), or eyes-closed rest (R) (Fig. 1). Each intervention was therefore completed by 20 participants, and 10 participants completed both interventions of each pair. EEG was recorded continuously during the interventions and during the testing block that followed each intervention, which comprised the SART, the N-back task, and a fatigue rating. EEG results are reported for the full sample (*N* = 30) and for a cleaned sample that excluded the two participants with the highest noise scores (*N* = 28).

**Figure 1.**
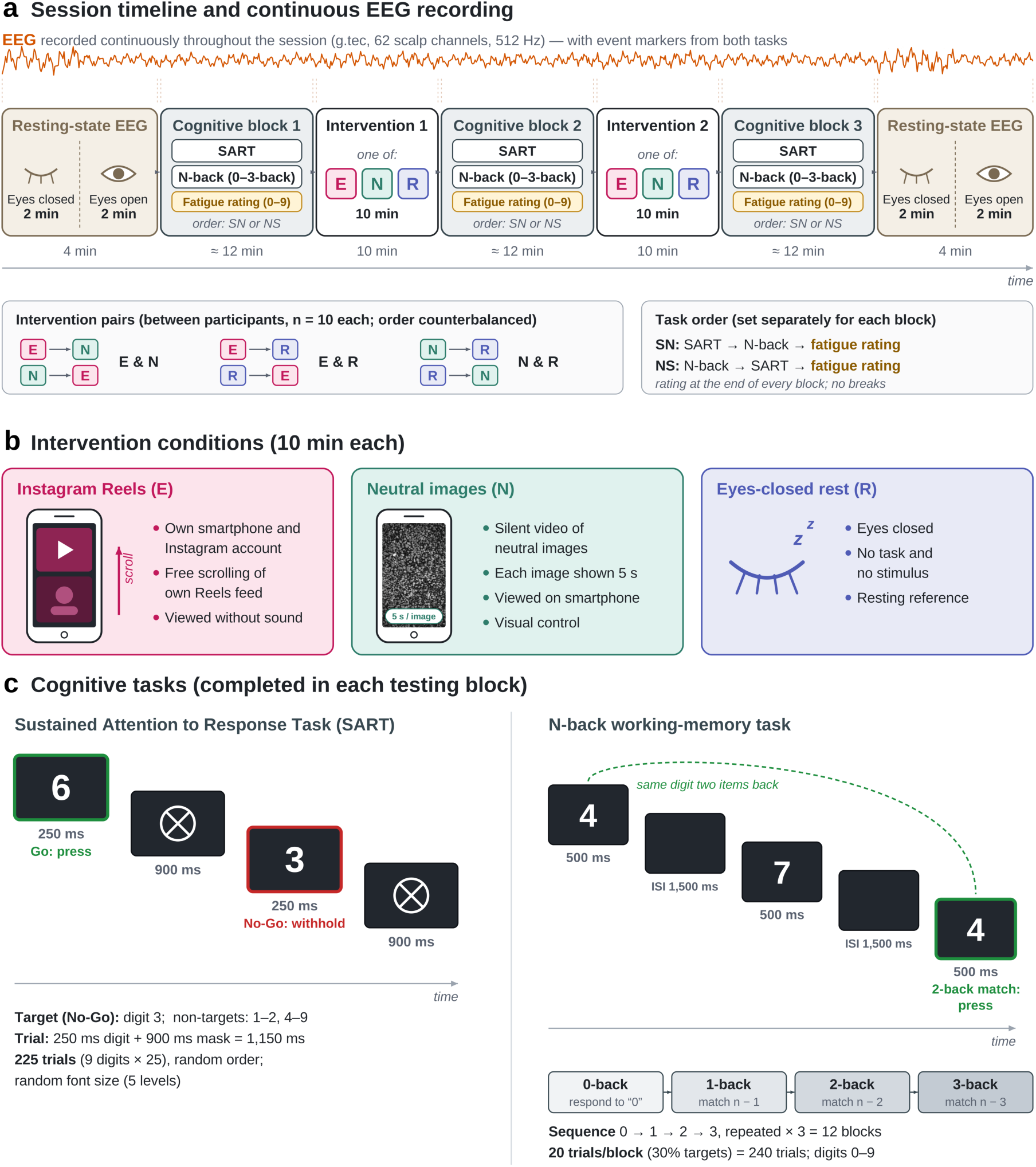
Experimental paradigm. (a) Session timeline. EEG was recorded continuously from the first to the last resting-state recording. Each participant received the two interventions of one condition pair in counterbalanced order; the task order (SART then N-back, SN, or the reverse, NS) was set for each block, and fatigue was rated at the end of every block. (b) The three intervention conditions. (c) Trial structure of the SART (left) and of the N-back task, shown with a 2-back example, together with the sequence of load levels (right).

### Sample and data quality

All 30 participants completed the session. Because each participant received two interventions, the data comprised 60 intervention recordings and 60 post-intervention testing blocks, 20 of each per condition. The intervention recordings were divided into 16,639 overlapping 4-s windows, of which 14,381 (86.4%) passed the artifact criteria and were retained, with similar retention across conditions (Reels 84.8%, neutral images 82.9%, rest 87.9%). No participant’s composite noise score exceeded the exclusion threshold (*z* > 2.00). For the sensitivity analyses, the two participants with the highest scores (intervention *z* = 1.93; task *z* = 1.85 and 1.75) were removed to form the cleaned sample (*N* = 28). Neither had completed the Reels-neutral pair, so the cleaned sample still included 10 participants who completed both Reels and neutral images and 9 participants for each of the other two pairs.

### Planned contrasts

Of the four planned contrasts, only C-T1 was significant after Holm correction, and only in the cleaned sample (Table 2). During the interventions, frontal-midline theta did not differ between Reels and neutral images (C-I1; absolute +0.014 ln(µV²), 95% CI [−0.177, 0.205], *p* = .881), and occipital alpha was numerically lower during Reels without reaching significance (C-I2; −0.221 [−0.584, 0.143], *p* = .226, Holm *p* = .903), with similar values in the cleaned sample (all *p* ≥ .30). In the 10 participants who completed both Reels and neutral images, both differences were also present (frontal-midline theta +0.126, paired *t* test *p* = .045; occipital alpha −0.225, Wilcoxon signed-rank test *p* = .027), but these uncorrected tests were not the planned inference.

**Table 2.** Planned (Two-Sided) Contrasts: Estimated Differences, 95% CI, Uncorrected p, and Holm-Corrected p.

| **Contrast** | **Phase / context** | **Measure** | **Full sample (N = 30)** | **Cleaned sample (N = 28)** |
|---|---|---|---|---|
| C-I1: frontal-midline theta, E − N | Intervention | Absolute | +0.014 [−0.177, 0.205], *p* = .881, Holm 1.00 | +0.014 [−0.185, 0.212], *p* = .890, Holm 1.00 |
| C-I1 | Intervention | Relative | +0.004 [−0.021, 0.029], *p* = .744, Holm 1.00 | +0.008 [−0.016, 0.032], *p* = .499, Holm 1.00 |
| C-I2: occipital alpha, E − N | Intervention | Absolute | −0.221 [−0.584, 0.143], *p* = .226, Holm .903 | −0.193 [−0.567, 0.181], *p* = .301, Holm 1.00 |
| C-I2 | Intervention | Relative | −0.027 [−0.094, 0.039], *p* = .413, Holm 1.00 | −0.024 [−0.090, 0.042], *p* = .474, Holm 1.00 |
| C-T1: occipital alpha, E − R | N-back | Absolute | +0.097 [0.015, 0.179], *p* = .022, Holm .087 | **+0.115 [0.030, 0.201], *p* = .010, Holm .039** |
| C-T1 | SART | Absolute | +0.001 [−0.076, 0.077], *p* = .985, Holm 1.00 | +0.030 [−0.031, 0.092], *p* = .321, Holm .726 |
| C-T1 | Full | Absolute | +0.065 [−0.004, 0.135], *p* = .064, Holm .257 | **+0.089 [0.022, 0.156], *p* = .011, Holm .046** |
| C-T1 | N-back; SART; Full | Relative | +0.013, *p* = .099; +0.001, *p* = .861; +0.009, *p* = .093 | +0.014, *p* = .113; +0.000, *p* = .963; +0.010, *p* = .099 |
| C-T2: frontal-midline theta, E − N | N-back; SART; Full | Absolute | −0.065, *p* = .396; −0.035, *p* = .659; −0.066, *p* = .357 | −0.076, *p* = .332; −0.080, *p* = .242; −0.077, *p* = .287 |
| C-T2 | N-back; SART; Full | Relative | −0.004, *p* = .366; −0.009, *p* = .118; −0.006, *p* = .167 | −0.005, *p* = .271; −0.009, *p* = .083; −0.007, *p* = .097 |

*Note.* E = Instagram Reels; N = neutral images; R = eyes-closed rest. Absolute power in ln(μV²); relative power as a fraction of 1-45 Hz power. Bold: Holm *p* < .05. Rows with three entries list the N-back, SART, and Full contexts; all of these had Holm *p* ≥ .29.

During the tasks, occipital absolute alpha was higher after Reels than after rest in the cleaned sample during the N-back (+0.115 [0.030, 0.201], *t*(26.1) = 2.79, *p* = .010, Holm *p* = .039) and the Full context (+0.089 [0.022, 0.156], *t*(26.1) = 2.72, *p* = .011, Holm *p* = .046), with the same direction in the full sample (N-back *p* = .022, Holm *p* = .087; Full *p* = .064). The effect withstood adjustment for task order and leave-one-out checks (worst *p* ≤ .035) but was absent in the SART (*p* ≥ .32) and for relative alpha (*p* ≥ .09). Neutral images differed from rest by a similar amount (see below), so C-T1 reflects a difference between viewing and rest rather than an effect specific to Reels. C-T2 was not significant (all *p* ≥ .08).

### Behavior and subjective fatigue

The N-back load manipulation was effective: *d*′ fell from 3.95 at 0-back to 1.77 at 3-back (*F*(3, 354.0) = 327.28, *p* < .001), and accuracy from 99.1% to 83.8% (*F*(3, 354.0) = 255.67, *p* < .001). No N-back index showed a condition effect (Fig. 2a): working-memory *d*′, *F*(2, 54.0) = 2.35, *p* = .105; working-memory accuracy, *F*(2, 54.0) = 1.88, *p* = .163; median hit RT, *F*(2, 54.0) = 0.25, *p* = .782; load slopes of *d*′ and RT, *p* ≥ .44 (all *p*_FDR_ ≥ .41). Working-memory *d*′ improved more after Reels than after neutral images (+0.25, 95% CI [0.02, 0.49], *p* = .036, *d* = 0.69), but this nominal difference did not survive Holm correction (*p* = .108). Neither Reels nor neutral images differed from rest (all *p* ≥ .18), and the secondary indices showed no condition effect (all *p* ≥ .18).

**Figure 2.**
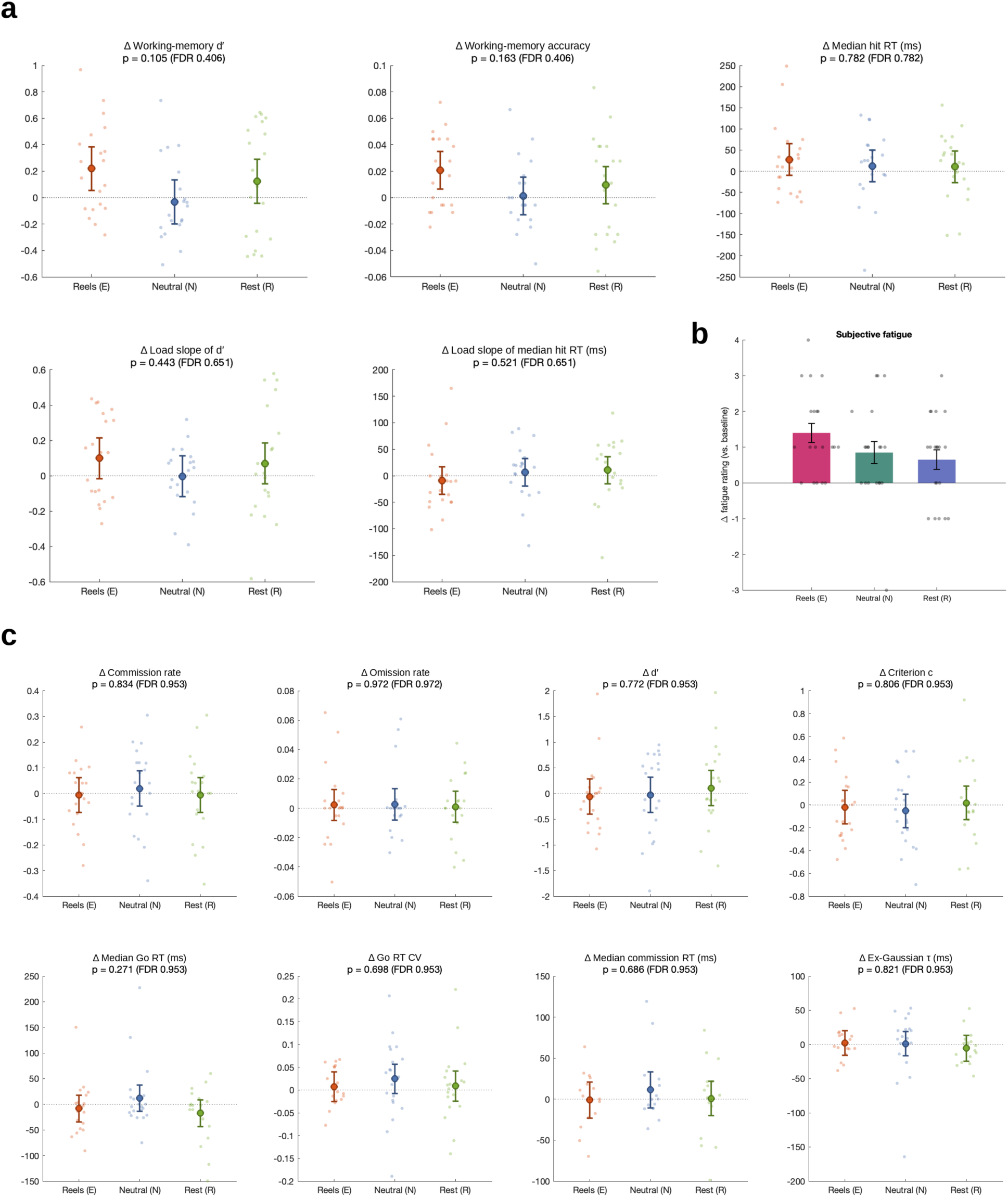
Behavioral results. (a) N-back: adjusted change from the pre-intervention block in working-memory *d*′ and accuracy (1-to 3-back), median hit RT, and the load slopes of *d*′ and median hit RT after Reels (E), neutral images (N), and rest (R); large markers show model-adjusted means with 95% CI, small dots individual change scores, and panel titles the omnibus p (FDR-corrected in parentheses); *N* = 30. (b) Subjective fatigue: change from baseline (mean ± SE; dots, individual participants). (c) SART: as in (a), for commission and omission rates, *d*′, criterion c, median Go RT, Go RT CV, median commission RT, and ex-Gaussian τ; *N* = 30 (RT-based indices, *n* = 23-29). Fatigue was higher after Reels than after rest (Holm *p* = .049); no N-back or SART omnibus test was significant.

No SART index differed by condition (Fig. 2c; Go RT indices, *n* = 29; commission RT, *n*= 23; ex-Gaussian parameters, *n* = 27). The omnibus tests for the eight primary indices had *p* values from .271 to .972 (all *p*_FDR_ ≥ .95; Supplementary Table S1), and every Reels contrast had *p* ≥ .27. Among the secondary indices, traditional post-error slowing showed a nominal omnibus effect (*F*(2, 25.4) = 3.68, *p* = .040, *p*_FDR_ = .317) that reflected a larger increase after neutral images than after rest (+98.8 ms, Holm *p* = .036); Reels did not differ from either condition (*p* ≥ .09). Intervention position and task order had no effect on any primary N-back or SART index (all *p* ≥ .09).

Fatigue on the 0-9 scale increased from baseline by 1.40 points after Reels, 0.85 after neutral images, and 0.65 after rest (omnibus *F*(2, 37.6) = 3.46, *p* = .042; Fig. 2b). Fatigue was higher after Reels than after rest (+0.80 [0.16, 1.45], *F*(1, 37.8) = 6.33, *p* = .016, Holm *p* = .049, BF_10_ ≈ 4.1) and, without reaching significance, than after neutral images (+0.61 [−0.04, 1.25], *p* = .064, Holm *p* = .127); neutral images and rest did not differ (*p* = .547). Fatigue was also higher after the second than after the first intervention (*b* = 0.60, *SE* = 0.24, *F*(1, 29.8) = 6.23, *p* = .018).

### EEG during the interventions

Watching either Reels or neutral images differed strongly from eyes-closed rest, and against this background Reels differed from neutral images in two spatially specific patterns (Table 3; all parameter families are listed in Supplementary Table S2, and forest plots and topographies are shown in Supplementary Figs. S1-S3).

**Table 3.** Intervention-EEG Condition Effects by Evidence Tier.

| **Pattern** | **Direction** | **Full sample (N = 30)** | **Cleaned sample (N = 28)** | **Evidence tier** |
|---|---|---|---|---|
| Viewing vs rest (alpha, delta, beta, gamma, ratios, IAF, SEF95, aperiodic) | Alpha: E, N < R; delta, beta, gamma, SEF95: E, N > R | All 62 electrodes for alpha; global Holm *p* < .001 | Same | Robust (manipulation check) |
| Posterior relative theta | E > N ≥ R | Occipital +0.037, Holm *p* < .001; 14 electrodes | Occipital +0.038, Holm *p* < .001; 14 electrodes | Robust |
| Posterior ln(α/θ) | E < N | Parietal and occipital, Holm *p* = .018; 19 electrodes | *p* = .025 and .019; 18 electrodes | Robust |
| Left centro-parietal alpha (relative, absolute, individualized) | E < N | 5-10 electrodes; central region *p* = .048; 10 of 10 within-person pairs | C3, CP3 (1-4 electrodes) | Robust core; extent sample-specific |
| Aperiodic exponent; SEF95 | Flatter; higher in E | 6; 20 electrodes | 3; 12 electrodes | Robust core |
| ln(α/β); ln engagement index | E < N; E > N | 20; 5 electrodes | 7; 1 electrodes | Sample-specific |
| Power above 30 Hz | E > N | Global Holm *p* = .046 | *p* = .116 | Exploratory; absent after artifact exclusion |
| C-I1 frontal-midline theta; C-I2 occipital alpha | - | *p* = .881; *p* = .226 | *p* = .890; *p* = .301 | Not supported (within-person *p* ≤ .049) |

*Note.* IAF = individual alpha frequency; SEF95 = 95% spectral edge frequency. Electrode counts are spatially dependent and should be compared with the 5% expected by chance.

Both viewing conditions showed the spectral profile of eyes-open visual processing, which is treated as a manipulation check because rest was recorded with the eyes closed. Global absolute alpha was 1.40 (Reels) and 1.19 (neutral images) ln units lower than at rest (E − R 95% CI [−1.68, −1.12]; omnibus *F*(2, 31.6) = 59.86, *p* < .001), relative power shifted toward delta, beta, and gamma, the individual alpha frequency (IAF) was about 0.67 Hz lower, the 95% spectral edge frequency (SEF95) was 7-9 Hz higher, and the aperiodic exponent was flatter (all Holm *p* ≤ .004). On every global measure Reels deviated further from rest than neutral images did, but the global E − N differences were not significant (for example, absolute alpha −0.208, *p* = .142).

The most consistent difference between the two viewing conditions was higher relative theta over parietal and occipital cortex during Reels (Fig. 3). Occipital relative theta was higher by 0.037 of total power (95% CI [0.020, 0.055], *t*(31.1) = 4.35, Holm *p* < .001; cleaned sample +0.038, Holm *p* < .001), and parietal relative theta by 0.030 ([0.006, 0.054], Holm *p* = .022; cleaned sample +0.033, Holm *p* = .009). The difference survived FDR and Holm correction at 14 parietal, parieto-occipital, and occipital electrodes in each sample, was reproduced by median-based estimates (13-14 electrodes), and was present in 9 of the 10 participants who completed both Reels and neutral images. In the within-person difference spectrum it appeared as an increase in the theta range together with a decrease in the alpha range (Fig. 4). The effect was specific to relative power and to posterior sites (occipital absolute theta, *p* = .197; frontal-midline and central relative theta, *p* ≥ .93), and posterior relative theta followed the ordering E > *N* ≥ R. Accordingly, the primary ratio index, ln(α/θ), was lower during Reels over parietal (−0.334 [−0.607, −0.060]) and occipital cortex (−0.362 [−0.660, −0.064]; both Holm *p* = .018; cleaned sample *p* = .025 and .019) and at 19 (full) and 18 (cleaned) centro-parietal to parieto-occipital electrodes after correction. Raw ratios, which varied far more between recordings during rest than during viewing, reproduced the viewing-rest differences but not this E − N difference (Supplementary Note).

**Figure 3.**
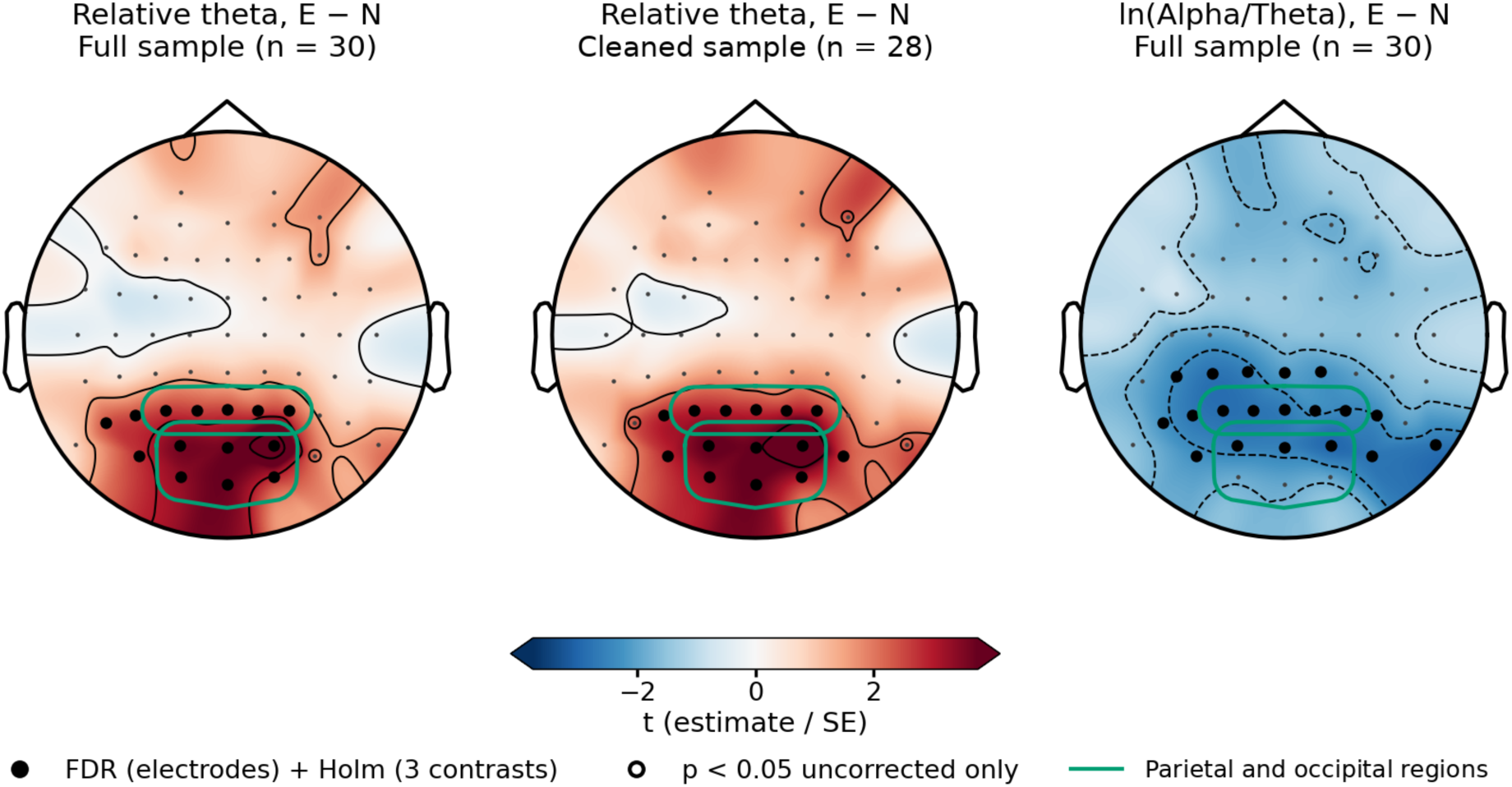
Reels − neutral differences in posterior theta during the interventions. Topographic maps of the electrode-wise *t* statistic (estimate/SE) for the E − N contrast in relative theta (4-8 Hz) in the full sample (left; *N* = 30) and the cleaned sample (middle; *N* = 28), and in ln(α/θ) in the full sample (right). Warm colours indicate higher values during Reels. Filled circles mark electrodes surviving FDR correction across electrodes and Holm correction across the three contrasts; open circles mark uncorrected *p* < .05 only. Green outlines show the parietal and occipital regions.

**Figure 4.**
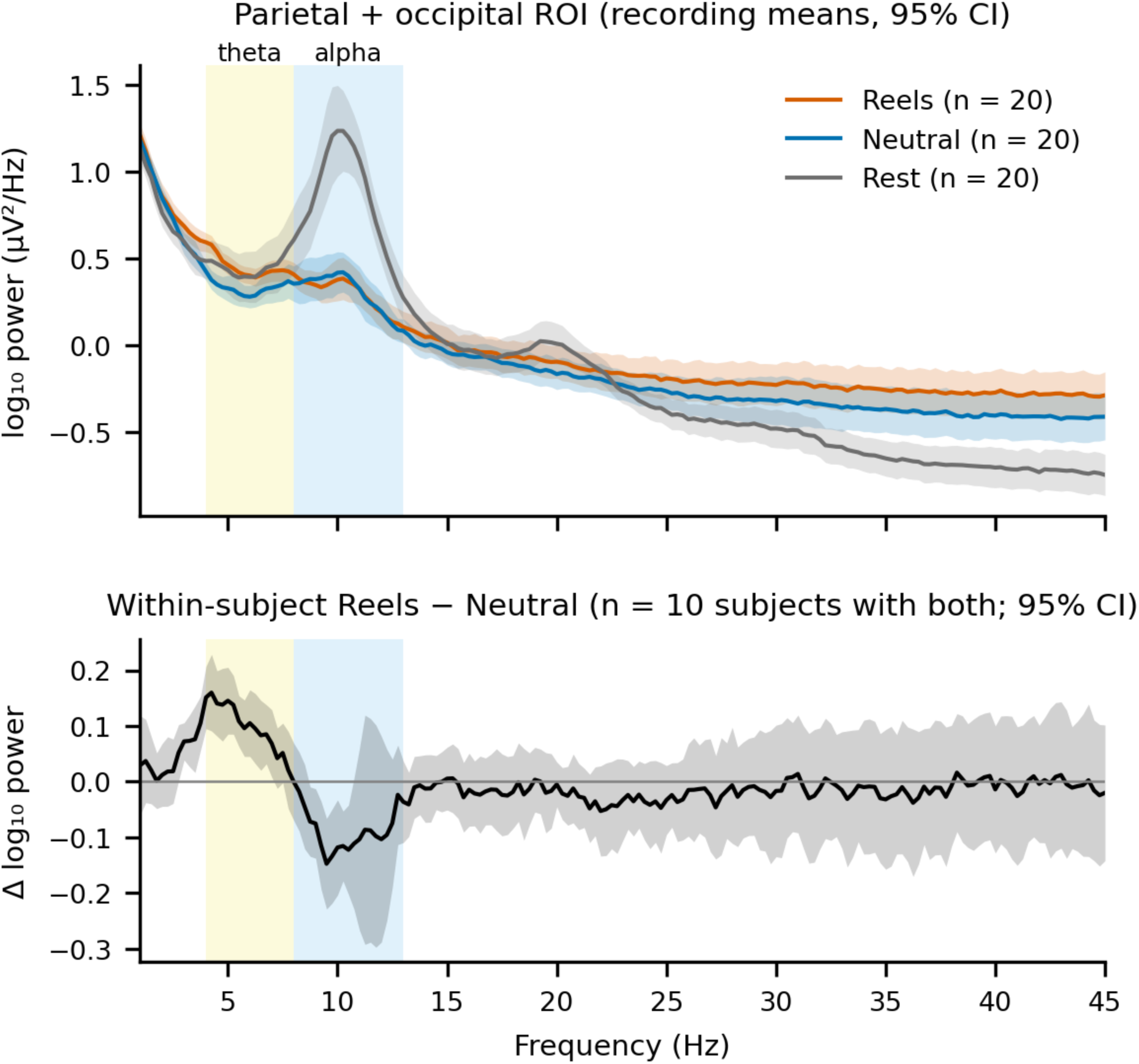
Posterior power spectra during the interventions. Upper panel: parietal-occipital log₁₀ power spectra (recording means, 95% CI) during Reels, neutral images, and rest (*n* = 20 recordings each), with the theta (4-8 Hz) and alpha (8-13 Hz) bands shaded; rest shows the eyes-closed alpha peak. Lower panel: within-person Reels − neutral difference spectrum (*n* = 10; 95% CI).

Alpha power over left centro-parietal cortex was lower during Reels. Relative alpha was lower at 10 electrodes after correction in the full sample (C5, C3, C1, CP5, CP3, CP1, CPz, CP2, P3, P1; −0.050 to −0.070) and at C3 and CP3 in the cleaned sample, and whole-scalp relative alpha was lower during Reels in all 10 participants who completed both Reels and neutral images (−0.037, paired *t* test *p* = .011), although the global mixed-model contrast was not significant (−0.039, *p* = .135). The same sites showed a flatter aperiodic exponent and a higher SEF95. The core of this pattern (C3, CP3) was robust, whereas its spatial extent was sample-specific.

Power above 30 Hz was higher during Reels than during neutral images at 17-24 electrodes per band in the full sample and globally for relative 30-45 Hz power (+0.023, Holm *p* = .046). This difference was absent in the cleaned sample (*p* = .116), in the 10 participants who completed both conditions (*p* = .52), and after three recordings flagged for high-frequency artifact were excluded (+0.010, *p* = .243), whereas the viewing-rest differences were unchanged; it is therefore treated as exploratory.

### EEG during the post-intervention tasks

During the tasks, condition effects converged on two patterns (Supplementary Tables S3 and S4; Supplementary Figs. S4-S19): a frontal alpha difference between Reels and neutral images, and a difference in posterior absolute power that both viewing conditions shared relative to rest. Both were clearest during the N-back, recurred in the Full context, and were weak or absent during the SART. Spectral-shape measures, aperiodic parameters, alpha asymmetry, and individualized alpha power showed no condition effect in any context.

During the N-back, relative alpha over frontal electrodes was higher after Reels than after neutral images (Fig. 5). In median-based models, the difference survived FDR and Holm correction at AF4 (+0.017 [0.004, 0.031], Holm *p* = .032), F6 (+0.018 [0.006, 0.030], Holm *p* = .009), and the frontal region (+0.017, Holm *p* = .044), and AF4 and F6 remained significant in the cleaned sample (Holm *p* = .024 and .014). In the primary mean-based electrode model, the difference was nominal at 7 (full) and 8 (cleaned) frontal electrodes, against 3.1 expected by chance, and positive at 60-61 of 62 electrodes. A model with recording half showed the same frontal distribution (Supplementary Fig. S20) and no change between the halves of the task.

**Figure 5.**
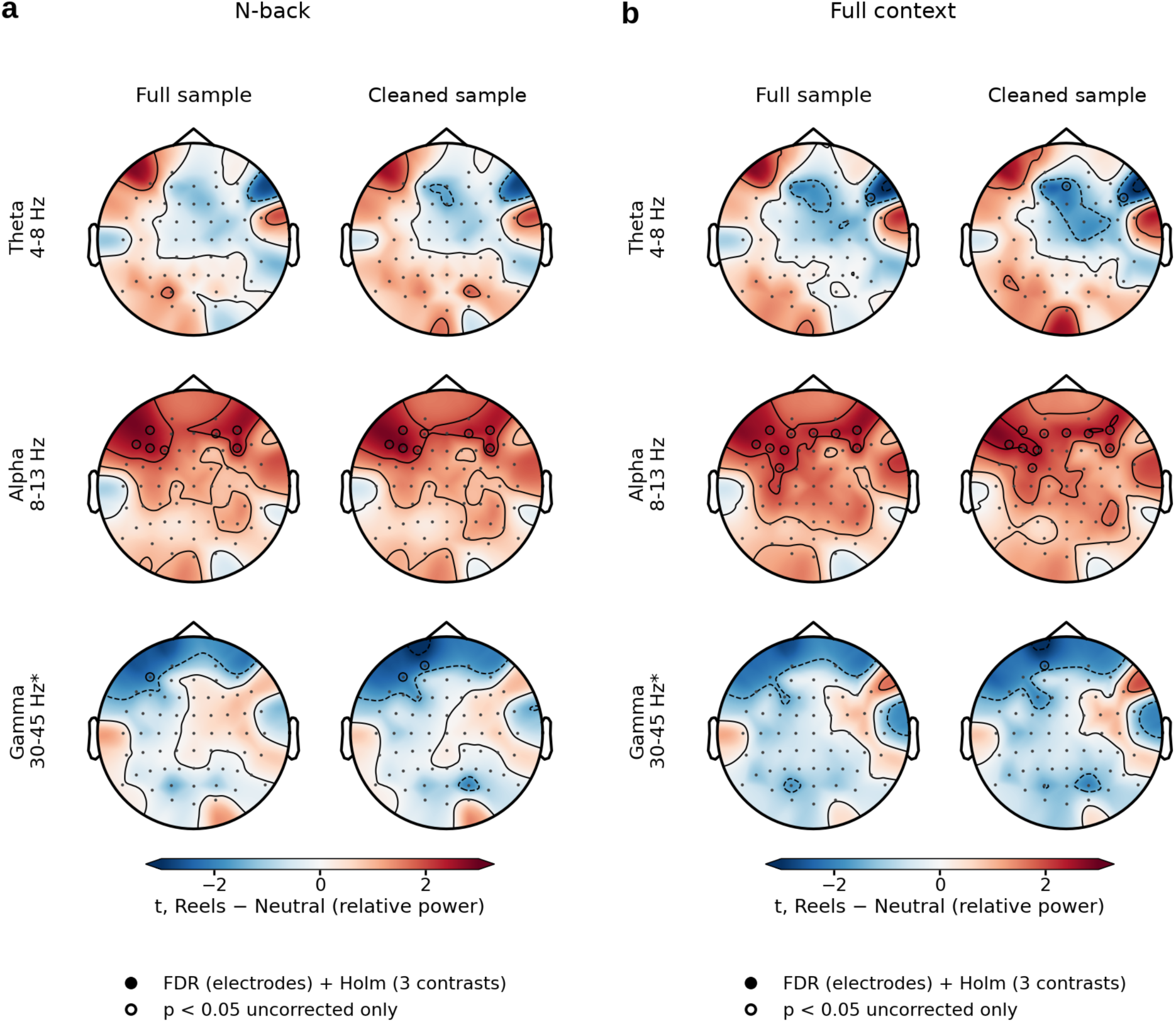
Reels − neutral differences in relative power during the post-intervention tasks. Electrode-wise *t* statistics for the E − N contrast in relative theta, alpha, and exploratory 30-45 Hz power during the N-back (a) and the Full context (b), each shown for the full sample and the cleaned sample. Warm colours indicate higher relative power after Reels; open circles mark uncorrected *p* < .05 (no electrode survived correction in this mean-based model). The frontal cluster of higher relative alpha after Reels is present in both contexts and both samples.

Neutral images lay close to rest, giving the ordering E > *N* ≥ R. Two related measures pointed the same way. The alpha/theta ratio was higher after Reels than after rest (global median-based ratio +0.243, Holm *p* = .021; cleaned sample +0.273, Holm *p* = .014), and the variability of relative alpha across windows was lower after Reels than after neutral images over central (−0.039 [−0.063, −0.015], Holm *p* = .007) and parietal regions (−0.033, Holm *p* = .012) in both samples (Supplementary Fig. S21). In the Full context, the frontal relative-alpha difference recurred at a nominal level at the same electrodes, and in the SART only a directional tendency remained.

Both viewing conditions also differed from rest in the same direction and by similar amounts. In the cleaned sample, absolute alpha and beta over occipital, parietal, and temporal cortex were higher, and occipital relative theta was lower, after Reels and after neutral images than after rest (N-back: occipital alpha E − R +0.126, N − R +0.133, both Holm *p* = .012; occipital relative theta E − R −0.0069, Holm *p* = .036, N − R −0.0102, Holm *p* = .003), with the same pattern in the Full context and no significant Reels-neutral contrast in these regions. In the full sample these effects were nominal (N-back, 7 of 90 regional tests against 4.5 expected). The C-T1 effect belongs to this shared pattern. Finally, independently of condition, N-back relative alpha was lower in the second than in the first post-intervention block (global −0.0148 [−0.0261, −0.0035], Holm *p* = .013; cleaned sample −0.0154, Holm *p* = .016; Supplementary Fig. S22), with no condition × position interaction.

## Discussion

Ten minutes of Instagram Reels changed EEG activity both during viewing and during the working-memory task that followed, but not task performance. During viewing, Reels shifted the posterior spectrum toward theta relative to neutral images and reduced alpha over a small left centro-parietal region; the two planned intervention contrasts were not significant. During the subsequent N-back, frontal relative alpha was higher and less variable after Reels than after neutral images. The only significant planned task contrast, higher occipital alpha after Reels than after rest, held in the cleaned sample only and was shared by neutral images. Fatigue rose more after Reels than after rest. Because no participant-level association between these measures was tested, the links drawn below are descriptive.

The most reliable difference between Reels and neutral images was a shift of posterior activity toward theta. It survived correction at 14 electrodes in both samples and was present in 9 of the 10 participants who completed both Reels and neutral images. It was specific to relative power, and the within-person difference spectrum showed a theta increase together with an alpha decrease, so it is best described as a change in the balance between the two bands. Neutral images moved posterior relative theta only slightly away from rest, whereas Reels moved it clearly, so the shift is something Reels added to the general state of looking at a screen. Rising task demands typically increase theta and decrease alpha [34], and lower posterior alpha is usually taken to indicate more processing of incoming visual input [31,32], so a greater visual and attentional load during Reels is a plausible reading. Alternatively, the shift may reflect stimulus properties: Reels contain motion, frequent cuts, and faces, whereas the neutral images were static and changed every 5 s. Increased theta has also been observed during mind wandering and low alertness [28], although the accompanying alpha decrease is more often associated with engagement with external input. The present design cannot separate engagement from stimulus dynamics, so the posterior theta-alpha shift is best treated as a marker of Reels viewing whose cognitive meaning remains to be tested.

The lower alpha over left centro-parietal cortex during Reels was found at 10 electrodes in the full sample but only at C3 and CP3 in the cleaned sample, so only these two electrodes showed the effect in both samples. Its location fits the sensorimotor mu rhythm, which is suppressed when people perform or observe movements [42]; all participants were right-handed and scrolled through Reels by hand, whereas the neutral images required no manual action. The same sites, however, showed a flatter aperiodic exponent and a higher spectral edge frequency, which is also the signature of muscle activity [36,37]. Both explanations point to the hand movements needed for scrolling rather than to a cognitive effect, which future studies could confirm with electromyography.

During the N-back, three measures pointed to stronger and steadier alpha after Reels: (i) relative alpha over frontal electrodes was higher after Reels than after neutral images, with neutral images close to rest (E > *N* ≥ R); (ii) the alpha/theta ratio was higher after Reels than after rest and, although not significantly, also higher than after neutral images; and (iii) the variability of relative alpha across windows was lower after Reels than after neutral images over central and parietal cortex. The frontal alpha difference had the same topography in both samples and did not change between the two halves of the task. It survived correction at AF4, F6, and the frontal region in median-based models, whereas in the primary mean-based model it was nominal at 7-8 frontal electrodes, about twice the number expected by chance, and positive at almost all electrodes. The difference was weak during the SART, and N-back performance did not differ between conditions.

An increase in alpha power can mean several things. In the classical view, alpha is an idling rhythm that grows when a region is less engaged, so higher alpha would indicate reduced cortical activation and disengagement from the task [34]. Alpha also increases with mental fatigue and declining alertness [43,44], and when attention drifts away from the task toward unrelated thoughts, as in mind wandering [28]. A more recent account treats alpha as active functional inhibition that serves the task: alpha increases in regions whose input would interfere with ongoing internal processing, and thereby protects it [31,33]. In line with this account, alpha is higher when attention is directed to internal rather than external material [45], increases with the number of items held in working memory [46], rises in anticipation of predictable distracters during working-memory maintenance, with larger increases predicting better performance [47], and increases over prefrontal cortex when the contents of working memory must be manipulated rather than merely retained [48].

Of these interpretations, active inhibition fits the present pattern best. Reduced engagement would be expected to impair performance, yet accuracy and response times were unchanged, and the difference was stable across the task rather than growing over time. A fatigue account is weakened by two observations: relative alpha during the N-back fell from the first to the second post-intervention block while fatigue rose, and frontal alpha after neutral images, after which fatigue was intermediate, lay close to that after rest. Mind wandering would be expected to increase errors or response-time variability, which did not occur, although a small cost could have gone undetected because the lower load levels were close to ceiling. In contrast, an inhibitory account best explains not only why alpha was higher while performance was preserved, but also why the difference appeared mainly in the N-back and was weak in the SART. The two tasks place opposite demands on the balance between internal and external processing. In the N-back, each digit is shown for only 500 ms, and the task is performed mainly on internal representations: the last one to three digits must be held in mind and updated on every trial, and each new digit must be compared with these stored items. Once a digit has been encoded, further visual input is irrelevant and can interfere with the stored items, so suppressing it through alpha-mediated inhibition helps the task. In the SART, by contrast, all task-relevant information is in the external stimulus stream: each digit must be perceived and evaluated as it appears, and a response must be withheld whenever the single No-Go digit is shown, but nothing has to be held across trials. Suppressing visual input would therefore not protect SART performance but impair it, so the mechanism that raised alpha during the N-back would not be expected to operate during the SART. For the same reason, a shift toward stronger shielding of internal representations after Reels would leave SART performance unaffected, which is consistent with the absence of any SART difference in either EEG or behavior. The lower window-to-window variability of alpha also fits a sustained rather than intermittent engagement of this mechanism during the N-back. On this reading, after a feed that repeatedly captures attention, participants may have relied more on top-down control to shield working-memory contents from external input, so that performance was maintained at a neural cost that did not appear in accuracy or speed [49].

The large differences between both viewing conditions (Reels and neutral images) and eyes-closed rest during the interventions, mainly much higher alpha during rest, reproduce the known physiology of opening the eyes [29,30] and say little about Reels as such. In the tasks that followed, the relation reversed: posterior absolute alpha was higher, and posterior relative theta lower, after both viewing conditions than after rest. Because neutral images differed from rest by a similar amount, C-T1 should not be read as an effect specific to Reels; rather, 10 min with the eyes closed seems to have changed the state in which participants began the next task. Three explanations are possible. First, participants may have started the task more alert after resting, and higher arousal is accompanied by lower alpha [30]. Second, after 10 min without visual input, opening the eyes and starting a visual task may have suppressed posterior alpha more strongly than usual; even short periods of visual deprivation increase the excitability of the visual cortex [50], although this has been shown after longer periods than those used here. Third, the reverse may have occurred after the viewing conditions: after 10 min of looking at a phone screen, participants may have begun the task with some visual or mental fatigue, which is accompanied by higher alpha [43,44]. The present design cannot distinguish between these explanations, and future studies that include an eyes-open rest condition and measures of arousal, such as skin conductance or pupil size, should examine them directly.

No N-back or SART index differed by condition. Several explanations for the gap between EEG and performance are possible. Performance can be protected by additional effort, so that the cost of a demanding state appears in subjective experience or physiology before it appears in accuracy or speed [49], and models that treat mental fatigue as a signal about the costs and benefits of continuing a task explain why higher fatigue need not be accompanied by poorer performance in short tasks [51,52]. Statistical sensitivity is a second explanation. Because the design included two reference conditions, each condition effect was tested in three pairwise contrasts, and correcting for these contrasts and for the number of measures set a stricter threshold than a two-condition comparison would; in addition, only 10 participants completed both interventions of each pair, the 0- and 1-back levels were close to ceiling, and widely used tasks can be insensitive to small changes [26]. Third, the tasks started at different delays after the intervention, so a brief effect could have faded before part of the testing took place. Fatigue after Reels did not differ significantly from fatigue after neutral images, so the data do not show whether the fatigue effect is specific to short-form video or reflects screen viewing in general; in a previous study, both smartphone use and a documentary increased mental fatigue [12].

The SART showed neither a performance nor an EEG difference between conditions, whereas two earlier studies found poorer inhibition after social media or smartphone use: fewer correct No-Go rejections after 15 min of scrolling an Instagram feed [13] and more No-Go errors after 45 min of free smartphone use [12]. Besides the different demands of the two tasks described above, several differences between these studies and the present one may account for this. First, both earlier studies compared separate groups of participants before and after exposure [12,13], whereas here each participant was compared with themselves. In a between-group design, stable differences between the groups can contribute to the result; in the Instagram study, for example, the social media group used Instagram for 7.8 h a week and the control group for 2.2 h, and 80% and 30% of them, respectively, met a criterion for social media addiction [13]. Second, the exposures differed: 45 min of free smartphone use, mostly on social networks [12], and 15 min of a mixed feed of Instagram posts and videos [13], compared with 10 min of Reels here. Third, the inhibition tasks of those studies contained more No-Go trials, 30 [12] and 42 [13], than the 25 No-Go trials of each SART block here, and the SART, which lasted about 4 min per block, also provided fewer EEG windows than the N-back. Fourth, the participants here were master’s students with moderate Instagram use and low psychological distress, tested in a laboratory while wearing an EEG cap, a group and a setting in which performance may be maintained more easily after a short break. Finally, findings for inhibition are themselves mixed: 30 min of social media use did not affect Go/No-Go performance in adolescents [22].

The absence of a performance effect agrees with experiments that found no immediate change after a short social media session [21,22] and differs from studies that found impaired vigilance and inhibition after smartphone use [12], poorer inhibitory control after Instagram scrolling [13], or impaired prospective memory after short videos [14,15]. These studies differed from the present one in the length and content of the exposure, in their mostly between-group designs, and in their outcomes; prospective memory, for example, was not assessed here. The neutral-image condition was close to the nature-image feed used by Aitken et al. [13], yet performance after Reels did not differ from that after neutral images. Frontal-midline theta, which has been linked to a short-video addiction tendency [38], did not differ between conditions here. The correlational associations of habitual short-form video use with attention and inhibition [3] describe differences between people that build up over months or years, which a single 10-min session would not be expected to reproduce. Exposure duration may also matter. The 10-min session lies at the short end of the range tested so far: costs to vigilance and inhibition were reported after 15 and 45 min [12,13], whereas 6 min of social media [21] and, in adolescents, 30 min [22] left working memory and inhibition unchanged, and effects on prospective memory appeared after only 10 min of short videos [14,15]. Duration alone therefore does not explain the differences between studies, but 10 min of Reels may have been enough to change brain activity without producing a measurable behavioral cost, which may require longer or repeated exposure.

The study has several limitations. The sample was small, no a priori power analysis was carried out, and only 10 participants completed both interventions of each pair. Only one of the four planned contrasts was significant, and only in the cleaned sample, so most EEG findings are exploratory and require replication. The four regions used for the planned contrasts were defined after the electrode-level results had been inspected, and several task-EEG effects reached correction only in median-based models. No electromyography was recorded, rest was recorded with the eyes closed, Reels were watched without sound on participants’ own phones and their content was not recorded, and the exposure lasted only 10 min. The fatigue scale could not record a rating of 0, and four participants’ ratings were corrected from the experimenter’s notes. Finally, all participants were right-handed master’s students in Tehran who used Instagram for 1-3 h a day, and the findings may not generalize to other ages, heavier users, or other cultures [53]. The study also has strengths: two reference conditions that separate effects of screen viewing from those specific to Reels, counterbalanced within-person comparisons, continuous EEG during exposure and testing, and parallel analyses in two samples.

A pre-registered replication should target the posterior theta-alpha shift during viewing and the frontal alpha difference during the N-back, ideally in a design in which every participant completes all three conditions, with a dynamic, non-social control matched to the cut rate of Reels, electromyography, several exposure durations, testing at fixed delays, and participant-level analyses that link EEG change to fatigue and performance. In sum, a single 10-min session of Instagram Reels was accompanied by a posterior shift toward theta during viewing and by higher frontal alpha during a subsequent working-memory task, and it left participants more fatigued than eyes-closed rest, without measurably changing sustained attention, response inhibition, or working memory. These EEG differences are candidates for confirmatory studies rather than evidence that brief Reels viewing impairs cognition.

## Methods

### Participants and design

The study used a within-subject, counterbalanced design (Fig. 1a). Each participant completed three testing blocks separated by two 10-min interventions and received two of the three intervention conditions: Instagram Reels (E), neutral images (N), or eyes-closed rest (R) (Fig. 1b). Each intervention was therefore completed by 20 participants, and 10 participants completed both interventions of each pair. The order of the interventions was counterbalanced (Supplementary Table S5).

Thirty master’s students in Tehran took part (15 women and 15 men; age *M* = 23.9 years, *SD* = 1.47, range = 21-26 years). All were right-handed. Data were collected between December 2025 and January 2026. Participants were recruited through a call for volunteers posted on university messaging channels, notice boards, and student social networks. To be eligible, participants had to be university students aged 18-26 years with an active Instagram account who reported using Instagram for 1-3 h a day over the previous month. Requiring an active account meant that participants watched Reels in a familiar interface. Participants also needed normal or corrected-to-normal vision.

Exclusion criteria included a self-reported history of psychiatric or cognitive disorders; a score of 20 or higher on the Kessler Psychological Distress Scale (K10) [54]; a history of seizures or other neurological disease; use of recreational drugs or stimulants in the preceding month; regular use of medication affecting the central nervous system; and participation in a similar study within the preceding six months. Before testing, a psychiatrist screened each participant for relevant medical conditions, including epilepsy and other neurological disorders, head injury, and the use of medication affecting the central nervous system. Testing sessions were not conducted, or their data were excluded, if participants failed to follow the experimental protocol, had experienced a severely stressful event within the preceding 24 h, had engaged in strenuous exercise the night before, had slept fewer than six hours, had consumed caffeine or nicotine within 8 h before testing, or had undertaken cognitively demanding activities within the preceding 2 h. All participants scored below the K10 exclusion threshold, and no sessions were excluded or rescheduled because of violations of the pre-session requirements.

The sample size of 20 participants per condition was chosen to match comparable short-exposure experiments, which included 15-40 participants per condition [12,14,15,17,20].

### Ethics declarations

The study was approved by the Research Ethics Committee of the University of Social Welfare and Rehabilitation Sciences, Tehran, Iran (approval code IR.USWR.REC.1404.203). All methods were performed in accordance with the relevant guidelines and regulations, including the Declaration of Helsinki. Written informed consent was obtained from all participants, and participants received monetary compensation for taking part. Before participation, participants were informed that the study involved cognitive testing, but the specific research aims were not disclosed until the end of the session. They were assured that their data would remain confidential and be stored using identification codes rather than names, and that they could withdraw from the study at any time.

### Cognitive tasks and fatigue rating

Sustained attention and response inhibition were measured with the SART [40] (Fig. 1c, left), programmed in MATLAB with Psychtoolbox [55]. The digits 1-9 appeared one at a time. Participants pressed the spacebar for every digit except 3, the No-Go target, for which they had to withhold the response. Each digit appeared 25 times, giving 225 trials in random order, 25 of which were No-Go trials. Each digit was shown for 250 ms and followed by a 900-ms mask, for a trial length of 1150 ms. Responses were accepted during both periods, and reaction time (RT) was measured from digit onset. Digit size varied randomly across five levels.

Working memory was measured with a visual N-back task [41], adapted from the version used by Aitken et al. [13], with the digits 0-9 at four load levels (Fig. 1c, right). In the 0-back condition, participants pressed the spacebar whenever the digit 0 appeared; in the 1-, 2-, and 3-back conditions, they pressed it whenever the current digit matched the one shown one, two, or three positions earlier. Each time it was administered, the N-back task comprised 240 trials in 12 blocks of 20 trials: the four load levels were presented in ascending order (0-, 1-, 2-, then 3-back), and this sequence was repeated three times. Between 26% and 30% of trials were targets. Each digit was shown for 500 ms and followed by a 1500-ms blank interval; responses were recorded in both periods and timed from digit onset.

At the end of every testing block, participants rated how tired they felt on a scale from 0 (least fatigue) to 9 (most fatigue), illustrated with faces, by pressing the corresponding number key.

### Interventions

Each intervention lasted 10 min. In the Reels condition (E), participants watched Reels from their own feeds on their own smartphones and Instagram accounts, scrolling as they normally would. The videos were not limited to a predefined set, so the condition reflected each participant’s everyday viewing rather than a standard set of clips. Sound was off, as in a previous Instagram experiment that presented no audio in either condition to avoid auditory confounds [13]; this also kept auditory input identical across the two viewing conditions, because the neutral images were silent. In the neutral-image condition (N), participants watched on their smartphones a silent video of neutral nature photographs, such as grey pebbles, in which each photograph was shown for 5 s. Their neutrality was checked in a pilot study in which 10 volunteers, who did not take part in the main experiment, viewed the video and were asked whether any photograph evoked emotions or personal memories. This condition matched the Reels condition for holding a phone and looking at its screen, but it contained no social content, no motion, and no scrolling; a comparable control, a feed of black- and-white nature images, was used in the same study [13]. In the rest condition (R), participants sat with their eyes closed, with no task or stimulus, and this condition served as a resting reference for both viewing conditions. The duration of 10 min was chosen to resemble a short break spent on social media and lies within the range of earlier exposure experiments, which used 9 to 45 min [12–14,18].

### Procedure

Data were collected at the National Brain Mapping Laboratory (NBML), Tehran, Iran, in a quiet room with the same lighting and temperature for all sessions. Participants had been asked to sleep at least six hours the night before, to avoid caffeine and nicotine for 8 h and smartphone use for 1 h before the session, and to avoid cognitively demanding work in the two hours before it. After the EEG cap was fitted, a baseline resting-state recording was made (2 min with eyes closed, then 2 min with eyes open). Participants then completed the first testing block, the first intervention, the second testing block, the second intervention, and the third testing block, and the session ended with a second resting-state recording (Fig. 1; Supplementary Table S6). These two resting-state recordings were not analysed in the present study. There were no breaks other than the interventions, and EEG was recorded continuously. Within each testing block, the two tasks ran in one of two orders, SART then N-back (SN) or N-back then SART (NS), and the order was planned separately for each block. The fatigue rating always came at the end of the block.

### EEG recording and preprocessing

EEG was recorded with a g.tec system at 512 Hz from 65 Ag/AgCl electrodes: 62 recording electrodes placed according to the extended 10-20 system, two earlobe electrodes (A1 and A2) used as reference, and a ground electrode at Fpz. The same setup was used throughout the session. Event triggers were sent through the parallel port at the start and end of each task and at every digit onset. In the SART they also marked mask onsets and responses. In the N-back, the codes identified the load level of each block, target and non-target digits, hits and false alarms at the moment of the response, and misses and correct rejections at the end of each trial. Trial outcomes for the behavioral analyses were scored from the task log files.

EEG data were preprocessed in MATLAB with EEGLAB using a semi-automatic pipeline adapted from the Harvard Automated Processing Pipeline for Electroencephalography (HAPPE) [56], in which all steps were automated and their output, including the flagged channels and independent components, was checked by visual inspection. The signals were re-referenced to the average of A1 and A2 and band-pass filtered between 1 and 100 Hz with a finite impulse response (FIR) filter, which retained the frequency range analysed here while removing slow drifts and high-frequency noise; a notch filter at 47-53 Hz was applied to remove 50 Hz line noise. The first and last 5 s of each recording were removed. The HAPPE workflow was followed closely, except that its wavelet-enhanced ICA step was replaced by ICLabel classification with visual inspection. Noisy channels were first flagged automatically from their spectra (1-45 Hz, normalized threshold ±3) and then confirmed or added by visual inspection. Extended independent component analysis (ICA) was performed on a 1-45 Hz filtered copy of the data. Independent components were classified with ICLabel, and components classified as eye artifacts with a probability of at least 0.80, or as muscle, cardiac, line-noise, or channel-noise artifacts with a probability of at least 0.90, were removed from the original 1-100 Hz data after visual inspection. Removed channels were reconstructed by spherical interpolation.

The preprocessed recordings were segmented into 4-s windows (2048 samples) with 50% overlap. A window was rejected if it contained a non-finite value, an absolute amplitude above 150 µV on any channel, or a channel with a standard deviation below 0.01 µV; in the intervention recordings, windows containing a boundary that marked removed data were also rejected. All remaining windows of each recording were used, and a power spectrum was computed for each with a 2048-point FFT (frequency resolution 0.25 Hz). A composite noise score was computed for each participant, separately for the intervention recordings and for the N-back and SART recordings, as the mean of standardized quality indicators: the percentage of rejected windows, the ratio of high-frequency to total power, and the absolute standardized deviation of log broadband power, together with, for the intervention recordings, the number of quality problems noted in the preprocessing log (available for 53 of 60 recordings). Participants with a composite score above the threshold of *z* > 2.00 were to be excluded.

### EEG measures

Spectral measures were computed for the intervention recordings and for three task contexts after each intervention: the N-back, the SART, and the Full context, in which the N-back and SART windows were pooled. Absolute band power was expressed as ln(µV²), and relative band power was calculated as a fraction of total power in the 1-45 Hz range. The frequency bands were delta (1-4 Hz), theta (4-8 Hz), alpha (8-13 Hz), and beta (13-30 Hz), with exploratory subdivisions of beta (13-20 and 20-30 Hz) and gamma-frequency bands (30-45, 55-80, and 80-100 Hz). Exploratory band-ratio measures included theta/beta, alpha/theta, alpha/beta, and the engagement index, defined as β/(α + θ) [57]. These ratios were computed from absolute band power using both mean-based and median-based estimates. For the intervention recordings, the natural logarithm of the mean-based ratio (e.g., ln(α/θ)) was also computed and served as the primary ratio index, because raw ratios varied far more between recordings during rest than during viewing; the three variants were corrected in separate FDR families, and the task analyses used the mean- and median-based ratios.

Additional measures included hemispheric alpha asymmetry, calculated as the difference between log-transformed right- and left-hemisphere alpha power; individual alpha frequency (IAF), defined for each channel as the peak frequency within 8-13 Hz of its mean power spectrum; individualized alpha power, the power in the band IAF ± 2 Hz of each channel; the exponent and offset of the aperiodic component of the power spectrum [58]; and the 95% spectral edge frequency (SEF95), defined as the frequency below which 95% of the cumulative power between 1 and 45 Hz was contained. Recording-level mean values were obtained by aggregating across valid windows. Median-based power estimates were also computed as exploratory robustness measures. The coefficient of variation (CV) of band power was calculated as the standard deviation divided by the mean across valid windows, providing an index of temporal variability in spectral power.

Measures were analysed at individual electrodes, across the whole scalp, and within five predefined regions of interest: frontal (23 electrodes), central (7), temporal (6), parietal (16), and occipital (8). The frontal region included AFz, Fp1, Fp2, AF3, AF4, F7, F3, Fz, F4, F8, FC5, FC1, FC2, FC6, AF7, AF8, F5, F1, F2, F6, FC3, FCz, and FC4. The central region included C5, C3, C1, Cz, C2, C4, and C6; the temporal region included T7, T8, FT7, FT8, TP7, and TP8; the parietal region included CP5, CP1, CP2, CP6, P7, P3, Pz, P4, P8, CP3, CPz, CP4, P5, P1, P2, and P6; and the occipital region included PO3, PO4, O1, O2, PO7, POz, PO8, and Oz. P9 and P10 were excluded from regional averages but retained in whole-scalp analyses. The planned contrasts and the regional band-power results during the interventions used four literature-based regions instead: frontal midline (Fz, FCz, F1, F2, FC1, FC2), central (C3, C1, Cz, C2, C4), parietal (P3, P1, Pz, P2, P4), and occipital (PO3, POz, PO4, O1, Oz, O2).

### Behavioral measures

For the N-back task, the primary indices were working-memory sensitivity (*d*′) and accuracy averaged over the 1-to 3-back levels, the median RT of hits at these levels, and the slopes of *d*′ and of median hit RT across the four load levels; the secondary indices were *d*′, accuracy, and median hit RT across all load levels, the response criterion *c*, the CV of hit RT, and the load slope of accuracy. Sensitivity was computed with the log-linear correction for extreme proportions [59]. For the SART, the primary indices were the commission rate (the percentage of No-Go trials with a response, where higher values mean poorer inhibition), the omission rate (the percentage of Go trials without a response), *d*′, the response criterion *c*, the median Go RT, the CV of Go RT, the median RT of commission errors, and the τ parameter of an ex-Gaussian fit to Go RTs, which captures the slow tail of the distribution. The secondary SART indices were the mean and SD of Go RT, the ex-Gaussian μ and σ, the difference between median Go RT and median commission RT, pre-error speeding, and post-error slowing computed in the traditional and the robust way. Because each block had only 25 No-Go trials, one commission error changes the commission rate by 4 percentage points.

Trials were scored as follows. In the SART, responses recorded less than 100 ms after digit onset were attributed to the preceding trial when that trial had no response of its own and were otherwise treated as anticipatory, in which case the trial was excluded. A SART block was excluded if more than 20% of presses were anticipatory, if more than 30% of trials were unusable, or if *d*′ was 0 or below. RT indices were not computed for a block when more than 10% of its responses had no measurable RT; commission RT required at least three commission errors, and the ex-Gaussian fit at least 50 correct Go RTs. In the N-back, a block was excluded if 0-back accuracy was below 80%, and RTs below 100 ms were not used in the RT indices.

### Statistical analysis

#### EEG

Each EEG measure was analysed with a linear mixed model, Metric ∼ Condition + (1 | Subject), fitted by restricted maximum likelihood with Satterthwaite degrees of freedom, separately for the intervention recordings and for each task context. Omnibus tests were corrected with the false discovery rate (FDR) procedure [60] within families, and pairwise contrasts (E − N, E − R, and N − R) were computed for units that survived FDR correction and were Holm-corrected across the three contrasts [61]. FDR families were formed separately for each measure (absolute and relative power were never pooled). At the electrode and region levels, each frequency band formed its own family across the 62 electrodes or across the regions of interest; for the global values, alpha was tested alone, and delta, theta, and beta, the two beta subdivisions, and the three gamma-frequency bands formed separate families. Each band ratio and each of its computation variants formed a separate family. No correction was applied across task contexts, measure variants, or the two samples. A positive contrast means a higher value in the first-named condition. The four planned contrasts were tested two-sided, with Holm correction across four tests (two hypotheses × absolute and relative power) in the intervention phase and in each task context. All analyses were run with equal weight in the full sample (*N* = 30) and in a cleaned sample that excluded the two participants with the highest noise scores (*N* = 28). Findings were graded as planned (planned contrasts surviving Holm correction), robust (surviving FDR and Holm correction in both samples), sample-specific (in one sample only), nominal (uncorrected *p* < .05), or near-threshold (.05 ≤ *p* < .10). Because neighbouring electrodes are not independent, the number of significant electrodes was compared with the 5% expected by chance rather than treated as a count of independent effects.

Several checks supported the main models. Median-based estimates were analysed in the same way. For the 10 participants who received both Reels and neutral images, paired *t* tests or Wilcoxon signed-rank tests were run as descriptive within-person checks without correction. Models with condition × recording half entered two observations per recording without modelling their dependence, which inflates their degrees of freedom (about 80-110, against about 30 elsewhere), so they were treated as supportive only. Forest plots of these follow-up models are shown in Supplementary Figs. S23-S25. The C-T1 effect was also checked with adjustment for task order and with leave-one-out analyses.

#### Behavior

N-back and SART indices were analysed as change from the block immediately preceding each intervention, in linear mixed models with condition, intervention position (first or second), and task order in the pre- and post-intervention blocks as predictors, no separate intercept, and a random intercept for each participant (change ∼ −1 + E + N + R + position + order_pre + order_post + (1 | participant)); the models were fitted with fitlme in MATLAB with Satterthwaite degrees of freedom. Omnibus F tests were FDR-corrected separately within the primary and within the secondary indices of each task, pairwise contrasts were Holm-corrected, and effects are reported as model-estimated differences with 95% confidence intervals and Cohen’s d (the model-estimated contrast divided by the square root of the sum of the random-intercept and residual variances). The effect of load in the N-back was tested in a separate model of raw scores with load and block (baseline, first and second post-intervention block) as fixed effects and a random intercept for each participant.

#### Fatigue

Fatigue ratings after the two interventions were analysed with a linear mixed model in MATLAB (fitlme, maximum likelihood) that included condition, the participant’s baseline rating (centered), and intervention position, with a random intercept for each participant. The omnibus effect of condition and the three pairwise contrasts were tested with Satterthwaite degrees of freedom [62], and contrasts were Holm-corrected. For the Reels-rest contrast, a Bayes factor was computed from the model estimate and its standard error, with a normal prior on the effect centered on zero and a standard deviation of 0.5 baseline standard deviations [63]; BF_10_ values above 3 were taken as moderate evidence for an effect. Statistical significance was set at *p* < .05, two-tailed, throughout.

### Use of generative AI

Generative AI (Claude, Anthropic) was used to assist with proofreading, language editing, improving manuscript readability, and drafting and revising selected passages under the authors’ direction. All AI-assisted content was critically reviewed and verified by the authors, who take full responsibility for the accuracy, integrity, and scientific content of the manuscript.

## Data availability

The processed spectral and behavioral data are available from Zahra Ramezani or the corresponding author upon reasonable request. Raw EEG data are not publicly available to protect participant privacy; requests for these data will be considered in accordance with participant consent and the study’s ethics approval.

## Code availability

The analysis code is available at https://github.com/Aminem-Mohammadi/Reels-EEG-Whole-Block-Spectral-Power.

## Acknowledgements

We thank the National Brain Mapping Laboratory (NBML), Tehran, Iran, for providing data acquisition services.

## Author contributions

Z.R. and A.M.M. conceived and designed the study, collected the data, performed the analyses, and prepared the figures. A.M.M. wrote the task programs and the analysis code. Z.R. wrote the first draft of the manuscript, and A.M.M. reviewed and edited it. Z.H. and B.D. supervised the study. All authors reviewed and approved the final manuscript.

## Additional information

Supplementary information accompanies this paper. Correspondence and requests for materials should be addressed to B.D.

