## Supplementary Information for "Brain rhythms during Instagram Reels viewing and subsequent performance on sustained attention and working memory: an EEG study"

### Watching Instagram Reels changes brain rhythms without impairing attention or working memory: an EEG study

### Contents

Supplementary Table S1. Behavioral condition contrasts

Supplementary Table S2. Intervention-EEG parameter inventory

Supplementary Table S3. Task-EEG parameter inventory

Supplementary Table S4. Task-EEG omnibus condition tests by family, level, and measure

Supplementary Table S5. Assignment of participants to interventions and task orders

Supplementary Table S6. Session timeline

Supplementary Note. Choice of the log-transformed alpha/theta ratio

Supplementary Figs. S1-S3. Intervention EEG: global and regional differences and topographies

Supplementary Figs. S4-S19. Task EEG: global and regional differences, within-person values, spectra, and SART topographies

Supplementary Figs. S20-S22. N-back relative alpha: recording-half model, variability, and change between blocks

Supplementary Figs. S23-S25. Follow-up models for the Full context, N-back, and SART

Abbreviations used throughout: E, Instagram Reels; N, neutral images; R, eyes-closed rest. Full sample, N = 30; cleaned sample, N = 28 (the two participants with the highest EEG noise scores excluded). Full context, the N-back and SART recordings of a testing block analysed together.

**Supplementary Table S1.** Behavioral condition contrasts

Estimated differences in change scores (95% CI, uncorrected *p*; Holm *p* for contrasts with *p* < .10). Change scores are the post-intervention block minus the immediately preceding block; N-back N = 30; SART N = 30 for error rates, *d*′ and *c*, n = 29 for Go RT indices, n = 27 for ex-Gaussian parameters, n = 23 for commission RT, Go − commission RT, pre-error speeding and robust post-error slowing, and n = 22 for traditional post-error slowing. Working-memory (WM) indices average the 1- to 3-back levels; load slopes are linear slopes across the 0- to 3-back levels. Accuracy and error rates are in percentage points.

| Outcome | E − N | E − R | N − R |
|---|---|---|---|
| **Primary indices** | | | |
| N-back WM d′ | +0.25 [0.02, 0.49], p = .036 (Holm .108) | +0.10 [−0.14, 0.33], p = .416 | −0.16 [−0.39, 0.08], p = .186 |
| N-back WM accuracy (points) | +1.9 [−0.1, 4.0], p = .059 (Holm .177) | +1.1 [−0.9, 3.1], p = .265 | −0.8 [−2.8, 1.2], p = .418 |
| N-back WM median hit RT (ms) | +15.1 [−38.1, 68.2], p = .572 | +16.9 [−35.9, 69.6], p = .524 | +1.8 [−51.1, 54.8], p = .946 |
| N-back load slope of d′ (per level) | +0.103 [−0.062, 0.267], p = .216 | +0.030 [−0.134, 0.193], p = .716 | −0.073 [−0.237, 0.091], p = .377 |
| N-back load slope of median hit RT (ms per level) | −15.8 [−52.5, 20.9], p = .392 | −19.8 [−56.2, 16.6], p = .281 | −4.0 [−40.5, 32.6], p = .827 |
| SART commission errors (points) | −2.5 [−12.3, 7.2], p = .604 | 0.0 [−9.7, 9.7], p = .999 | +2.5 [−7.2, 12.2], p = .601 |
| SART omission errors (points) | −0.03 [−1.52, 1.45], p = .964 | +0.13 [−1.34, 1.60], p = .859 | +0.16 [−1.31, 1.64], p = .824 |
| SART d′ | −0.03 [−0.52, 0.46], p = .896 | −0.16 [−0.65, 0.32], p = .500 | −0.13 [−0.62, 0.35], p = .587 |
| SART criterion c | +0.032 [−0.178, 0.241], p = .761 | −0.037 [−0.244, 0.171], p = .726 | −0.069 [−0.277, 0.140], p = .513 |
| SART median Go RT (ms) | −20.0 [−56.6, 16.5], p = .276 | +9.0 [−27.8, 45.9], p = .625 | +29.1 [−7.5, 65.7], p = .117 |
| SART Go RT CV | −0.018 [−0.064, 0.029], p = .447 | −0.001 [−0.048, 0.045], p = .948 | +0.016 [−0.030, 0.062], p = .488 |
| SART median commission RT (ms) | −12.5 [−43.9, 18.9], p = .426 | −1.9 [−32.6, 28.8], p = .900 | +10.6 [−20.0, 41.1], p = .488 |
| SART ex-Gaussian τ (ms) | +1.1 [−24.5, 26.7], p = .931 | +7.7 [−18.7, 34.0], p = .562 | +6.5 [−19.6, 32.7], p = .616 |
| Fatigue (0-9) | +0.61 [−0.04, 1.25], p = .064 (Holm .127) | +0.80 [0.16, 1.45], p = .016 (Holm .049) | +0.20 [−0.46, 0.85], p = .547 |
| **Secondary indices** | | | |
| N-back d′, all loads | +0.23 [−0.04, 0.49], p = .091 (Holm .274) | +0.06 [−0.20, 0.33], p = .632 | −0.16 [−0.43, 0.10], p = .218 |
| N-back accuracy, all loads (points) | +1.4 [−0.1, 3.0], p = .066 (Holm .198) | +0.8 [−0.7, 2.3], p = .302 | −0.6 [−2.2, 0.9], p = .402 |
| N-back median hit RT, all loads (ms) | +15.0 [−11.0, 41.0], p = .251 | +20.5 [−5.3, 46.3], p = .117 | +5.4 [−20.4, 31.3], p = .675 |
| N-back criterion c | +0.050 [−0.093, 0.194], p = .484 | −0.036 [−0.178, 0.106], p = .615 | −0.086 [−0.229, 0.057], p = .231 |
| N-back hit RT CV | −0.015 [−0.089, 0.059], p = .689 | −0.001 [−0.075, 0.073], p = .977 | +0.014 [−0.060, 0.088], p = .708 |
| N-back load slope of accuracy (points per level) | +1.10 [−0.26, 2.47], p = .112 | +0.46 [−0.90, 1.82], p = .499 | −0.64 [−2.00, 0.72], p = .349 |
| SART mean Go RT (ms) | −17.5 [−51.9, 17.0], p = .314 | +4.1 [−30.7, 38.9], p = .814 | +21.6 [−13.0, 56.1], p = .216 |
| SART SD of Go RT (ms) | −10.7 [−30.6, 9.3], p = .285 | −0.2 [−20.4, 20.0], p = .981 | +10.4 [−9.6, 30.5], p = .299 |
| SART ex-Gaussian μ (ms) | −21.9 [−70.8, 27.0], p = .372 | +6.0 [−44.3, 56.4], p = .811 | +27.9 [−22.0, 77.8], p = .266 |
| SART ex-Gaussian σ (ms) | −2.5 [−20.9, 15.9], p = .785 | +5.1 [−13.8, 24.0], p = .589 | +7.6 [−11.1, 26.4], p = .418 |
| SART Go − commission RT (ms) | −7.5 [−45.4, 30.3], p = .690 | +11.4 [−25.6, 48.4], p = .537 | +18.9 [−17.9, 55.7], p = .305 |
| SART pre-error speeding (ms) | −35.8 [−82.0, 10.4], p = .125 | −39.1 [−84.7, 6.5], p = .091 (Holm .272) | −3.3 [−48.8, 42.2], p = .884 |
| SART post-error slowing, traditional (ms) | −62.7 [−136.3, 10.8], p = .091 (Holm .182) | +36.1 [−36.2, 108.4], p = .313 | +98.8 [23.4, 174.3], p = .012 (Holm .036) |
| SART post-error slowing, robust (ms) | −63.0 [−171.9, 45.9], p = .249 | −9.7 [−118.0, 98.7], p = .858 | +53.3 [−55.1, 161.8], p = .326 |

**Supplementary Table S2.** Intervention-EEG parameter inventory: FDR-surviving omnibus condition tests and Holm-surviving E − N contrasts (full sample / cleaned sample)

| Analysis family | Parameters | Spatial levels | Tests per sample | Omnibus, FDR | E − N, FDR + Holm |
|---|---|---|---|---|---|
| Global band power | Log absolute and relative power, 9 bands | Whole scalp | 18 | 12 / 12 | 1 / 0 |
| Regional band power | As above | 5 regions | 90 | 67 / 68 | 7 / 4 |
| Literature-defined regions (incl. C-I1, C-I2) | As above | 4 regions | 72 | 51 / 53 | 11 / 7 |
| Electrode band power | As above | 62 electrodes | 1116 | 747 / 698 | 130 / 72 |
| Median-based robust estimates | Median absolute and relative power; CV of absolute and relative power; total power | Electrode, region | 2584 | 1253 / 1112 | 104 / 73 |
| Band-ratio indices | Theta/beta, alpha/theta, alpha/beta, engagement index; ln-mean (primary), mean- and median-based | Global, region, electrode | 816 | 630 / 614 | 48 / 33 |
| Alpha asymmetry | ln(right alpha) − ln(left alpha) | 5 electrode pairs, regions, global | 22 | 0 / 0 | - |
| Spectral shape | IAF, SEF95 | Electrode, region, global | 136 | 102 / 101 | 25 / 14 |
| Aperiodic (1/f) component | Exponent and offset; FOOOF and robust fit | Electrode, region, global | 272 | 238 / 231 | 7 / 3 |
| Individualized alpha | Log absolute and relative power in IAF ± 2 Hz | Electrode | 124 | 124 / 124 | 14 / 5 |
| Condition × recording half | Condition main effect | Electrode, global | 1134 | 872 / 854 | 395 / 310 |
| Condition × recording half | Time-half main effect; interaction | Electrode, global | 1134 each | 0 / 0; 0 / 0 | - |
| Condition × intervention position | Condition; position; interaction | Electrode, global | 1134 each | 655 / 680; 0 / 0; 0 / 0 | 120 / 78 |

Counts include exploratory beta sub-bands and gamma bands; most surviving omnibus tests reflect viewing-rest differences. Counts from the recording-half model are inflated by pseudo-replication. A cross-participant sign-consistency analysis was run as a descriptive diagnostic and is reported separately.

**Supplementary Table S3.** Task-EEG parameter inventory and FDR-surviving omnibus condition tests (full sample / cleaned sample)

| Analysis family | Parameters | Spatial levels | Tests per sample | N-back | SART | Full |
|---|---|---|---|---|---|---|
| Global band power | Log absolute and relative power, 9 bands | Whole scalp | 18 | 0 / 0 | 0 / 0 | 0 / 0 |
| Regional band power | As above | 5 regions (frontal, central, temporal, parietal, occipital) | 90 | 0 / 11 | 0 / 0 | 0 / 7 |
| Literature-defined regions (incl. C-T1, C-T2) | As above | 4 regions (frontal midline, central, parietal, occipital) | 72 | 0 / 5 | 0 / 0 | 0 / 2 |
| Electrode band power | As above | 62 electrodes | 1116 | 0 / 1 | 0 / 0 | 0 / 0 |
| Median-based robust estimates | Median absolute and relative power; CV of absolute and relative power; total power | Electrode, region | 2584 | 6 / 7 | 0 / 0 | 0 / 3 |
| Band-ratio indices | Theta/beta, alpha/theta, alpha/beta, engagement index; mean- and median-based | Global, region, electrode | 544 | 1 / 4 | 0 / 0 | 1 / 4 |
| Alpha asymmetry | ln(right alpha) − ln(left alpha) | 5 electrode pairs, regions, global | 22 | 0 / 0 | 0 / 0 | 0 / 0 |
| Spectral shape | IAF, SEF95 | Electrode, region, global | 136 | 0 | 0 | 0 |
| Aperiodic (1/f) component | Exponent and offset; FOOOF and robust fit | Electrode, region, global | 272 | 0 | 0 | 0 |
| Individualized alpha | Log absolute and relative power in IAF ± 2 Hz | Electrode | 124 | 0 / 0 | 0 / 0 | 0 / 0 |
| Condition × recording half | Condition main effect | Electrode, global | 1134 | 42 / 46 | 1 / 7 | - |
| Condition × recording half | Time-half main effect; interaction | Electrode, global | 1134 each | 0 / 0 | 0 / 0 | - |
| Condition × intervention position | Condition; position; interaction | Electrode, global | 1134 each | 6 / 0; 34 / 34; 0 / 0 | 0 / 0; 0 / 2; 0 / 0 | - |
| Reconstructed Full (N-back + SART windows) | Band power | Global, region, electrode | 1224 | - | - | 0 / 10 |

Spectral-shape and aperiodic analyses were run on the full sample only. The task pipeline computed only mean- and median-based band ratios.

**Supplementary Table S4.** Task-EEG omnibus condition tests by family, level, and measure

Each cell gives FDR-surviving tests / tests in that family and level (nominally significant tests, uncorrected $p < .05$, in parentheses). FS = full sample (N = 30); CS = cleaned sample (N = 28). Spectral-shape and aperiodic analyses were run on the full sample only; the recording-half and position models were run for the N-back and SART only; the reconstructed Full analysis is shown in the Full columns. Expected nominal count under the null = 5% of tests.

| Family | Level; measure | N-back FS | N-back CS | SART FS | SART CS | Full FS | Full CS |
|---|---|---|---|---|---|---|---|
| Global band power | Global; absolute power | 0/9 (0) | 0/9 (0) | 0/9 (0) | 0/9 (0) | 0/9 (0) | 0/9 (0) |
| Global band power | Global; relative power | 0/9 (0) | 0/9 (0) | 0/9 (0) | 0/9 (0) | 0/9 (0) | 0/9 (0) |
| Regional band power (5 regions) | Region; absolute power | 0/45 (6) | 10/45 (13) | 0/45 (0) | 0/45 (1) | 0/45 (0) | 7/45 (9) |
| Regional band power (5 regions) | Region; relative power | 0/45 (1) | 1/45 (4) | 0/45 (0) | 0/45 (0) | 0/45 (1) | 0/45 (4) |
| Literature-defined regions (4) | Region; absolute power | 0/36 (4) | 4/36 (7) | 0/36 (0) | 0/36 (0) | 0/36 (0) | 2/36 (5) |
| Literature-defined regions (4) | Region; relative power | 0/36 (1) | 1/36 (2) | 0/36 (0) | 0/36 (0) | 0/36 (1) | 0/36 (2) |
| Electrode band power | Electrode; absolute power | 0/558 (39) | 0/558 (62) | 0/558 (1) | 0/558 (10) | 0/558 (10) | 0/558 (42) |
| Electrode band power | Electrode; relative power | 0/558 (14) | 1/558 (27) | 0/558 (5) | 0/558 (7) | 0/558 (13) | 0/558 (19) |
| Median-based robust estimates | Electrode; median absolute | 0/558 (21) | 0/558 (38) | 0/558 (0) | 0/558 (7) | 0/558 (9) | 0/558 (22) |
| Median-based robust estimates | Electrode; CV absolute | 0/558 (26) | 0/558 (38) | 0/558 (4) | 0/558 (18) | 0/558 (7) | 0/558 (8) |
| Median-based robust estimates | Electrode; CV relative | 0/558 (47) | 0/558 (47) | 0/558 (6) | 0/558 (15) | 0/558 (5) | 0/558 (4) |
| Median-based robust estimates | Electrode; median relative | 3/558 (17) | 2/558 (27) | 0/558 (6) | 0/558 (10) | 0/558 (13) | 0/558 (19) |
| Median-based robust estimates | Electrode; total power | 0/124 (6) | 0/124 (20) | 0/124 (0) | 0/124 (1) | 0/124 (4) | 0/124 (7) |
| Median-based robust estimates | Global; median absolute | 0/9 (1) | 0/9 (0) | 0/9 (0) | 0/9 (0) | 0/9 (0) | 0/9 (1) |
| Median-based robust estimates | Global; CV absolute | 0/9 (0) | 0/9 (0) | 0/9 (0) | 0/9 (1) | 0/9 (0) | 0/9 (0) |
| Median-based robust estimates | Global; CV relative | 0/9 (0) | 0/9 (0) | 0/9 (0) | 0/9 (0) | 0/9 (0) | 0/9 (0) |
| Median-based robust estimates | Global; median relative | 0/9 (0) | 0/9 (0) | 0/9 (0) | 0/9 (0) | 0/9 (0) | 0/9 (0) |
| Median-based robust estimates | Global; total power | 0/2 (0) | 0/2 (0) | 0/2 (0) | 0/2 (0) | 0/2 (0) | 0/2 (0) |
| Median-based robust estimates | Region; median absolute | 0/45 (1) | 2/45 (7) | 0/45 (0) | 0/45 (0) | 0/45 (0) | 1/45 (6) |
| Median-based robust estimates | Region; CV absolute | 0/45 (2) | 0/45 (5) | 0/45 (0) | 0/45 (1) | 0/45 (1) | 0/45 (2) |
| Median-based robust estimates | Region; CV relative | 2/45 (5) | 2/45 (3) | 0/45 (0) | 0/45 (0) | 0/45 (0) | 0/45 (0) |
| Median-based robust estimates | Region; median relative | 1/45 (1) | 1/45 (4) | 0/45 (0) | 0/45 (0) | 0/45 (1) | 2/45 (4) |
| Median-based robust estimates | Region; total power | 0/10 (0) | 0/10 (2) | 0/10 (0) | 0/10 (0) | 0/10 (0) | 0/10 (2) |
| Band-ratio indices | Electrode; alpha/beta | 0/124 (4) | 0/124 (3) | 0/124 (0) | 0/124 (0) | 0/124 (0) | 0/124 (0) |
| Band-ratio indices | Electrode; alpha/theta | 0/124 (34) | 0/124 (44) | 0/124 (0) | 0/124 (6) | 0/124 (38) | 0/124 (50) |
| Band-ratio indices | Electrode; engagement index | 0/124 (0) | 0/124 (3) | 0/124 (0) | 0/124 (2) | 0/124 (0) | 0/124 (1) |
| Band-ratio indices | Electrode; theta/beta | 0/124 (3) | 0/124 (5) | 0/124 (2) | 0/124 (3) | 0/124 (1) | 0/124 (5) |
| Band-ratio indices | Global; alpha/beta | 0/2 (0) | 0/2 (0) | 0/2 (0) | 0/2 (0) | 0/2 (0) | 0/2 (0) |
| Band-ratio indices | Global; alpha/theta | 1/2 (1) | 2/2 (2) | 0/2 (0) | 0/2 (0) | 1/2 (1) | 2/2 (2) |
| Band-ratio indices | Global; engagement index | 0/2 (0) | 0/2 (0) | 0/2 (0) | 0/2 (0) | 0/2 (0) | 0/2 (0) |
| Band-ratio indices | Global; theta/beta | 0/2 (0) | 0/2 (0) | 0/2 (0) | 0/2 (0) | 0/2 (0) | 0/2 (0) |
| Band-ratio indices | Region; alpha/beta | 0/10 (0) | 0/10 (0) | 0/10 (0) | 0/10 (0) | 0/10 (0) | 0/10 (0) |
| Band-ratio indices | Region; alpha/theta | 0/10 (4) | 2/10 (4) | 0/10 (0) | 0/10 (0) | 0/10 (2) | 2/10 (5) |
| Band-ratio indices | Region; engagement index | 0/10 (0) | 0/10 (1) | 0/10 (0) | 0/10 (0) | 0/10 (0) | 0/10 (1) |
| Band-ratio indices | Region; theta/beta | 0/10 (0) | 0/10 (0) | 0/10 (0) | 0/10 (0) | 0/10 (0) | 0/10 (1) |

| Family | Level; measure | N-back FS | N-back CS | SART FS | SART CS | Full FS | Full CS |
|---|---|---|---|---|---|---|---|
| Alpha asymmetry | Global | 0/2 (0) | 0/2 (0) | 0/2 (0) | 0/2 (0) | 0/2 (0) | 0/2 (0) |
| Alpha asymmetry | Pair; frontal | 0/6 (0) | 0/6 (0) | 0/6 (0) | 0/6 (0) | 0/6 (0) | 0/6 (0) |
| Alpha asymmetry | Pair; posterior | 0/4 (0) | 0/4 (0) | 0/4 (0) | 0/4 (0) | 0/4 (0) | 0/4 (0) |
| Alpha asymmetry | Region (all regions) | 0/10 (0) | 0/10 (0) | 0/10 (0) | 0/10 (0) | 0/10 (0) | 0/10 (0) |
| Spectral shape | Electrode; IAF | 0/62 (0) | - | 0/62 (5) | - | 0/62 (1) | - |
| Spectral shape | Electrode; SEF95 | 0/62 (0) | - | 0/62 (0) | - | 0/62 (1) | - |
| Spectral shape | Global; IAF | 0/1 (0) | - | 0/1 (0) | - | 0/1 (0) | - |
| Spectral shape | Global; SEF95 | 0/1 (0) | - | 0/1 (0) | - | 0/1 (0) | - |
| Spectral shape | Region; IAF | 0/5 (0) | - | 0/5 (0) | - | 0/5 (0) | - |
| Spectral shape | Region; SEF95 | 0/5 (0) | - | 0/5 (0) | - | 0/5 (0) | - |
| Aperiodic component | Electrode; FOOOF exponent | 0/62 (0) | - | 0/62 (0) | - | 0/62 (2) | - |
| Aperiodic component | Electrode; FOOOF offset | 0/62 (0) | - | 0/62 (0) | - | 0/62 (0) | - |
| Aperiodic component | Electrode; robust-fit exponent | 0/62 (1) | - | 0/62 (0) | - | 0/62 (1) | - |
| Aperiodic component | Electrode; robust-fit offset | 0/62 (1) | - | 0/62 (0) | - | 0/62 (0) | - |
| Aperiodic component | Global; FOOOF exponent | 0/1 (0) | - | 0/1 (0) | - | 0/1 (0) | - |
| Aperiodic component | Global; FOOOF offset | 0/1 (0) | - | 0/1 (0) | - | 0/1 (0) | - |
| Aperiodic component | Global; robust-fit exponent | 0/1 (0) | - | 0/1 (0) | - | 0/1 (0) | - |
| Aperiodic component | Global; robust-fit offset | 0/1 (0) | - | 0/1 (0) | - | 0/1 (0) | - |
| Aperiodic component | Region; FOOOF exponent | 0/5 (0) | - | 0/5 (0) | - | 0/5 (0) | - |
| Aperiodic component | Region; FOOOF offset | 0/5 (0) | - | 0/5 (0) | - | 0/5 (0) | - |
| Aperiodic component | Region; robust-fit exponent | 0/5 (0) | - | 0/5 (0) | - | 0/5 (0) | - |
| Aperiodic component | Region; robust-fit offset | 0/5 (0) | - | 0/5 (0) | - | 0/5 (0) | - |
| Individualized alpha | Electrode; absolute | 0/62 (6) | 0/62 (9) | 0/62 (2) | 0/62 (1) | 0/62 (1) | 0/62 (9) |
| Individualized alpha | Electrode; relative | 0/62 (5) | 0/62 (8) | 0/62 (1) | 0/62 (0) | 0/62 (5) | 0/62 (4) |
| Condition × recording half | Condition; electrode; absolute | 24/558 (121) | 31/558 (135) | 0/558 (8) | 4/558 (18) | - | - |
| Condition × recording half | Condition; electrode; relative | 16/558 (54) | 13/558 (58) | 1/558 (26) | 3/558 (23) | - | - |
| Condition × recording half | Condition; global; absolute | 1/9 (3) | 1/9 (4) | 0/9 (0) | 0/9 (0) | - | - |
| Condition × recording half | Condition; global; relative | 1/9 (1) | 1/9 (1) | 0/9 (1) | 0/9 (0) | - | - |
| Condition × recording half | Half; electrode; absolute | 0/558 (0) | 0/558 (0) | 0/558 (0) | 0/558 (0) | - | - |
| Condition × recording half | Half; electrode; relative | 0/558 (6) | 0/558 (8) | 0/558 (0) | 0/558 (6) | - | - |
| Condition × recording half | Half; global; absolute | 0/9 (0) | 0/9 (0) | 0/9 (0) | 0/9 (0) | - | - |
| Condition × recording half | Half; global; relative | 0/9 (0) | 0/9 (0) | 0/9 (0) | 0/9 (0) | - | - |
| Condition × recording half | Interaction; electrode; absolute | 0/558 (0) | 0/558 (0) | 0/558 (0) | 0/558 (0) | - | - |
| Condition × recording half | Interaction; electrode; relative | 0/558 (0) | 0/558 (0) | 0/558 (1) | 0/558 (21) | - | - |
| Condition × recording half | Interaction; global; absolute | 0/9 (0) | 0/9 (0) | 0/9 (0) | 0/9 (0) | - | - |
| Condition × recording half | Interaction; global; relative | 0/9 (0) | 0/9 (0) | 0/9 (0) | 0/9 (1) | - | - |
| Condition × intervention position | Condition; electrode; absolute | 0/558 (9) | 0/558 (9) | 0/558 (6) | 0/558 (24) | - | - |
| Condition × intervention position | Condition; electrode; relative | 6/558 (61) | 0/558 (65) | 0/558 (24) | 0/558 (13) | - | - |
| Condition × intervention position | Condition; global; absolute | 0/9 (0) | 0/9 (0) | 0/9 (0) | 0/9 (1) | - | - |
| Condition × intervention position | Condition; global; relative | 0/9 (1) | 0/9 (1) | 0/9 (1) | 0/9 (0) | - | - |
| Condition × intervention position | Position; electrode; absolute | 0/558 (5) | 0/558 (5) | 0/558 (5) | 2/558 (31) | - | - |
| Condition × intervention position | Position; electrode; relative | 33/558 (55) | 33/558 (56) | 0/558 (2) | 0/558 (9) | - | - |
| Condition × intervention position | Position; global; absolute | 0/9 (0) | 0/9 (0) | 0/9 (0) | 0/9 (0) | - | - |
| Condition × intervention position | Position; global; relative | 1/9 (1) | 1/9 (1) | 0/9 (0) | 0/9 (0) | - | - |
| Condition × intervention position | Interaction; electrode; absolute | 0/558 (18) | 0/558 (18) | 0/558 (10) | 0/558 (25) | - | - |
| Condition × intervention position | Interaction; electrode; relative | 0/558 (49) | 0/558 (41) | 0/558 (12) | 0/558 (16) | - | - |
| Condition × intervention position | Interaction; global; absolute | 0/9 (0) | 0/9 (0) | 0/9 (0) | 0/9 (0) | - | - |
| Condition × intervention | Interaction; global; relative | 0/9 (0) | 0/9 (0) | 0/9 (0) | 0/9 (0) | - | - |

| Family | Level; measure | N-back FS | N-back CS | SART FS | SART CS | Full FS | Full CS |
|---|---|---|---|---|---|---|---|
| position | | | | | | | |
| Reconstructed Full | Electrode; absolute power | - | - | - | - | 0/558 (12) | 1/558 (52) |
| Reconstructed Full | Electrode; relative power | - | - | - | - | 0/558 (10) | 0/558 (23) |
| Reconstructed Full | Global; absolute power | - | - | - | - | 0/9 (0) | 0/9 (0) |
| Reconstructed Full | Global; relative power | - | - | - | - | 0/9 (0) | 0/9 (0) |
| Reconstructed Full | Region; absolute power | - | - | - | - | 0/45 (1) | 8/45 (11) |
| Reconstructed Full | Region; relative power | - | - | - | - | 0/45 (1) | 1/45 (5) |

**Supplementary Table S5.** Assignment of participants to interventions and task orders

| Participant | Gender | Block 1 | Intervention 1 | Block 2 | Intervention 2 | Block 3 |
|---|---|---|---|---|---|---|
| S1 | Female | NS | N | NS | R | SN |
| S2 | Female | NS | N | SN | R | NS |
| S3 | Male | NS | N | SN | R | SN |
| S4 | Female | SN | E | NS | R | SN |
| S5 | Female | SN | E | SN | N | NS |
| S6 | Female | NS | E | SN | N | NS |
| S7 | Female | NS | E | SN | R | NS |
| S8 | Female | SN | R | NS | N | SN |
| S9 | Male | NS | E | NS | R | SN |
| S10 | Male | SN | N | NS | R | NS |
| S11 | Female | SN | N | SN | E | NS |
| S12 | Male | SN | R | NS | N | NS |
| S13 | Female | NS | R | SN | N | NS |
| S14 | Female | SN | N | NS | E | SN |
| S15 | Female | SN | E | SN | R | NS |
| S16 | Male | SN | R | NS | E | NS |
| S17 | Male | SN | E | NS | R | NS |
| S18 | Female | NS | R | SN | E | NS |
| S19 | Male | SN | R | SN | N | NS |
| S20 | Male | NS | R | SN | E | SN |
| S21 | Female | NS | N | SN | E | NS |
| S22 | Male | NS | R | NS | E | SN |
| S23 | Male | SN | N | SN | R | NS |
| S24 | Female | NS | R | NS | N | SN |
| S25 | Male | NS | E | NS | N | SN |
| S26 | Male | NS | N | NS | E | SN |
| S27 | Male | SN | N | NS | E | NS |
| S28 | Male | SN | E | NS | N | NS |
| S29 | Female | SN | R | SN | E | NS |
| S30 | Male | NS | E | SN | N | SN |

Blocks 1-3, testing blocks, with the task order in each block (SN, SART then N-back; NS, N-back then SART). Interventions: E, Instagram Reels; N, neutral images; R, eyes-closed rest. Each pair of interventions was completed by 10 participants.

**Supplementary Table S6.** Session timeline

| Step | Component | Duration | Content | Use in the present analyses |
|---|---|---|---|---|
| 1 | Baseline resting state | 4 min | 2 min eyes closed, then 2 min eyes open | Not analysed |
| 2 | Testing block 1 | 12.3 min | SART (225 trials of 1.15 s; 4.3 min) and N-back (240 trials of 2 s in 12 blocks of 20; 8 min), in SART-N-back or N-back-SART order; fatigue rating at the end | Pre-intervention block for intervention 1 |
| 3 | Intervention 1 | 10 min | Instagram Reels (E), neutral images (N), or eyes-closed rest (R) | Intervention EEG |
| 4 | Testing block 2 | 12.3 min | As block 1 | Post-intervention block for intervention 1 and pre-intervention block for intervention 2 |
| 5 | Intervention 2 | 10 min | The second assigned condition | Intervention EEG |
| 6 | Testing block 3 | 12.3 min | As block 1 | Post-intervention block for intervention 2 |
| 7 | Final resting state | 4 min | 2 min eyes closed, then 2 min eyes open | Not analysed |

EEG was recorded continuously throughout the session (about 65 min). Durations of the testing blocks are nominal (number of trials × trial duration) and exclude the self-paced instruction screens shown before each task and each N-back block and the self-paced fatigue rating. There were no breaks other than the interventions. The order of the two tasks was planned separately for each testing block, and the order of the interventions was counterbalanced across participants (main text, Fig. 1).

### Supplementary Note. Choice of the log-transformed alpha/theta ratio

Band ratios were computed from the recording-level mean absolute power in each band, so that alpha/theta was the mean 8-13 Hz power divided by the mean 4-8 Hz power. The raw ratio was strongly heteroscedastic across conditions. During eyes-closed rest, the posterior alpha peak made the ratio both large and highly variable between recordings (parietal region, $M = 4.39$, $SD = 2.73$; occipital region, $M = 6.21$, $SD = 4.64$; 20 recordings), whereas during viewing it was close to 1 and varied much less (parietal $SD = 0.32$ for Reels and 0.50 for neutral images; occipital $SD = 0.30$ and 0.56). The between-recording standard deviation was therefore about 5 to 16 times larger during rest than during viewing. After log transformation, the spread was much more similar across conditions (parietal $SD = 0.30$, 0.37, and 0.68 for Reels, neutral images, and rest; occipital $SD = 0.27$, 0.34, and 0.75).

The mixed models estimate a single residual variance for all three conditions, so the large variance of the raw ratio during rest inflated the standard error of every contrast, including E − N ($SE = 0.50$ for the parietal and 0.85 for the occipital region). The raw ratios therefore reproduced the large viewing-rest differences (parietal E − R = −3.48 and N − R = −3.10; occipital E − R = −5.25 and N − R = −4.85; all Holm $p < .001$) but not the smaller E − N difference (parietal −0.37, $p = .457$; occipital −0.40, $p = .638$; median-based ratios, $p = .639$ and .797). The raw E − N difference was nevertheless in the same direction and of a similar relative size as the difference in the log-transformed ratio: the raw parietal ratio was about 30% lower during Reels than during neutral images (−0.37 relative to a neutral-image mean of 1.26), and the log-ratio difference of −0.334 corresponds to a 28% lower ratio.

For these reasons, $\ln(\alpha/\theta)$ was used as the primary ratio index. It was lower during Reels than during neutral images over parietal (−0.334, 95% CI [−0.607, −0.060]) and occipital cortex (−0.362, [−0.660, −0.064]; both Holm $p = .018$). The pattern was the same in the cleaned sample (raw ratio, parietal and occipital E − N $p = .489$ and .634; log-transformed ratio, $p = .025$ and .019).

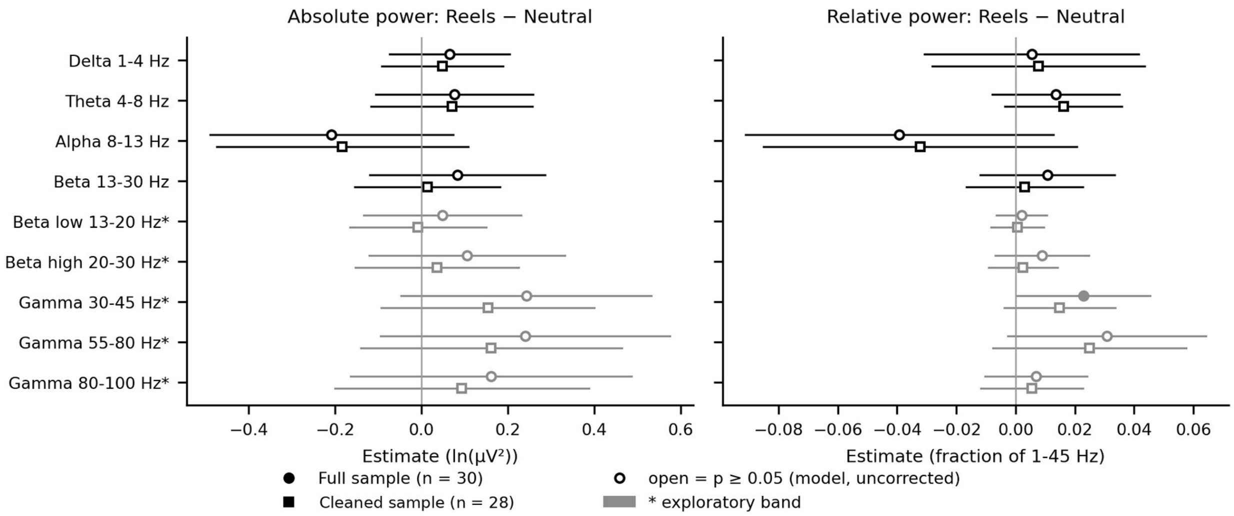


**Supplementary Fig. S1. Intervention EEG: global Reels - neutral differences.** E − N differences in whole-scalp absolute power (left, ln μV²) and relative power (right, fraction of 1-45 Hz power) during the interventions. Points show model-estimated differences with 95% CIs in the full sample (circles; N = 30) and the cleaned sample (squares; N = 28); filled symbols, uncorrected $p < .05$; open symbols, $p \geq .05$; grey, exploratory bands (*).

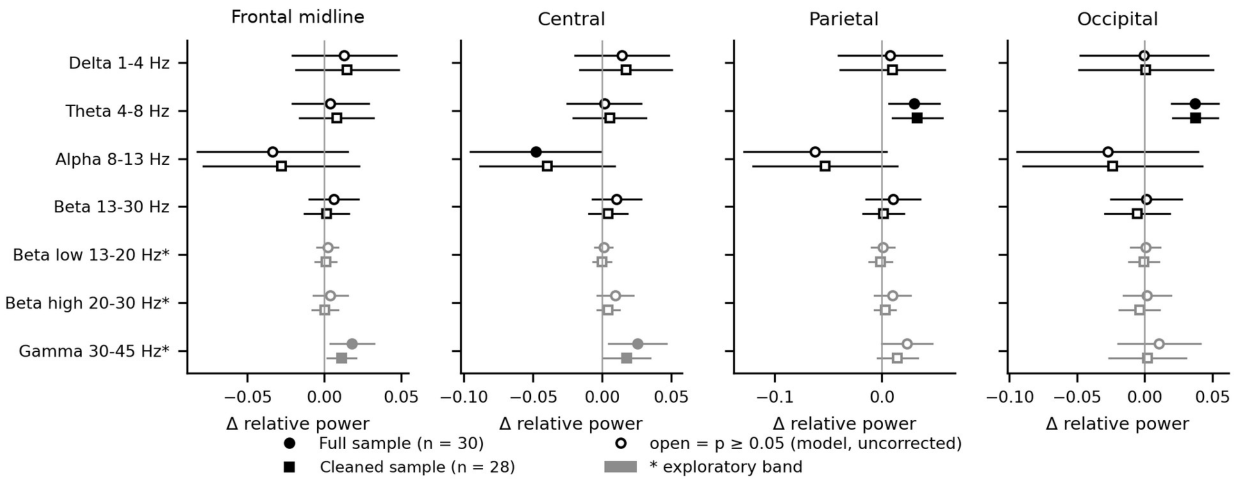


**Supplementary Fig. S2. Intervention EEG: regional Reels - neutral differences.** E − N differences in relative power in the four literature-defined regions (frontal midline, central, parietal, and occipital). Points show model-estimated differences with 95% CIs in the full sample (circles; N = 30) and the cleaned sample (squares; N = 28); filled symbols, uncorrected $p < .05$; open symbols, $p \geq .05$; grey, exploratory bands (*).

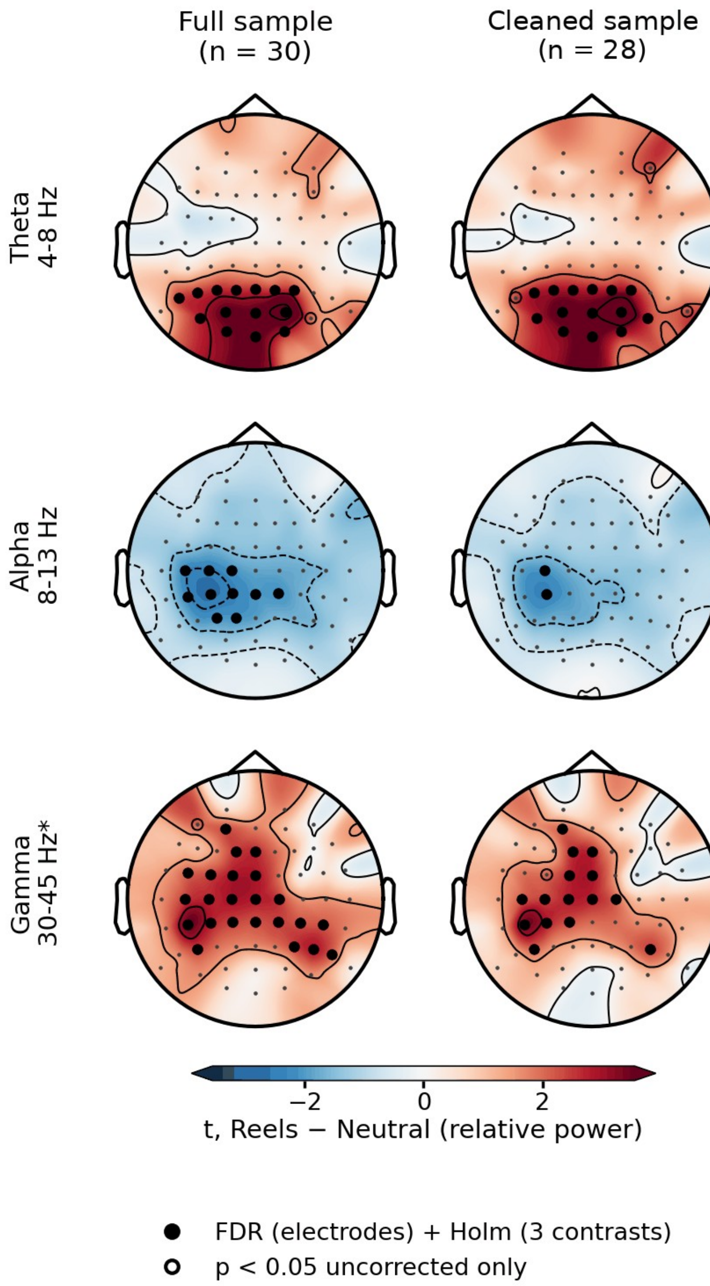


**Supplementary Fig. S3. Intervention EEG: topographies of the Reels - neutral difference.** Electrode-wise *t* statistics (estimate/SE) of the E − N contrast for relative theta, alpha, and 30-45 Hz power in the full (left) and cleaned (right) samples. Filled circles, electrodes surviving FDR correction across electrodes and Holm correction across the three contrasts; open circles, uncorrected $p < .05$ only. The posterior theta increase was similar in the two samples, whereas the left centro-parietal alpha decrease narrowed from 10 to 2 electrodes and the 30-45 Hz increase from 23 to 15 electrodes in the cleaned sample.

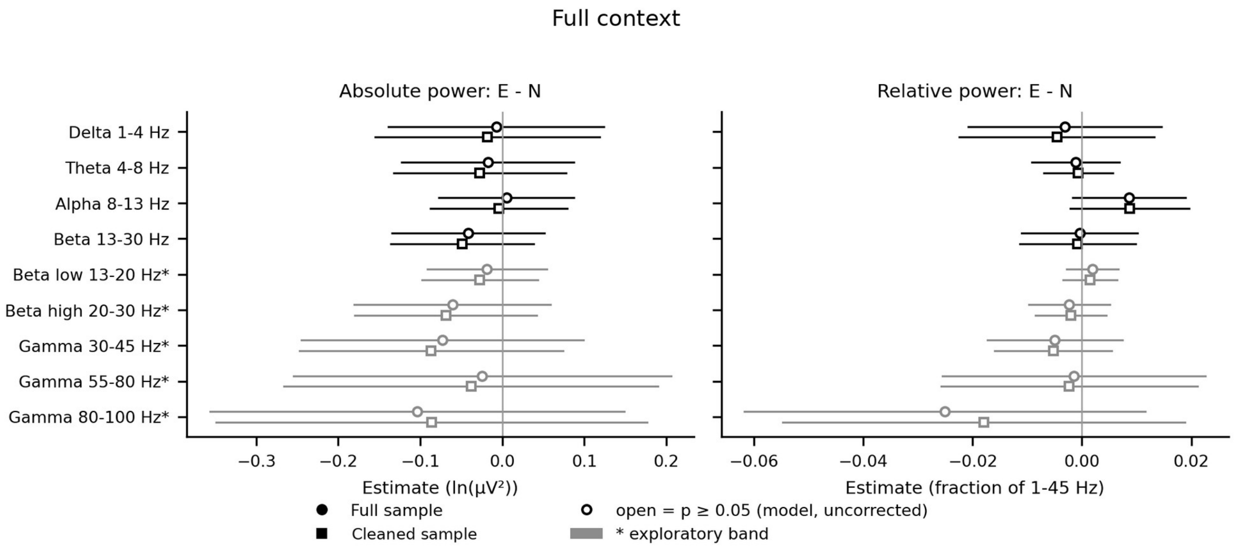


**Supplementary Fig. S4. Task EEG, Full context: global Reels - neutral differences.** E − N differences in whole-scalp absolute (left) and relative (right) power after the interventions; Full context, the N-back and SART recordings analysed together. Points show model-estimated differences with 95% CIs in the full sample (circles; N = 30) and the cleaned sample (squares; N = 28); filled symbols, uncorrected $p < .05$; open symbols, $p \geq .05$; grey, exploratory bands (*).

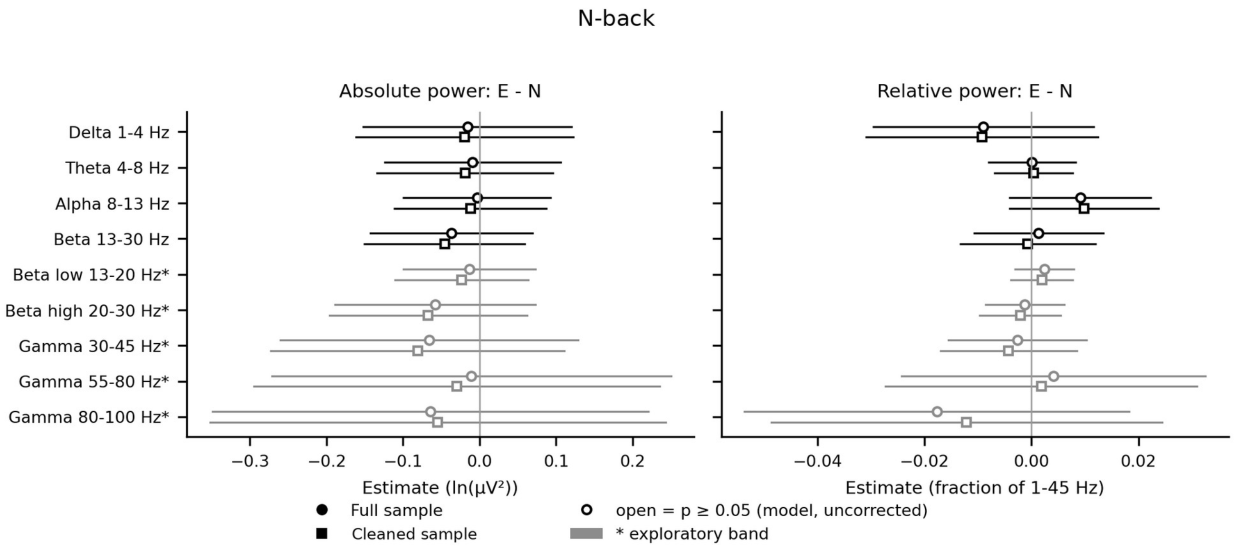


**Supplementary Fig. S5. Task EEG, N-back: global Reels - neutral differences.** E − N differences in whole-scalp absolute (left) and relative (right) power after the interventions. Points show model-estimated differences with 95% CIs in the full sample (circles; N = 30) and the cleaned sample (squares; N = 28); filled symbols, uncorrected $p < .05$; open symbols, $p \geq .05$; grey, exploratory bands (*).

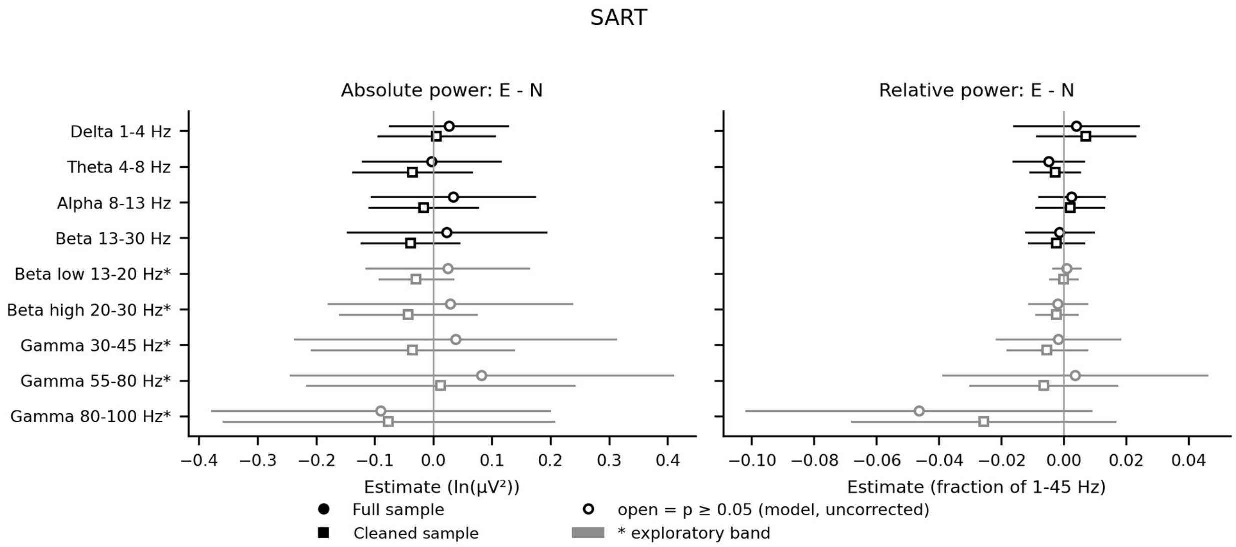


**Supplementary Fig. S6. Task EEG, SART: global Reels - neutral differences.** E − N differences in whole-scalp absolute (left) and relative (right) power after the interventions. Points show model-estimated differences with 95% CIs in the full sample (circles; N = 30) and the cleaned sample (squares; N = 28); filled symbols, uncorrected $p < .05$; open symbols, $p \geq .05$; grey, exploratory bands (*).

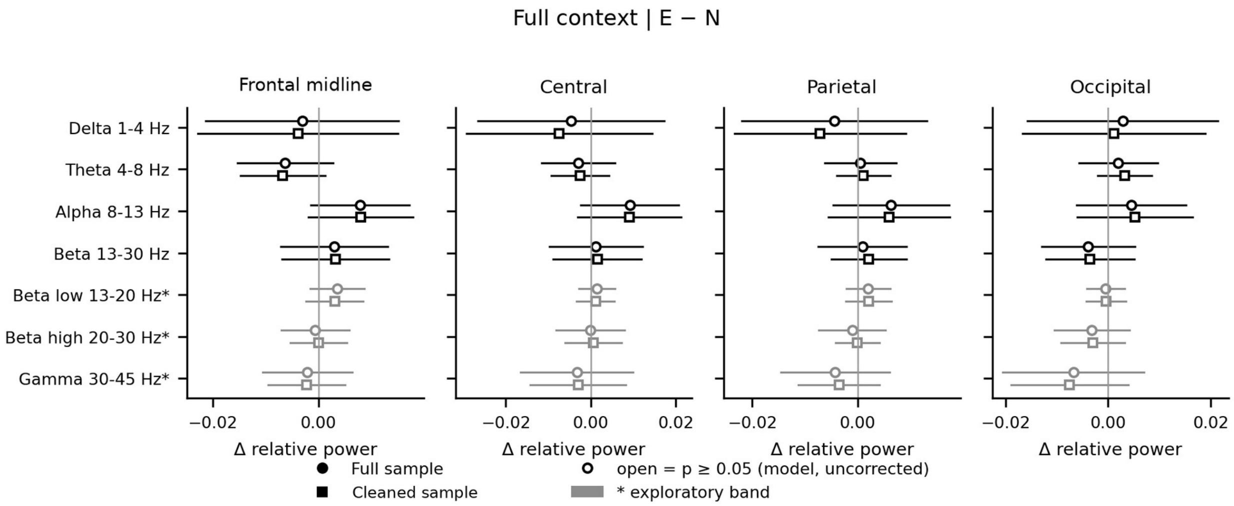


**Supplementary Fig. S7. Task EEG, Full context: regional Reels - neutral differences.** E − N differences in relative power in the four literature-defined regions. Points show model-estimated differences with 95% CIs in the full sample (circles; N = 30) and the cleaned sample (squares; N = 28); filled symbols, uncorrected $p < .05$; open symbols, $p \geq .05$; grey, exploratory bands (*).

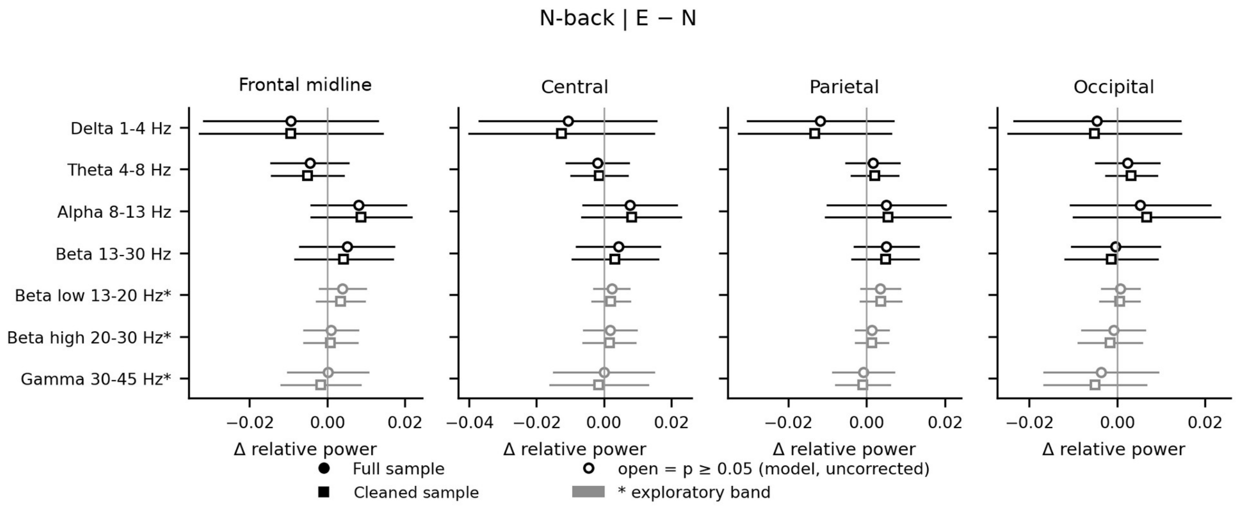


**Supplementary Fig. S8. Task EEG, N-back: regional Reels - neutral differences.** E − N differences in relative power in the four literature-defined regions. Points show model-estimated differences with 95% CIs in the full sample (circles; N = 30) and the cleaned sample (squares; N = 28); filled symbols, uncorrected $p < .05$; open symbols, $p \geq .05$; grey, exploratory bands (*).

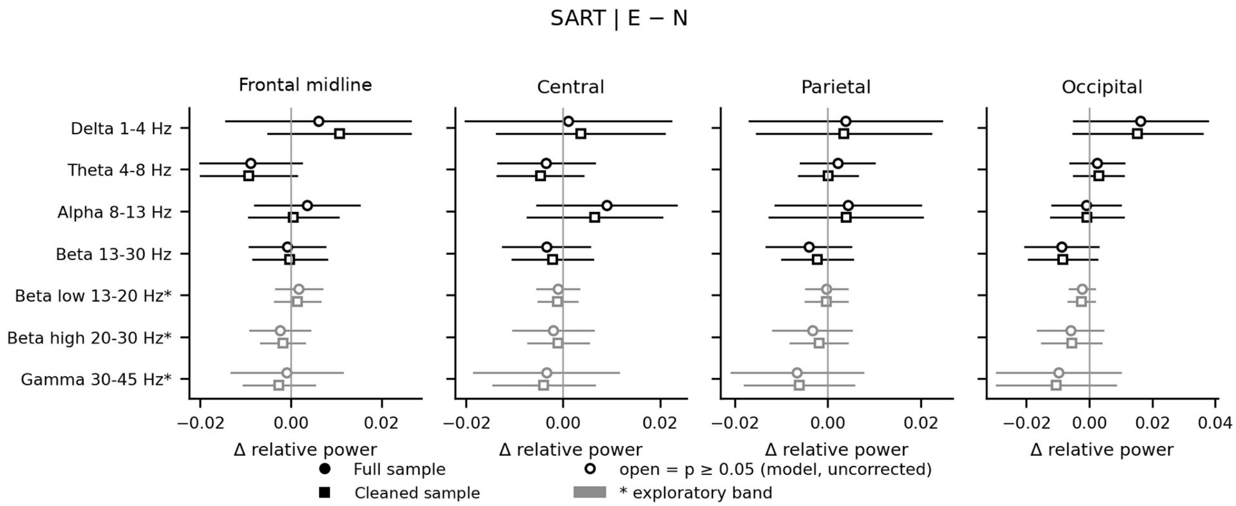


**Supplementary Fig. S9. Task EEG, SART: regional Reels - neutral differences.** E − N differences in relative power in the four literature-defined regions. Points show model-estimated differences with 95% CIs in the full sample (circles; N = 30) and the cleaned sample (squares; N = 28); filled symbols, uncorrected $p < .05$; open symbols, $p \geq .05$; grey, exploratory bands (*).

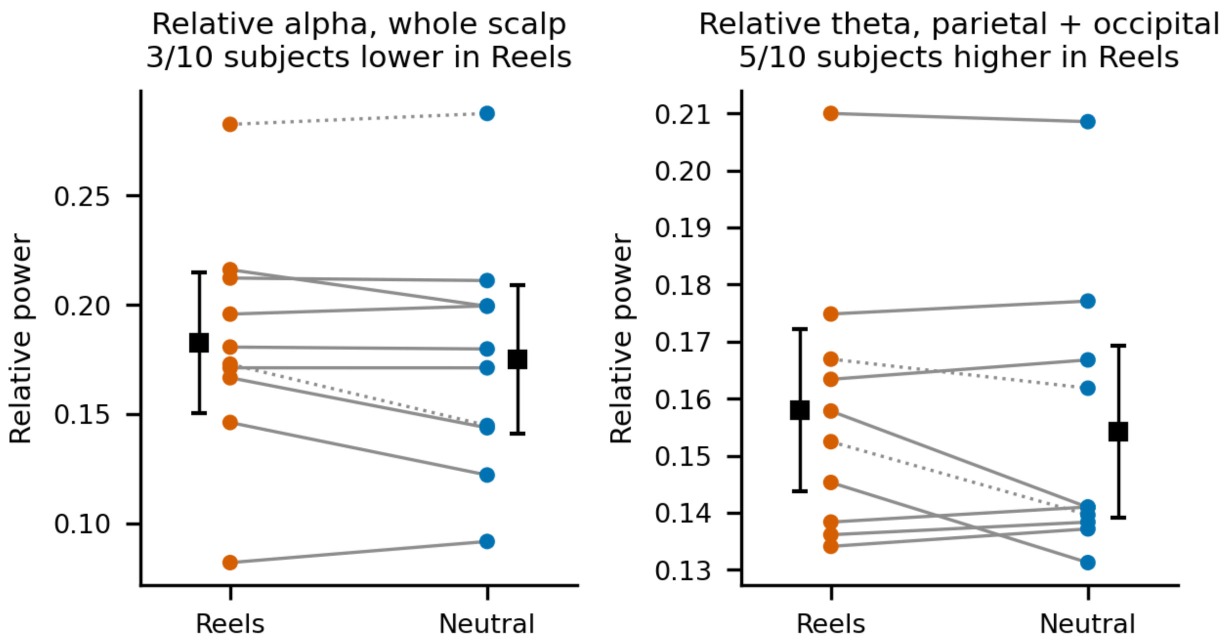


**Supplementary Fig. S10. Task EEG, Full context: within-person values.** Whole-scalp relative alpha (left) and parietal-occipital relative theta (right) after Reels and after neutral images in the 10 participants who completed both interventions. Lines connect the two values of each participant; squares show the mean with its 95% CI. Descriptive, not corrected for multiple comparisons.

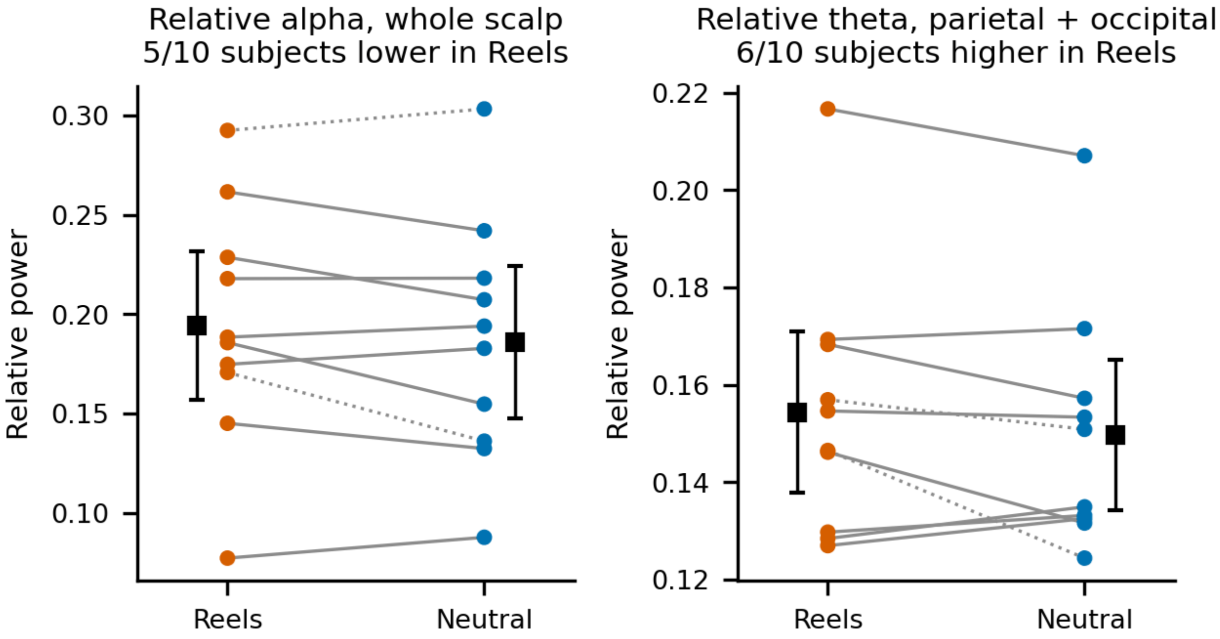


**Supplementary Fig. S11. Task EEG, N-back: within-person values.** Whole-scalp relative alpha (left) and parietal-occipital relative theta (right) after Reels and after neutral images in the 10 participants who completed both interventions. Lines connect the two values of each participant; squares show the mean with its 95% CI. Descriptive, not corrected for multiple comparisons.

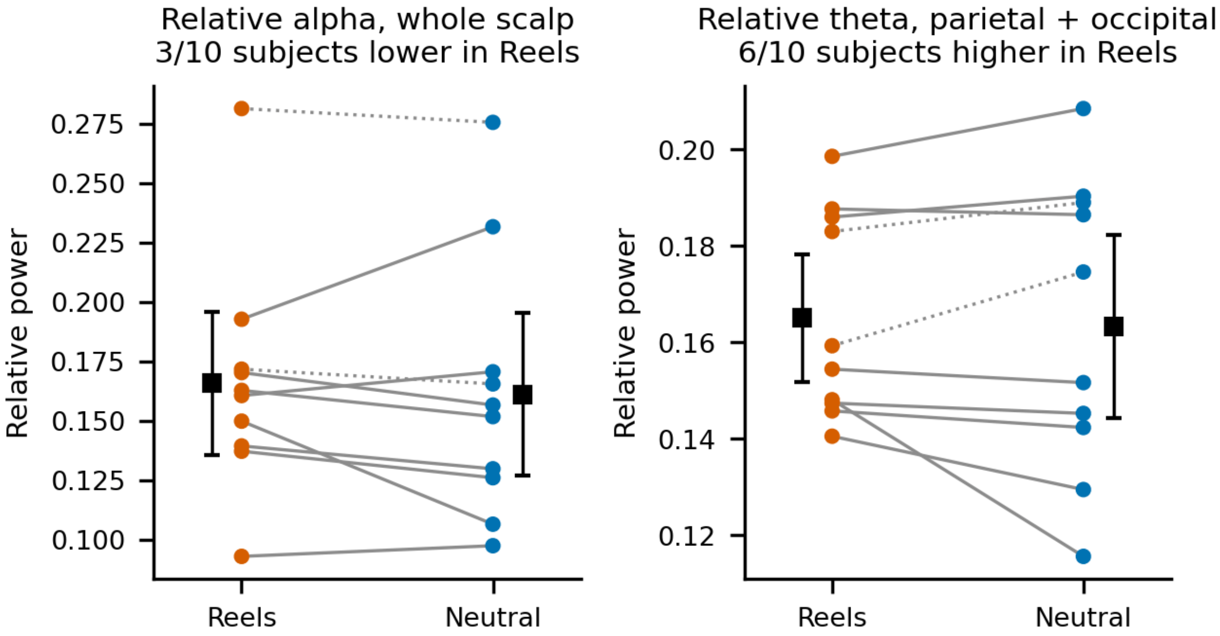


**Supplementary Fig. S12. Task EEG, SART: within-person values.** Whole-scalp relative alpha (left) and parietal-occipital relative theta (right) after Reels and after neutral images in the 10 participants who completed both interventions. Lines connect the two values of each participant; squares show the mean with its 95% CI. Descriptive, not corrected for multiple comparisons.

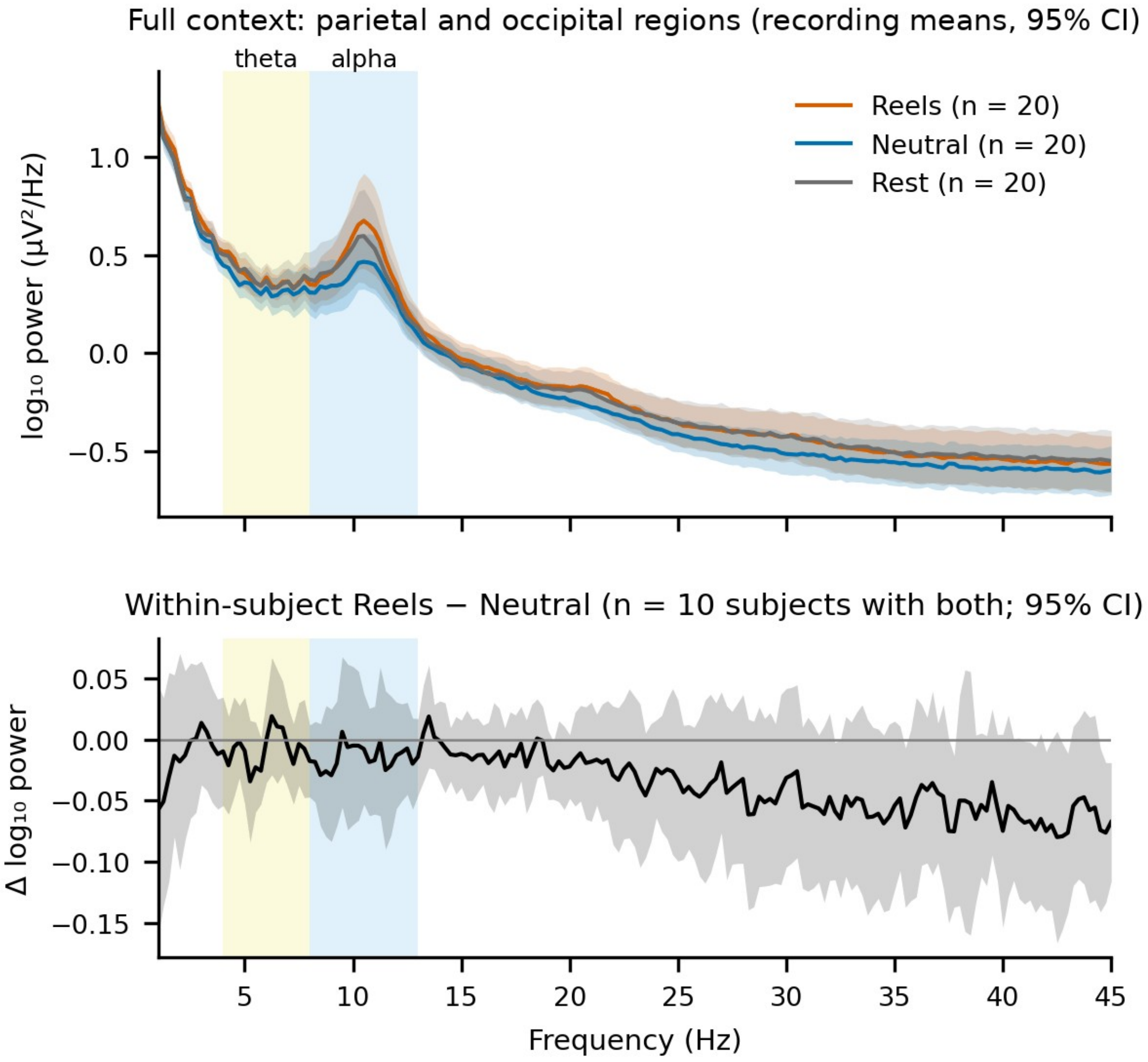


**Supplementary Fig. S13. Task EEG, Full context: posterior power spectra.** Upper panel: parietal-occipital log10 power spectra (recording means, 95% CI) after Reels, neutral images, and rest (n = 20 recordings each), with the theta (4-8 Hz) and alpha (8-13 Hz) bands shaded. Lower panel: within-person Reels − neutral difference spectrum in the 10 participants who completed both interventions (95% CI).

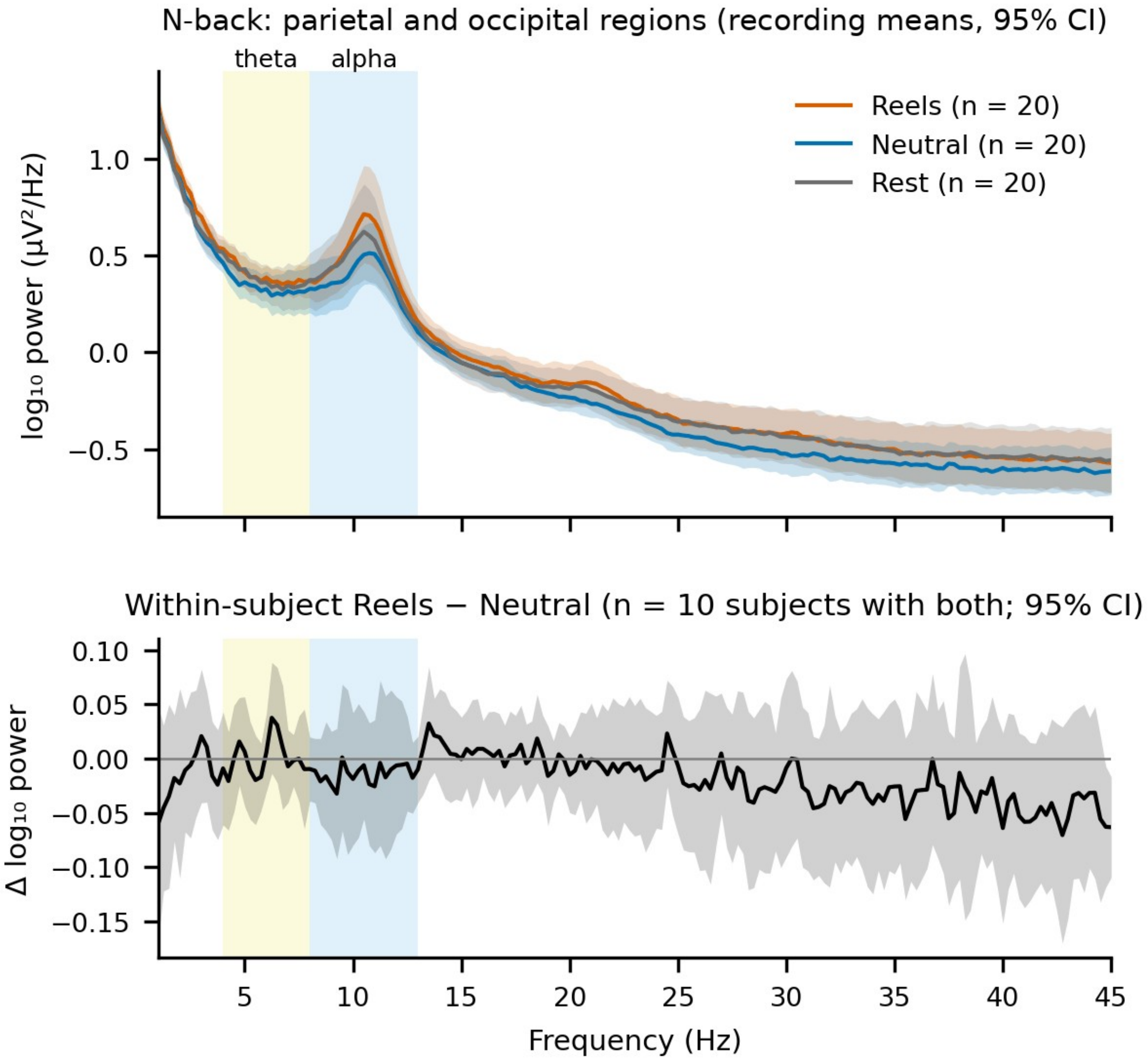


**Supplementary Fig. S14. Task EEG, N-back: posterior power spectra.** Upper panel: parietal-occipital log10 power spectra (recording means, 95% CI) after Reels, neutral images, and rest (n = 20 recordings each), with the theta (4-8 Hz) and alpha (8-13 Hz) bands shaded. Lower panel: within-person Reels − neutral difference spectrum in the 10 participants who completed both interventions (95% CI).

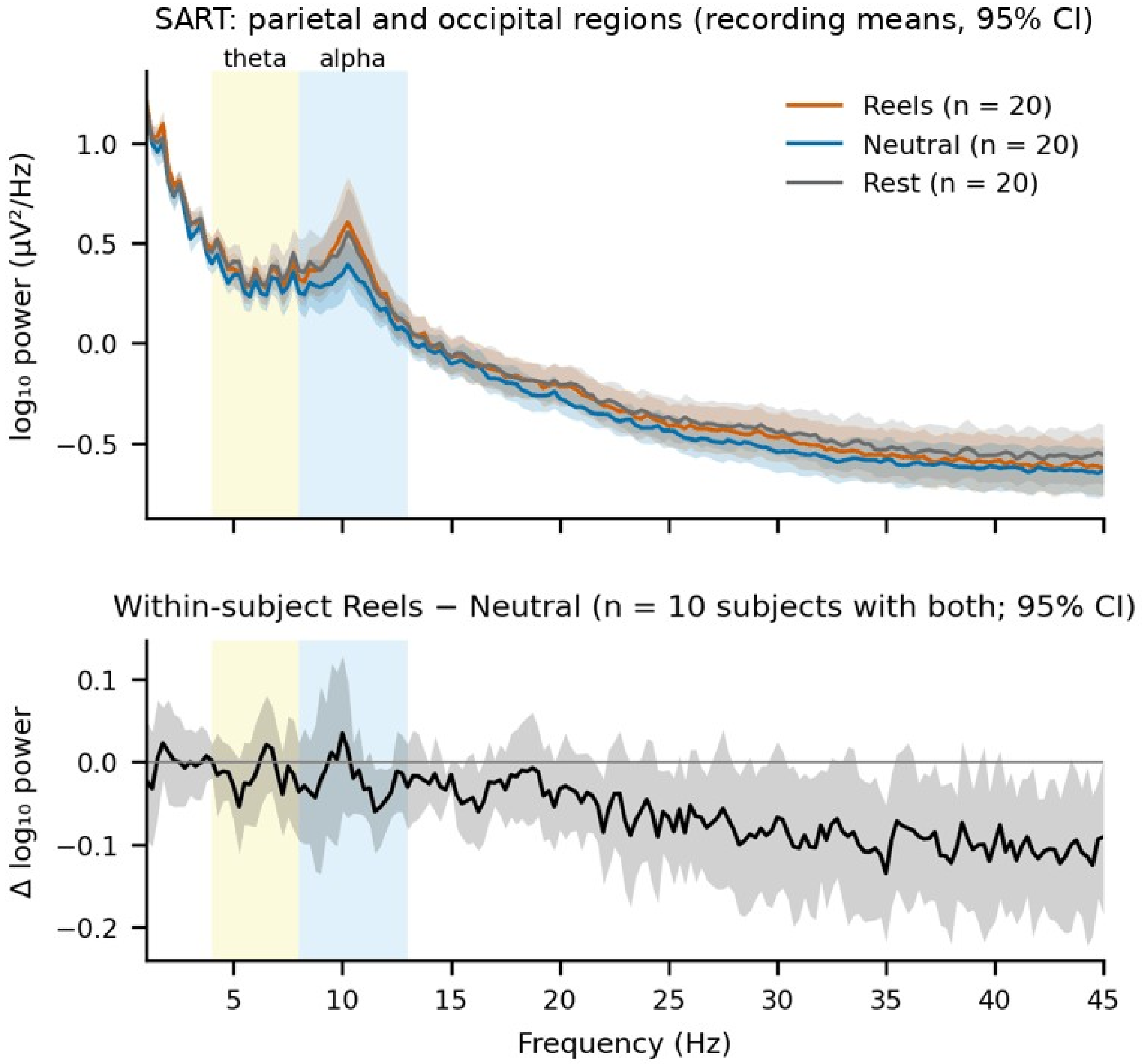


**Supplementary Fig. S15. Task EEG, SART: posterior power spectra.** Upper panel: parietal-occipital log10 power spectra (recording means, 95% CI) after Reels, neutral images, and rest (n = 20 recordings each), with the theta (4-8 Hz) and alpha (8-13 Hz) bands shaded. Lower panel: within-person Reels − neutral difference spectrum in the 10 participants who completed both interventions (95% CI).

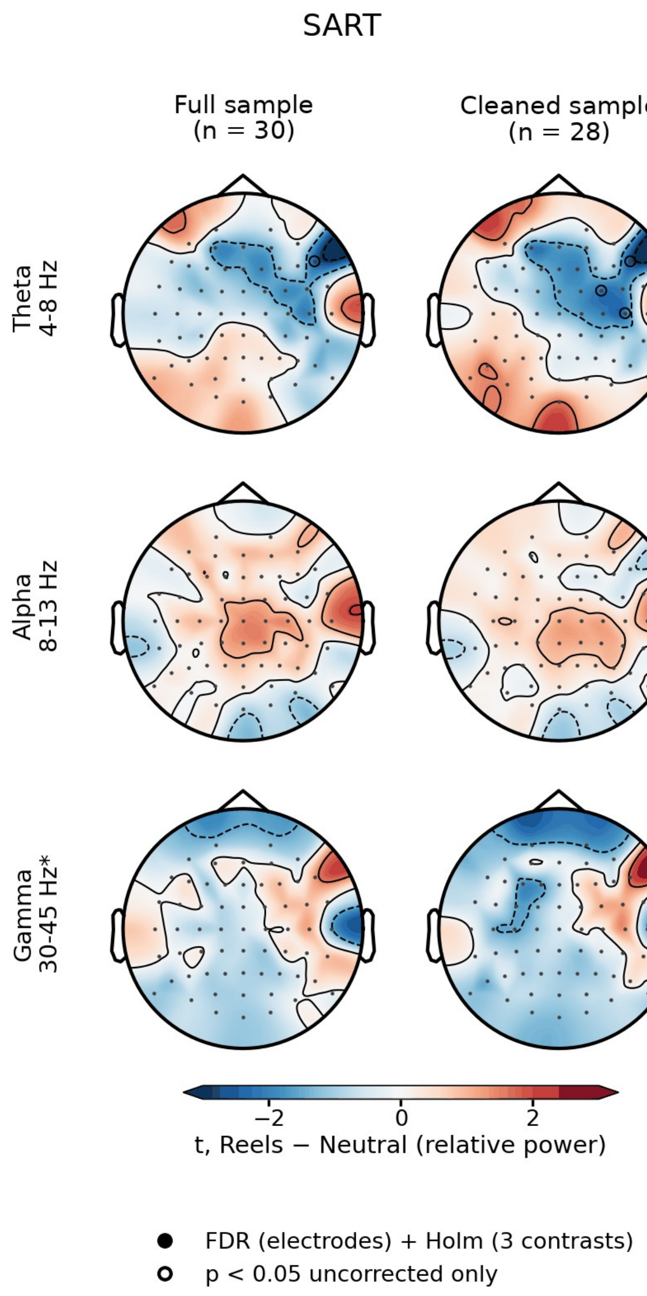


**Supplementary Fig. S16. Task EEG, SART: topographies of the Reels - neutral difference.** Electrode-wise *t* statistics of the E − N contrast for relative theta, alpha, and 30-45 Hz power in the full (left) and cleaned (right) samples. Filled circles, electrodes surviving FDR correction across electrodes and Holm correction across the three contrasts; open circles, uncorrected $p < .05$ only.

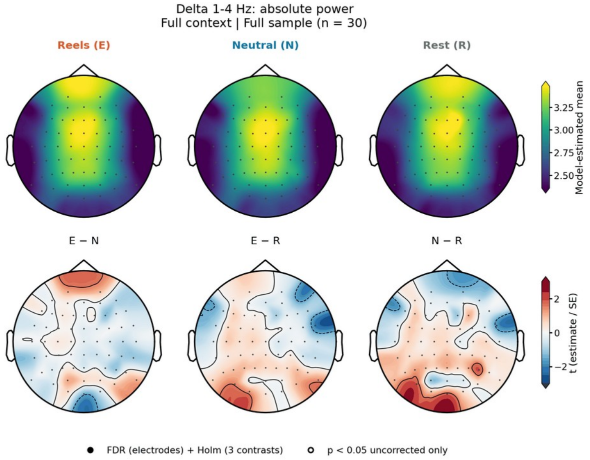
Delta 1-4 Hz: absolute power
Full context | Full sample (n = 30)
Reels (E)
Neutral (N)
Rest (R)
Model-estimated mean
E − N
E − R
N − R
t (estimate / SE)
FDR (electrodes) + Holm (3 contrasts)
p < 0.05 uncorrected only

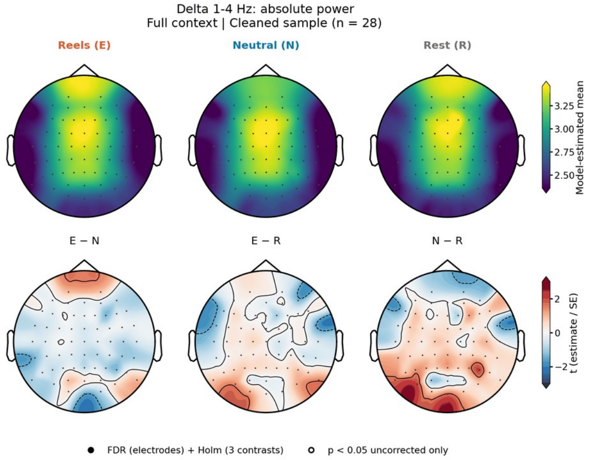
Delta 1-4 Hz: absolute power
Full context | Cleaned sample (n = 28)
Reels (E)
Neutral (N)
Rest (R)
Model-estimated mean
E − N
E − R
N − R
t (estimate / SE)
FDR (electrodes) + Holm (3 contrasts)
p < 0.05 uncorrected only

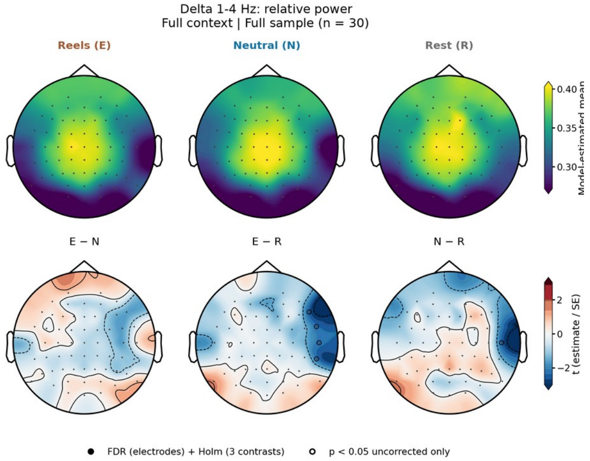
Delta 1-4 Hz: relative power
Full context | Full sample (n = 30)
Reels (E)
Neutral (N)
Rest (R)
E − N
E − R
N − R
t (estimate / SE)
FDR (electrodes) + Holm (3 contrasts)
p < 0.05 uncorrected only

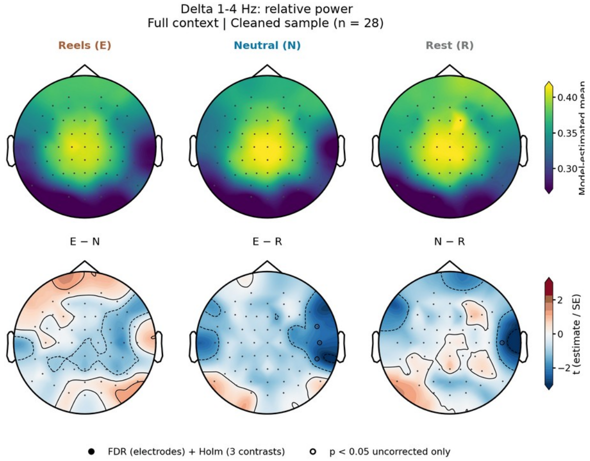
Delta 1-4 Hz: relative power
Full context | Cleaned sample (n = 28)
Reels (E)
Neutral (N)
Rest (R)
E − N
E − R
N − R
t (estimate / SE)
FDR (electrodes) + Holm (3 contrasts)
p < 0.05 uncorrected only

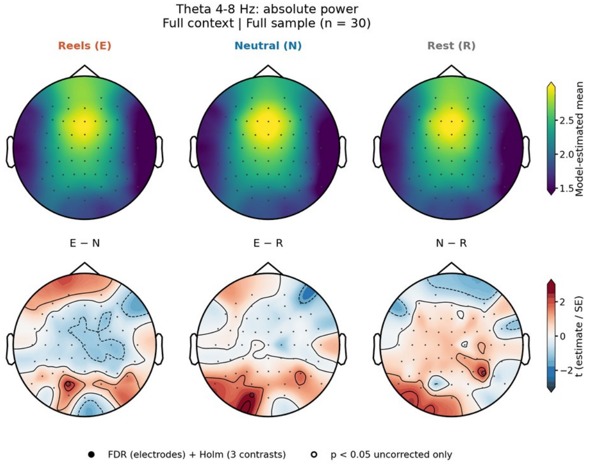
Theta 4-8 Hz: absolute power
Full context | Full sample (n = 30)
Reels (E)
Neutral (N)
Rest (R)
Model-estimated mean
E − N
E − R
N − R
t (estimate / SE)
FDR (electrodes) + Holm (3 contrasts)
p < 0.05 uncorrected only

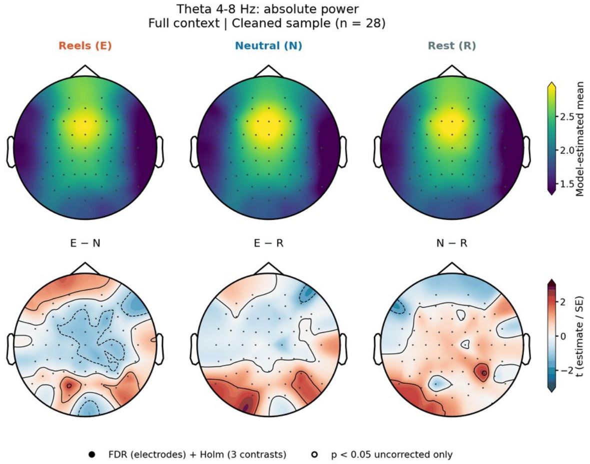
Theta 4-8 Hz: absolute power
Full context | Cleaned sample (n = 28)
Reels (E)
Neutral (N)
Rest (R)
Model-estimated mean
E − N
E − R
N − R
t (estimate / SE)
FDR (electrodes) + Holm (3 contrasts)
p < 0.05 uncorrected only

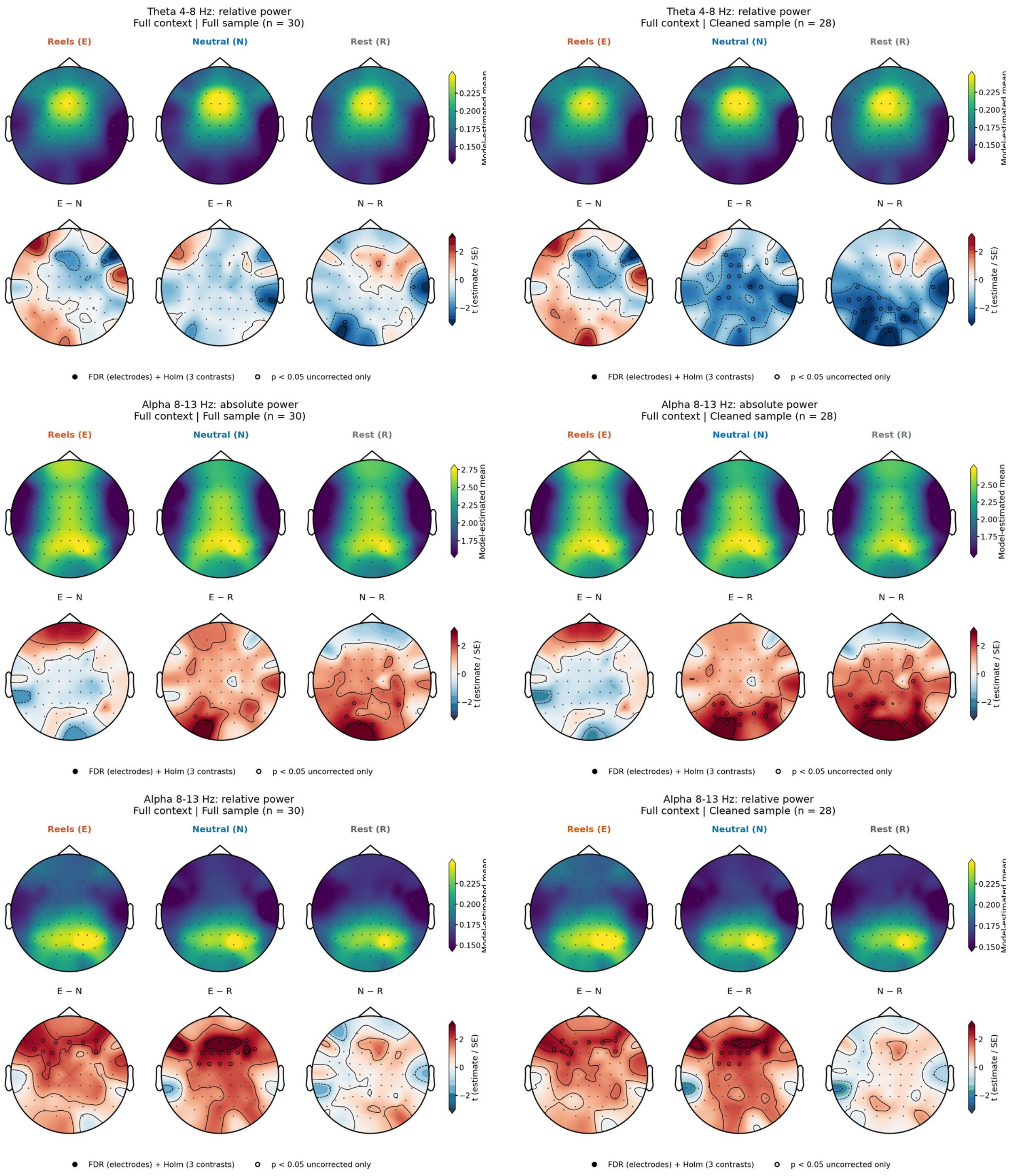
Theta 4-8 Hz: relative power
Full context | Full sample (n = 30)
Theta 4-8 Hz: relative power
Full context | Cleaned sample (n = 28)
Alpha 8-13 Hz: absolute power
Full context | Full sample (n = 30)
Alpha 8-13 Hz: absolute power
Full context | Cleaned sample (n = 28)
Alpha 8-13 Hz: relative power
Full context | Full sample (n = 30)
Alpha 8-13 Hz: relative power
Full context | Cleaned sample (n = 28)
Reels (E)
Neutral (N)
Rest (R)
E − N
E − R
N − R
Model-estimated mean
t (estimate / SE)
FDR (electrodes) + Holm (3 contrasts)
p < 0.05 uncorrected only

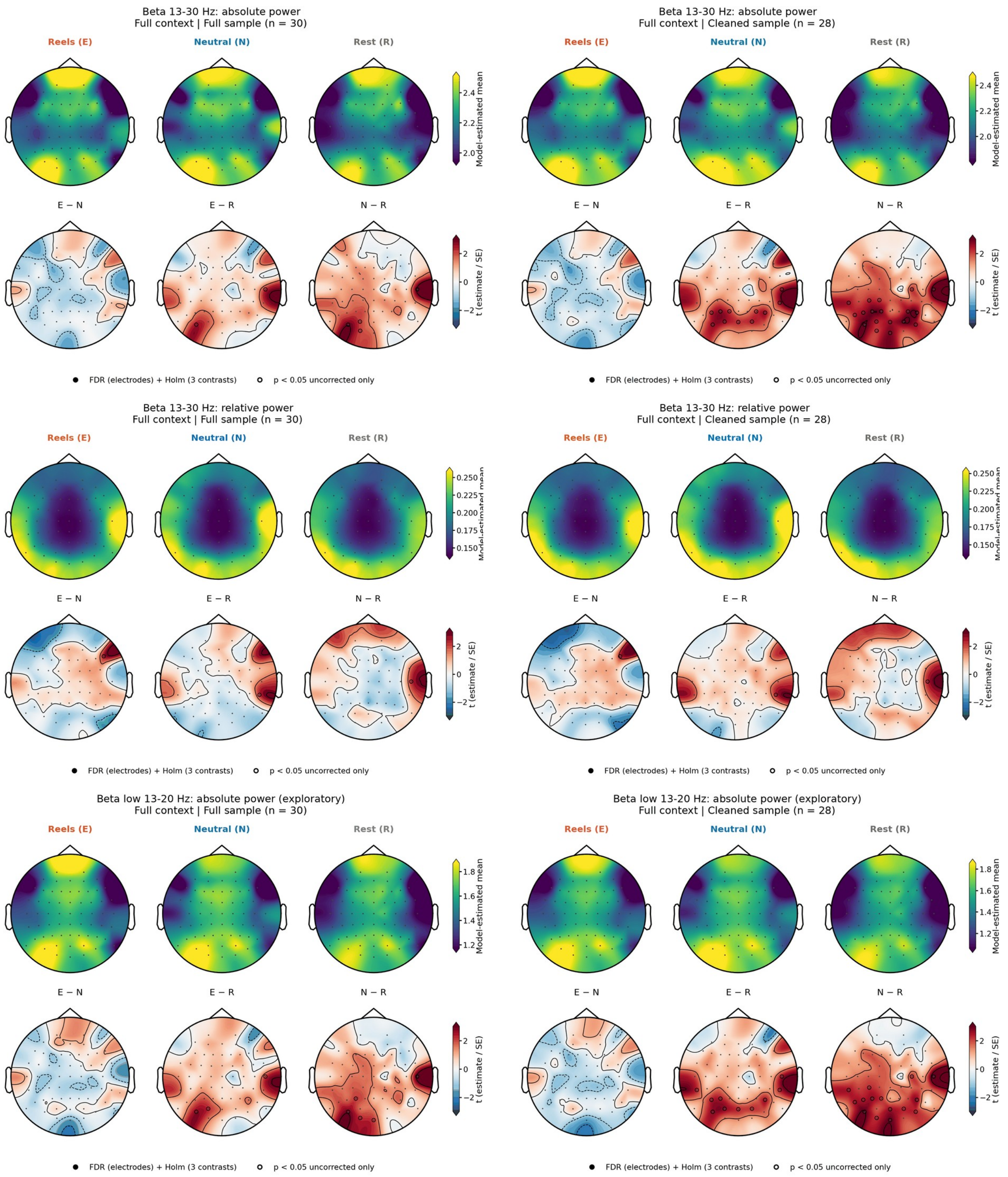
Beta 13-30 Hz: absolute power
Full context | Full sample (n = 30)
Reels (E)
Neutral (N)
Rest (R)
Model-estimated mean
E − N
E − R
N − R
t (estimate / SE)
FDR (electrodes) + Holm (3 contrasts)
p < 0.05 uncorrected only
Beta 13-30 Hz: absolute power
Full context | Cleaned sample (n = 28)
Reels (E)
Neutral (N)
Rest (R)
Model-estimated mean
E − N
E − R
N − R
t (estimate / SE)
FDR (electrodes) + Holm (3 contrasts)
p < 0.05 uncorrected only
Beta 13-30 Hz: relative power
Full context | Full sample (n = 30)
Reels (E)
Neutral (N)
Rest (R)
E − N
E − R
N − R
t (estimate / SE)
FDR (electrodes) + Holm (3 contrasts)
p < 0.05 uncorrected only
Beta 13-30 Hz: relative power
Full context | Cleaned sample (n = 28)
Reels (E)
Neutral (N)
Rest (R)
E − N
E − R
N − R
t (estimate / SE)
FDR (electrodes) + Holm (3 contrasts)
p < 0.05 uncorrected only
Beta low 13-20 Hz: absolute power (exploratory)
Full context | Full sample (n = 30)
Reels (E)
Neutral (N)
Rest (R)
Model-estimated mean
E − N
E − R
N − R
t (estimate / SE)
FDR (electrodes) + Holm (3 contrasts)
p < 0.05 uncorrected only
Beta low 13-20 Hz: absolute power (exploratory)
Full context | Cleaned sample (n = 28)
Reels (E)
Neutral (N)
Rest (R)
Model-estimated mean
E − N
E − R
N − R
t (estimate / SE)
FDR (electrodes) + Holm (3 contrasts)
p < 0.05 uncorrected only

Beta low 13-20 Hz: relative power (exploratory)
Full context | Full sample (n = 30)

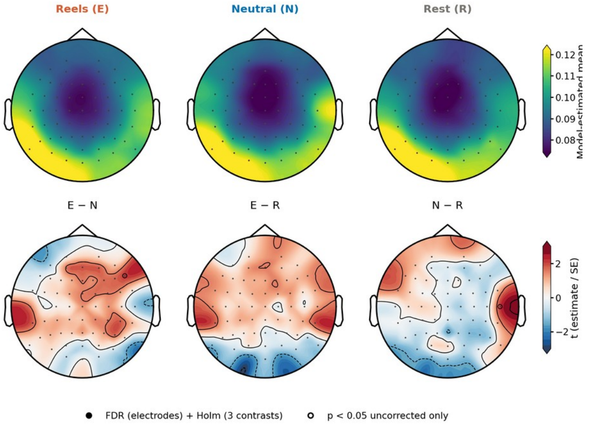


Beta low 13-20 Hz: relative power (exploratory)
Full context | Cleaned sample (n = 28)

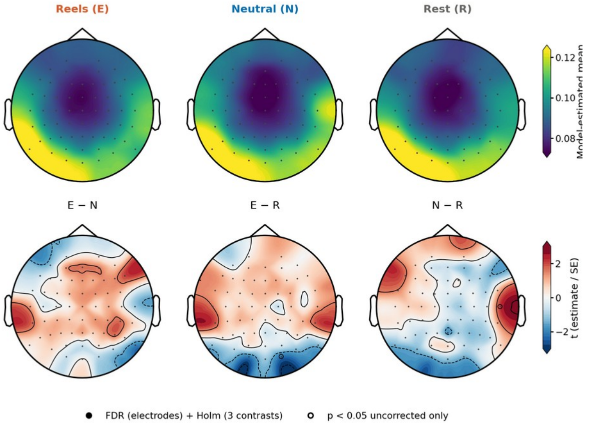


Beta high 20-30 Hz: absolute power (exploratory)
Full context | Full sample (n = 30)

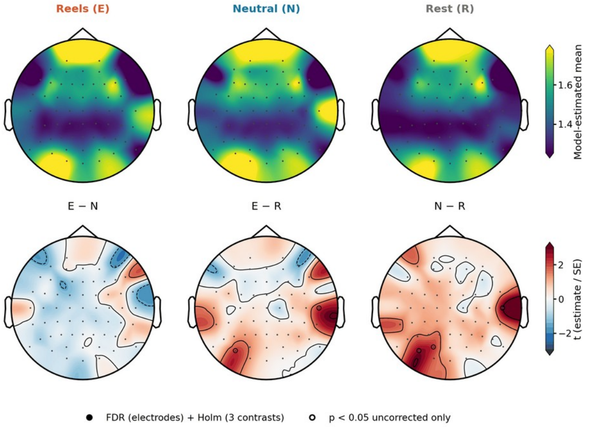


Beta high 20-30 Hz: absolute power (exploratory)
Full context | Cleaned sample (n = 28)

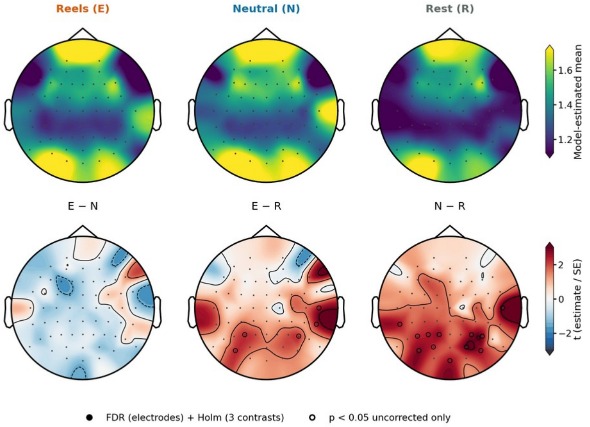


Beta high 20-30 Hz: relative power (exploratory)
Full context | Full sample (n = 30)

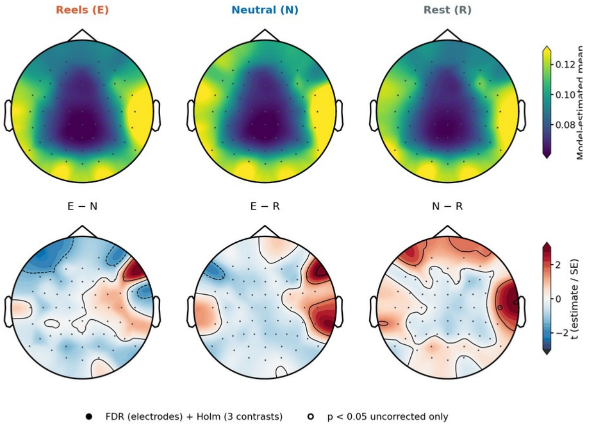


Beta high 20-30 Hz: relative power (exploratory)
Full context | Cleaned sample (n = 28)

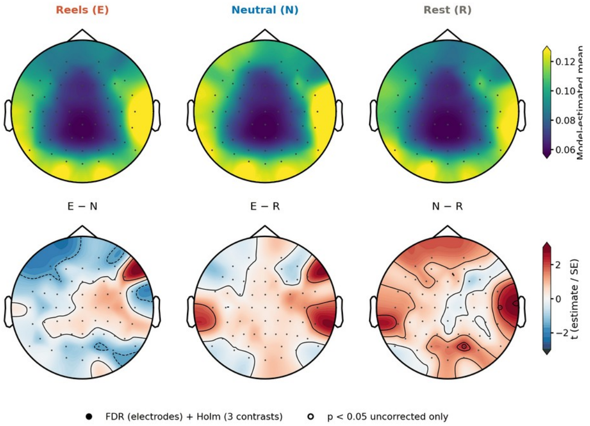

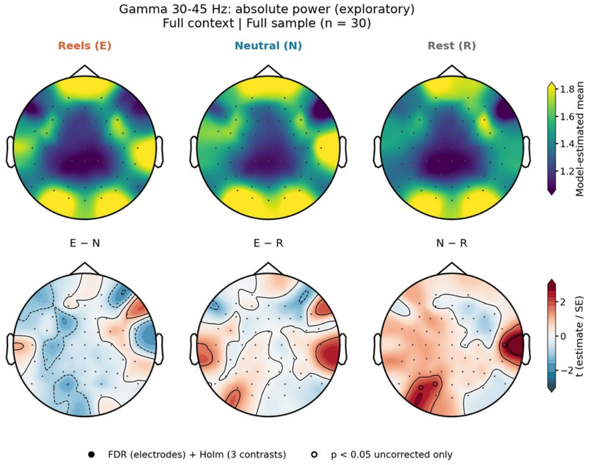
Gamma 30-45 Hz: absolute power (exploratory)
Full context | Full sample (n = 30)
Reels (E)
Neutral (N)
Rest (R)
Model-estimated mean
E − N
E − R
N − R
t (estimate / SE)
FDR (electrodes) + Holm (3 contrasts)
p < 0.05 uncorrected only

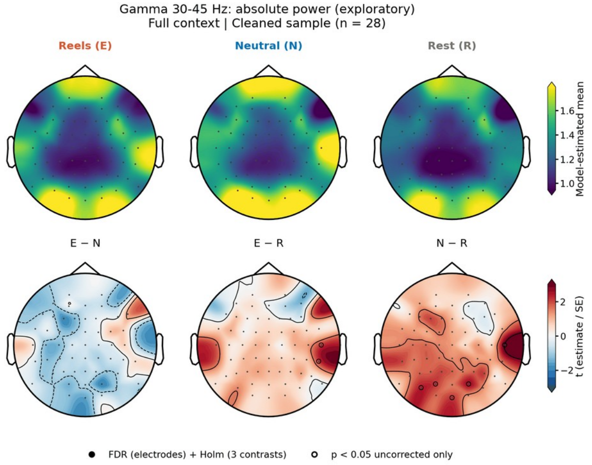
Gamma 30-45 Hz: absolute power (exploratory)
Full context | Cleaned sample (n = 28)
Reels (E)
Neutral (N)
Rest (R)
Model-estimated mean
E − N
E − R
N − R
t (estimate / SE)
FDR (electrodes) + Holm (3 contrasts)
p < 0.05 uncorrected only

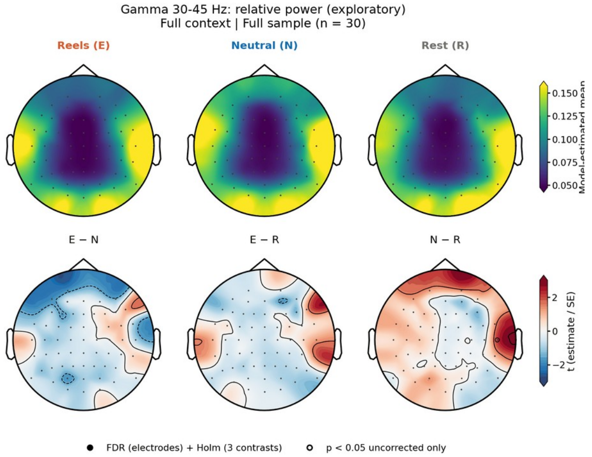
Gamma 30-45 Hz: relative power (exploratory)
Full context | Full sample (n = 30)
Reels (E)
Neutral (N)
Rest (R)
E − N
E − R
N − R
t (estimate / SE)
FDR (electrodes) + Holm (3 contrasts)
p < 0.05 uncorrected only

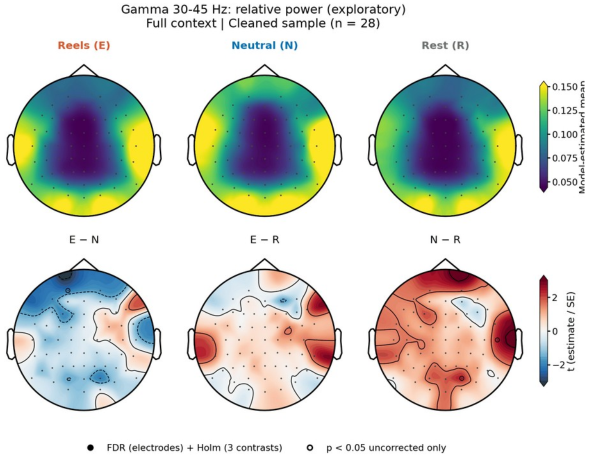
Gamma 30-45 Hz: relative power (exploratory)
Full context | Cleaned sample (n = 28)
Reels (E)
Neutral (N)
Rest (R)
E − N
E − R
N − R
t (estimate / SE)
FDR (electrodes) + Holm (3 contrasts)
p < 0.05 uncorrected only

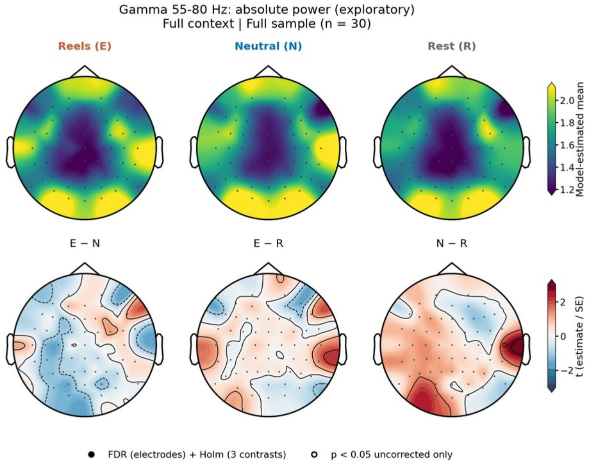
Gamma 55-80 Hz: absolute power (exploratory)
Full context | Full sample (n = 30)
Reels (E)
Neutral (N)
Rest (R)
Model-estimated mean
E − N
E − R
N − R
t (estimate / SE)
FDR (electrodes) + Holm (3 contrasts)
p < 0.05 uncorrected only

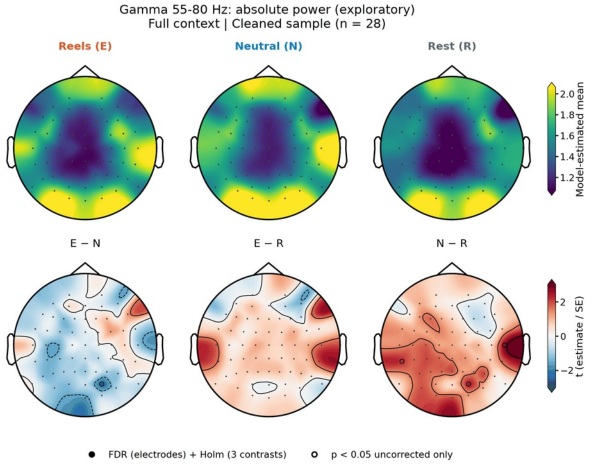
Gamma 55-80 Hz: absolute power (exploratory)
Full context | Cleaned sample (n = 28)
Reels (E)
Neutral (N)
Rest (R)
Model-estimated mean
E − N
E − R
N − R
t (estimate / SE)
FDR (electrodes) + Holm (3 contrasts)
p < 0.05 uncorrected only

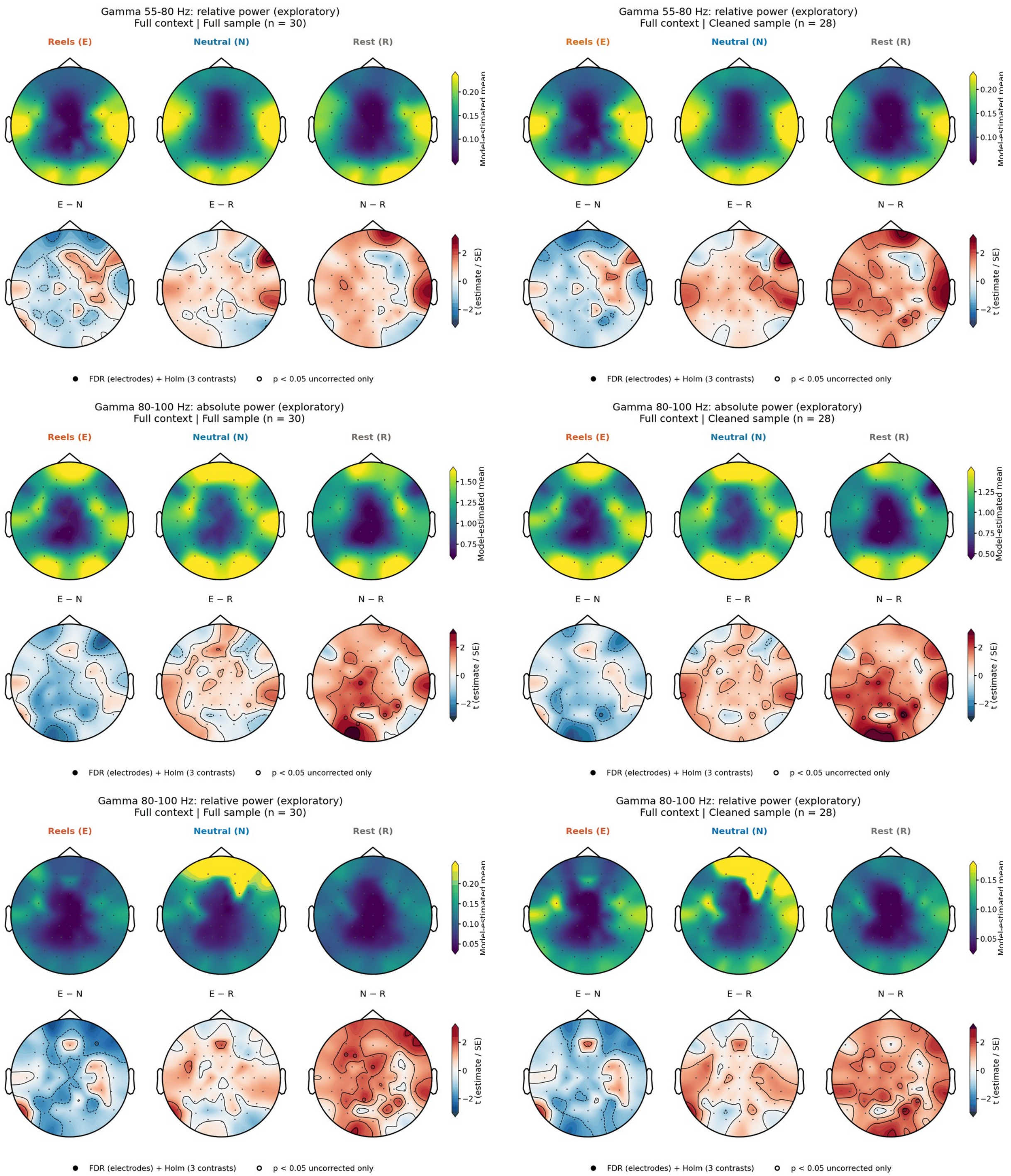


**Supplementary Fig. S17. Task EEG, Full context: electrode topographies for all frequency bands.** For each band and measure (absolute power, ln µV²; relative power, fraction of 1-45 Hz power), the left panel shows the full sample (N = 30) and the right panel the cleaned sample (N = 28). Upper row of each panel: model-estimated means after Reels (E), neutral images (N), and rest (R); lower row: *t* statistics (estimate/SE) of the E − N, E − R, and N − R contrasts. Filled circles, electrodes surviving FDR correction across electrodes and Holm correction across the three contrasts; open circles, uncorrected $p < .05$ only. Bands marked exploratory were not part of the primary analyses.

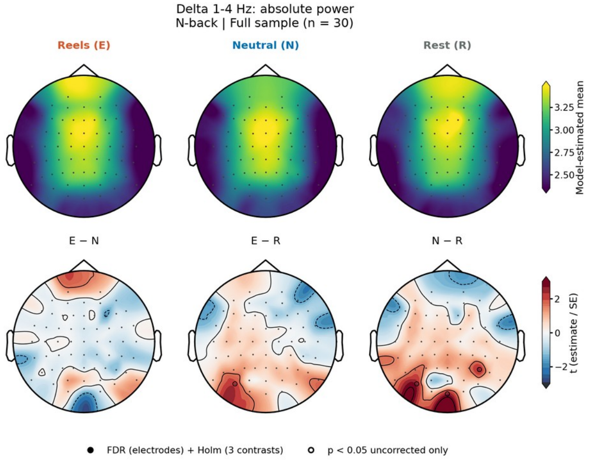
Delta 1-4 Hz: absolute power
N-back | Full sample (n = 30)
Reels (E)
Neutral (N)
Rest (R)
Model-estimated mean
E − N
E − R
N − R
t (estimate / SE)
FDR (electrodes) + Holm (3 contrasts)
p < 0.05 uncorrected only

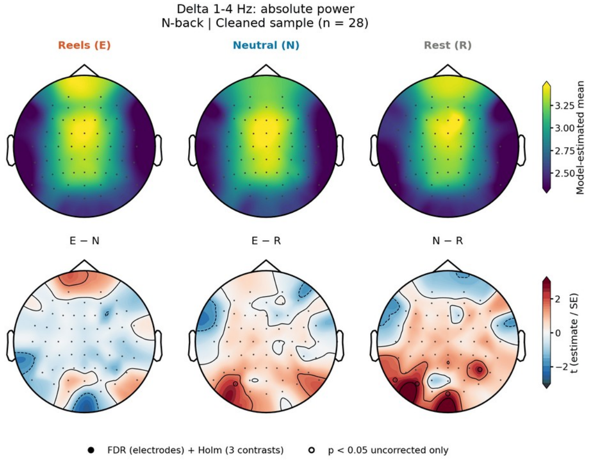
Delta 1-4 Hz: absolute power
N-back | Cleaned sample (n = 28)
Reels (E)
Neutral (N)
Rest (R)
Model-estimated mean
E − N
E − R
N − R
t (estimate / SE)
FDR (electrodes) + Holm (3 contrasts)
p < 0.05 uncorrected only

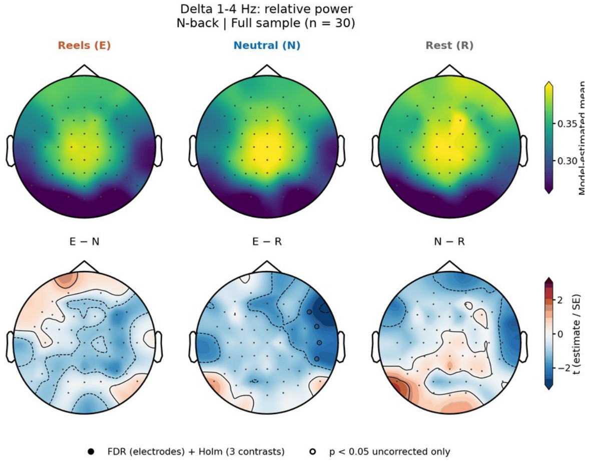
Delta 1-4 Hz: relative power
N-back | Full sample (n = 30)
Reels (E)
Neutral (N)
Rest (R)
E − N
E − R
N − R
t (estimate / SE)
FDR (electrodes) + Holm (3 contrasts)
p < 0.05 uncorrected only

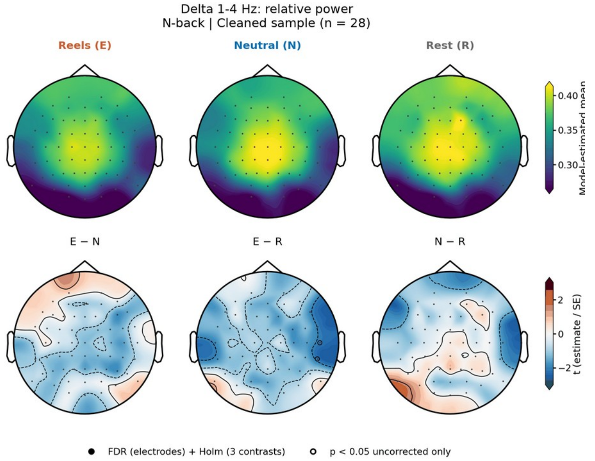
Delta 1-4 Hz: relative power
N-back | Cleaned sample (n = 28)
Reels (E)
Neutral (N)
Rest (R)
E − N
E − R
N − R
t (estimate / SE)
FDR (electrodes) + Holm (3 contrasts)
p < 0.05 uncorrected only

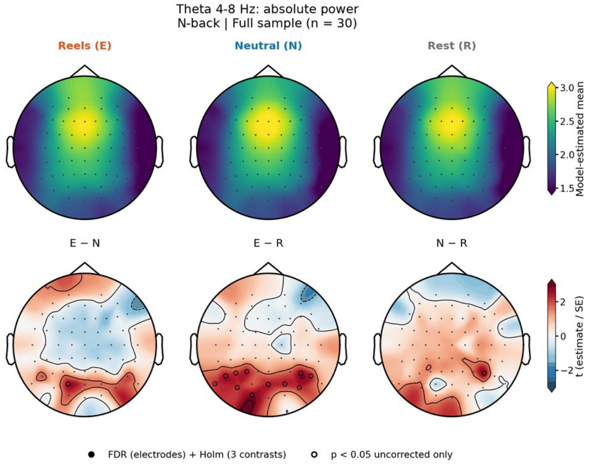
Theta 4-8 Hz: absolute power
N-back | Full sample (n = 30)
Reels (E)
Neutral (N)
Rest (R)
Model-estimated mean
E − N
E − R
N − R
t (estimate / SE)
FDR (electrodes) + Holm (3 contrasts)
p < 0.05 uncorrected only

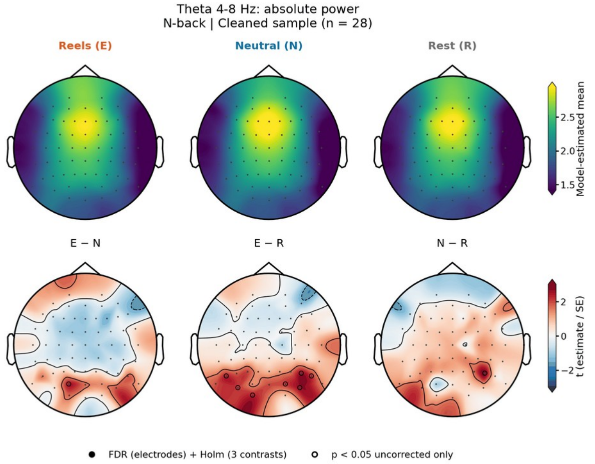
Theta 4-8 Hz: absolute power
N-back | Cleaned sample (n = 28)
Reels (E)
Neutral (N)
Rest (R)
Model-estimated mean
E − N
E − R
N − R
t (estimate / SE)
FDR (electrodes) + Holm (3 contrasts)
p < 0.05 uncorrected only

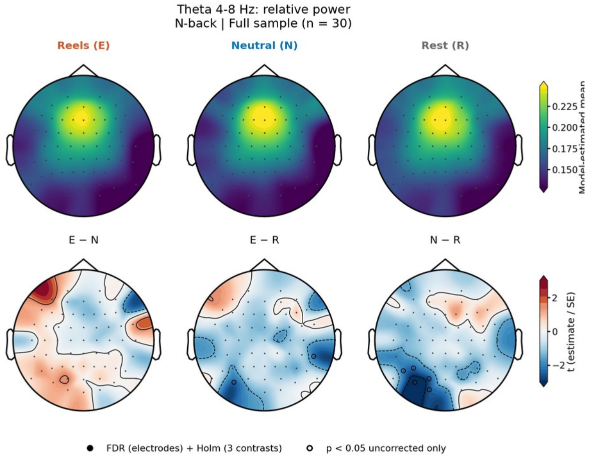
Theta 4-8 Hz: relative power
N-back | Full sample (n = 30)
Reels (E)
Neutral (N)
Rest (R)
Model-estimated mean
E − N
E − R
N − R
t (estimate / SE)
FDR (electrodes) + Holm (3 contrasts)
p < 0.05 uncorrected only

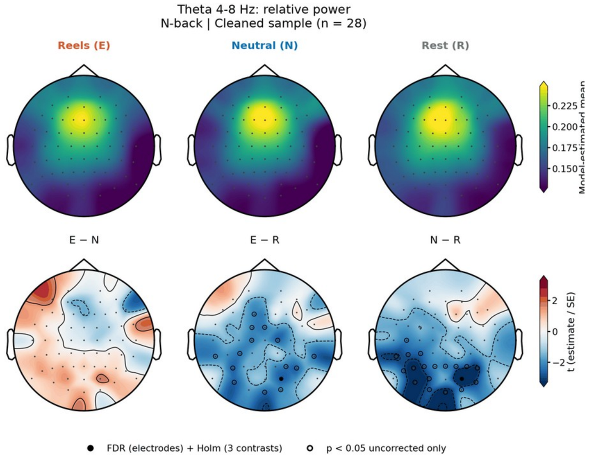
Theta 4-8 Hz: relative power
N-back | Cleaned sample (n = 28)
Reels (E)
Neutral (N)
Rest (R)
Model-estimated mean
E − N
E − R
N − R
t (estimate / SE)
FDR (electrodes) + Holm (3 contrasts)
p < 0.05 uncorrected only

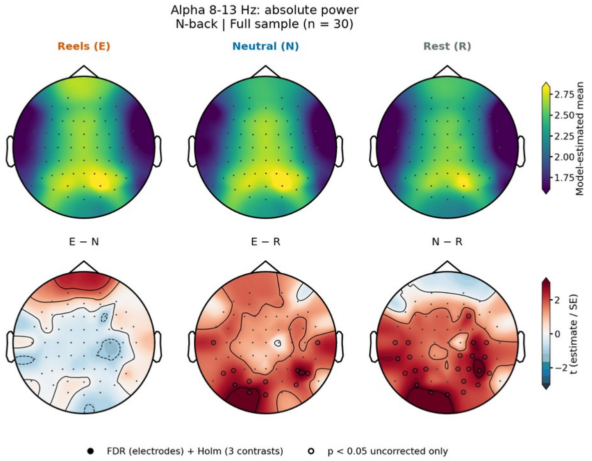
Alpha 8-13 Hz: absolute power
N-back | Full sample (n = 30)
Reels (E)
Neutral (N)
Rest (R)
Model-estimated mean
E − N
E − R
N − R
t (estimate / SE)
FDR (electrodes) + Holm (3 contrasts)
p < 0.05 uncorrected only

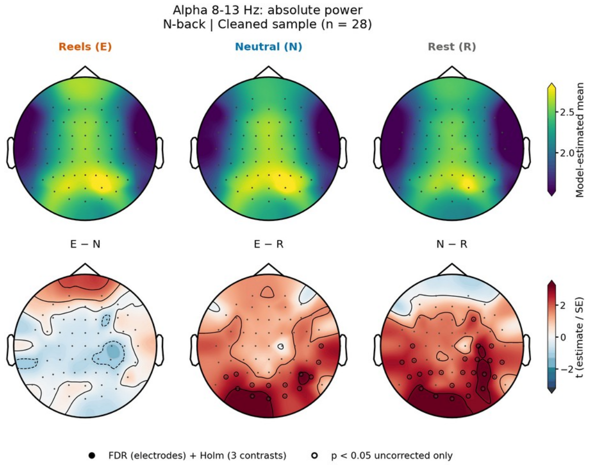
Alpha 8-13 Hz: absolute power
N-back | Cleaned sample (n = 28)
Reels (E)
Neutral (N)
Rest (R)
Model-estimated mean
E − N
E − R
N − R
t (estimate / SE)
FDR (electrodes) + Holm (3 contrasts)
p < 0.05 uncorrected only

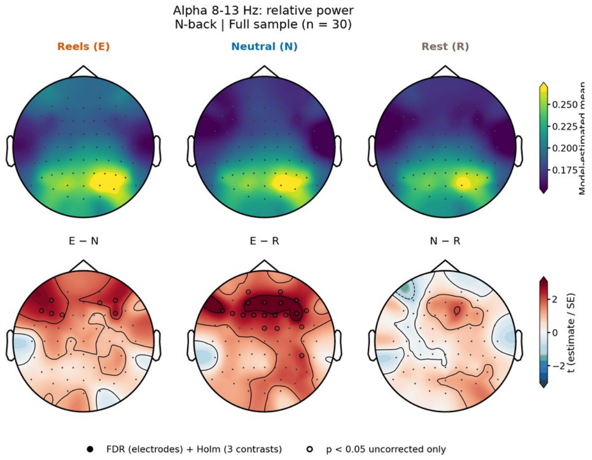
Alpha 8-13 Hz: relative power
N-back | Full sample (n = 30)
Reels (E)
Neutral (N)
Rest (R)
Model-estimated mean
E − N
E − R
N − R
t (estimate / SE)
FDR (electrodes) + Holm (3 contrasts)
p < 0.05 uncorrected only

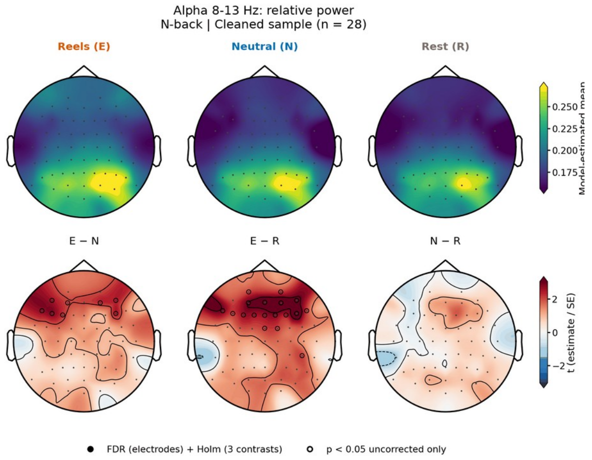
Alpha 8-13 Hz: relative power
N-back | Cleaned sample (n = 28)
Reels (E)
Neutral (N)
Rest (R)
Model-estimated mean
E − N
E − R
N − R
t (estimate / SE)
FDR (electrodes) + Holm (3 contrasts)
p < 0.05 uncorrected only

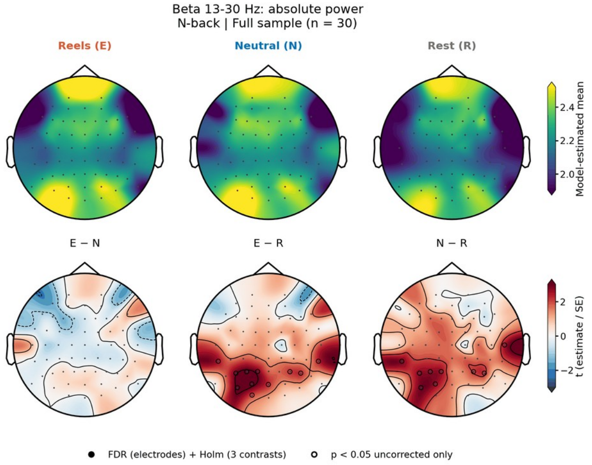
Beta 13-30 Hz: absolute power
N-back | Full sample (n = 30)
Reels (E)
Neutral (N)
Rest (R)
Model-estimated mean
E − N
E − R
N − R
t (estimate / SE)
FDR (electrodes) + Holm (3 contrasts)
p < 0.05 uncorrected only

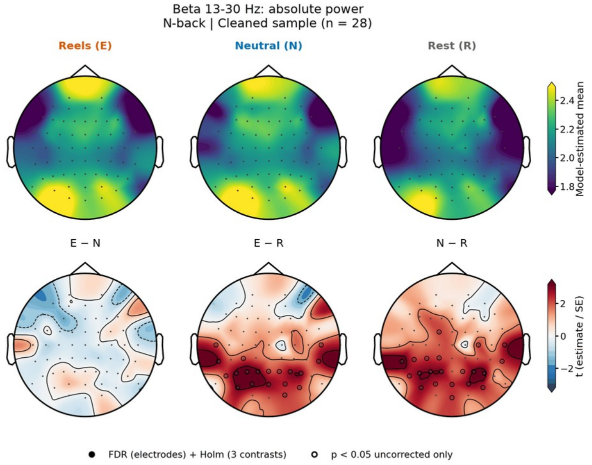
Beta 13-30 Hz: absolute power
N-back | Cleaned sample (n = 28)
Reels (E)
Neutral (N)
Rest (R)
Model-estimated mean
E − N
E − R
N − R
t (estimate / SE)
FDR (electrodes) + Holm (3 contrasts)
p < 0.05 uncorrected only

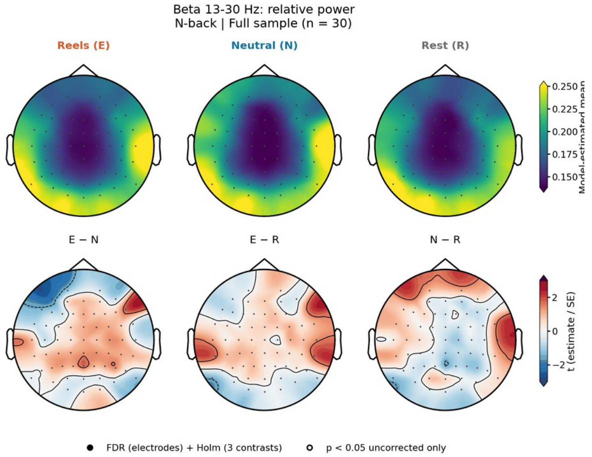
Beta 13-30 Hz: relative power
N-back | Full sample (n = 30)
Reels (E)
Neutral (N)
Rest (R)
E − N
E − R
N − R
t (estimate / SE)
FDR (electrodes) + Holm (3 contrasts)
p < 0.05 uncorrected only

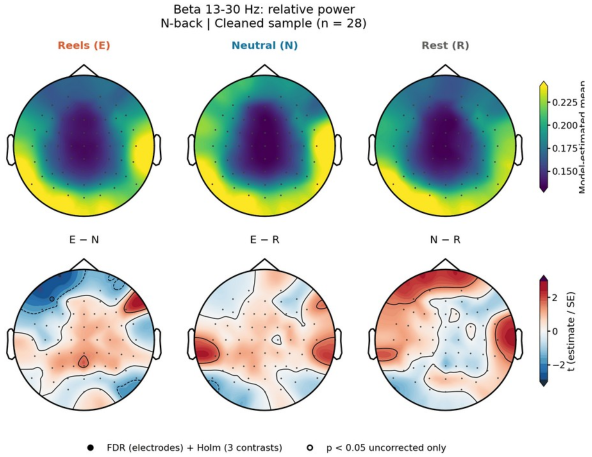
Beta 13-30 Hz: relative power
N-back | Cleaned sample (n = 28)
Reels (E)
Neutral (N)
Rest (R)
E − N
E − R
N − R
t (estimate / SE)
FDR (electrodes) + Holm (3 contrasts)
p < 0.05 uncorrected only

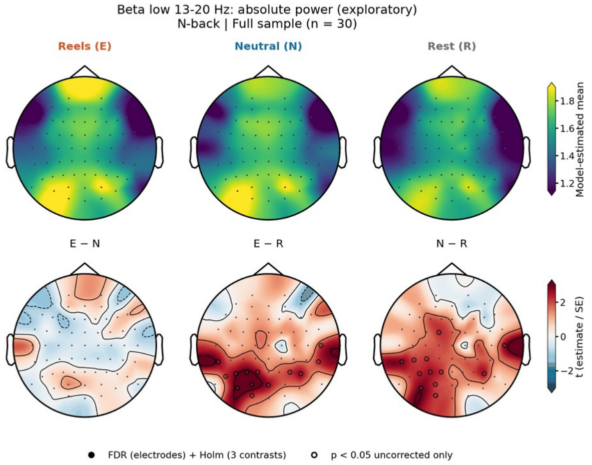
Beta low 13-20 Hz: absolute power (exploratory)
N-back | Full sample (n = 30)
Reels (E)
Neutral (N)
Rest (R)
Model-estimated mean
E − N
E − R
N − R
t (estimate / SE)
FDR (electrodes) + Holm (3 contrasts)
p < 0.05 uncorrected only

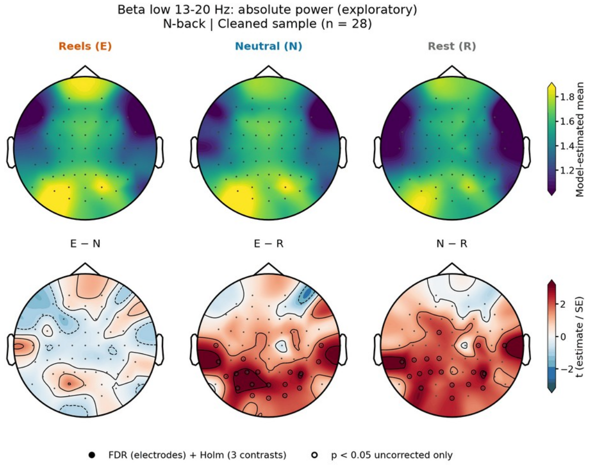
Beta low 13-20 Hz: absolute power (exploratory)
N-back | Cleaned sample (n = 28)
Reels (E)
Neutral (N)
Rest (R)
Model-estimated mean
E − N
E − R
N − R
t (estimate / SE)
FDR (electrodes) + Holm (3 contrasts)
p < 0.05 uncorrected only

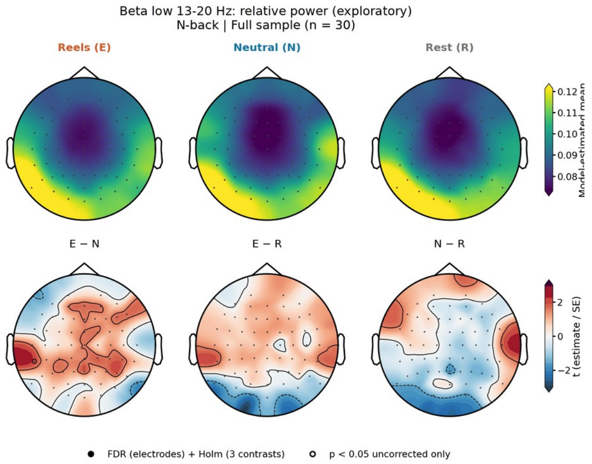
Beta low 13-20 Hz: relative power (exploratory)
N-back | Full sample (n = 30)
Reels (E)
Neutral (N)
Rest (R)
Model-estimated mean
0.12
0.11
0.10
0.09
0.08
E − N
E − R
N − R
t (estimate / SE)
2
0
−2
FDR (electrodes) + Holm (3 contrasts)
p < 0.05 uncorrected only

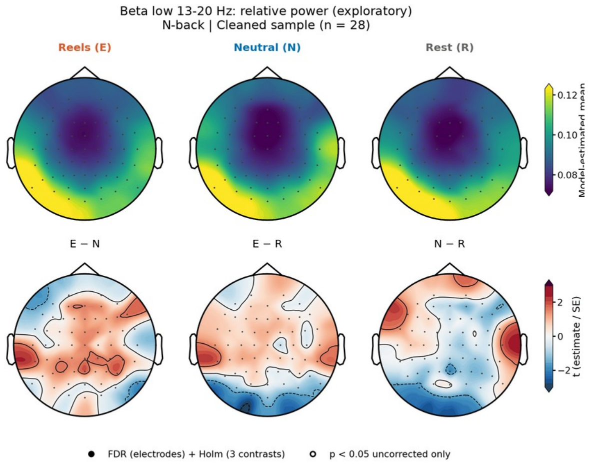
Beta low 13-20 Hz: relative power (exploratory)
N-back | Cleaned sample (n = 28)
Reels (E)
Neutral (N)
Rest (R)
Model-estimated mean
0.12
0.10
0.08
E − N
E − R
N − R
t (estimate / SE)
2
0
−2
FDR (electrodes) + Holm (3 contrasts)
p < 0.05 uncorrected only

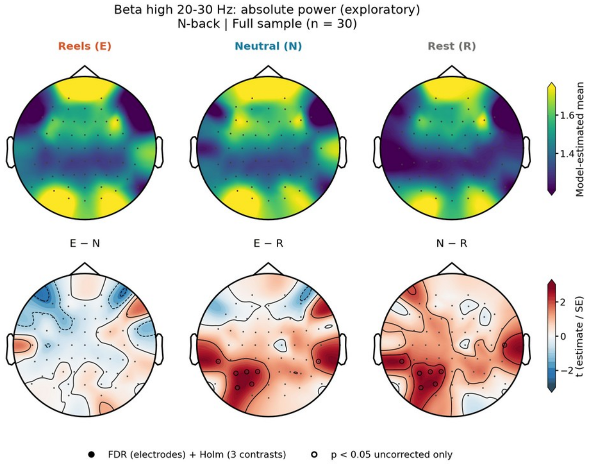
Beta high 20-30 Hz: absolute power (exploratory)
N-back | Full sample (n = 30)
Reels (E)
Neutral (N)
Rest (R)
Model-estimated mean
1.6
1.4
E − N
E − R
N − R
t (estimate / SE)
2
0
−2
FDR (electrodes) + Holm (3 contrasts)
p < 0.05 uncorrected only

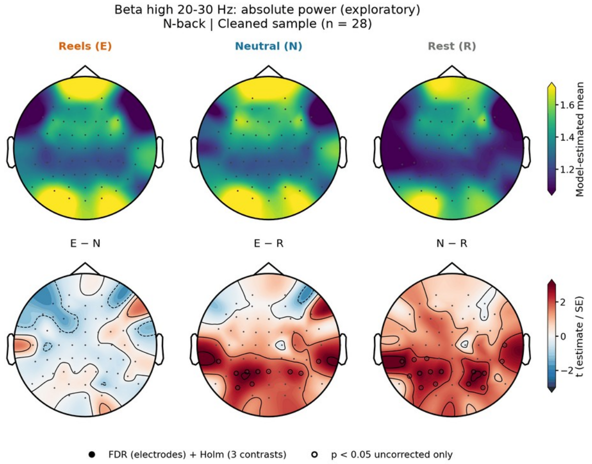
Beta high 20-30 Hz: absolute power (exploratory)
N-back | Cleaned sample (n = 28)
Reels (E)
Neutral (N)
Rest (R)
Model-estimated mean
1.6
1.4
1.2
E − N
E − R
N − R
t (estimate / SE)
2
0
−2
FDR (electrodes) + Holm (3 contrasts)
p < 0.05 uncorrected only

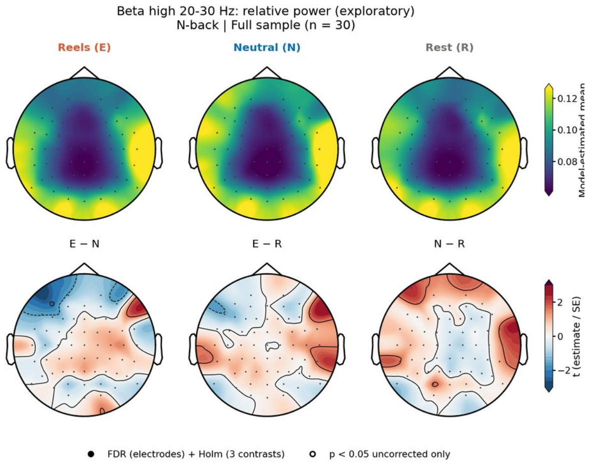
Beta high 20-30 Hz: relative power (exploratory)
N-back | Full sample (n = 30)
Reels (E)
Neutral (N)
Rest (R)
Model-estimated mean
0.12
0.10
0.08
E − N
E − R
N − R
t (estimate / SE)
2
0
−2
FDR (electrodes) + Holm (3 contrasts)
p < 0.05 uncorrected only

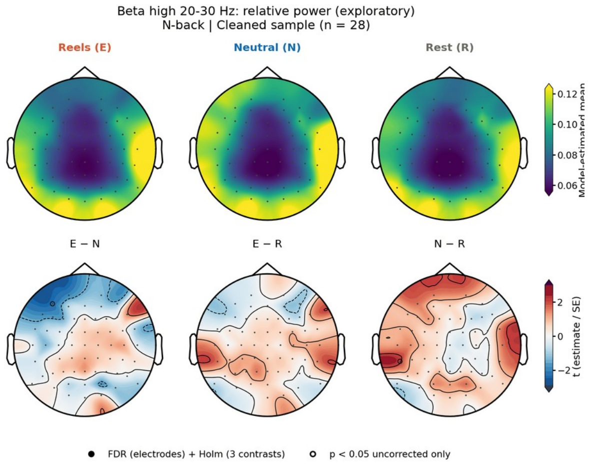
Beta high 20-30 Hz: relative power (exploratory)
N-back | Cleaned sample (n = 28)
Reels (E)
Neutral (N)
Rest (R)
Model-estimated mean
0.12
0.10
0.08
0.06
E − N
E − R
N − R
t (estimate / SE)
2
0
−2
FDR (electrodes) + Holm (3 contrasts)
p < 0.05 uncorrected only

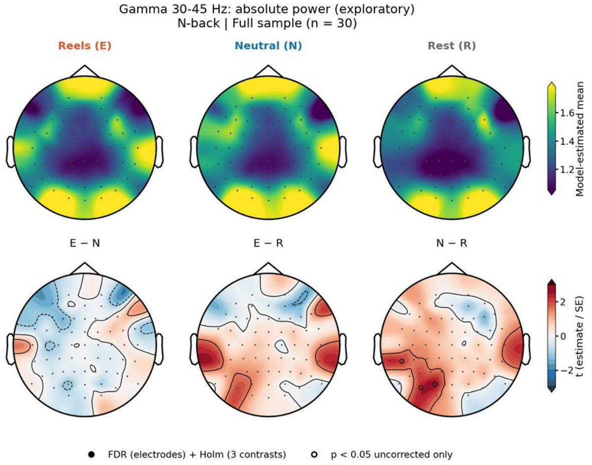
Gamma 30-45 Hz: absolute power (exploratory)
N-back | Full sample (n = 30)
Reels (E)
Neutral (N)
Rest (R)
Model-estimated mean
E − N
E − R
N − R
t (estimate / SE)
FDR (electrodes) + Holm (3 contrasts)
p < 0.05 uncorrected only

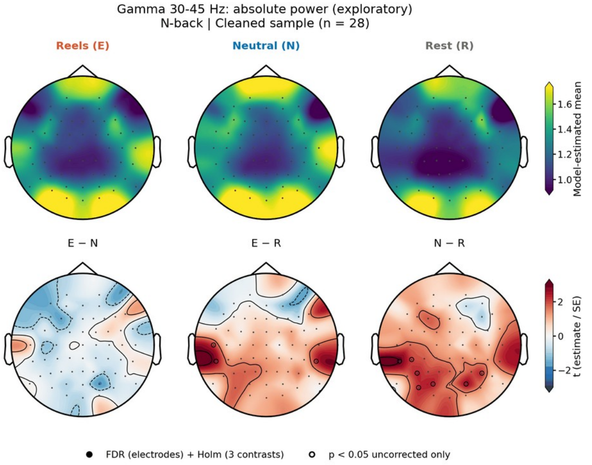
Gamma 30-45 Hz: absolute power (exploratory)
N-back | Cleaned sample (n = 28)
Reels (E)
Neutral (N)
Rest (R)
Model-estimated mean
E − N
E − R
N − R
t (estimate / SE)
FDR (electrodes) + Holm (3 contrasts)
p < 0.05 uncorrected only

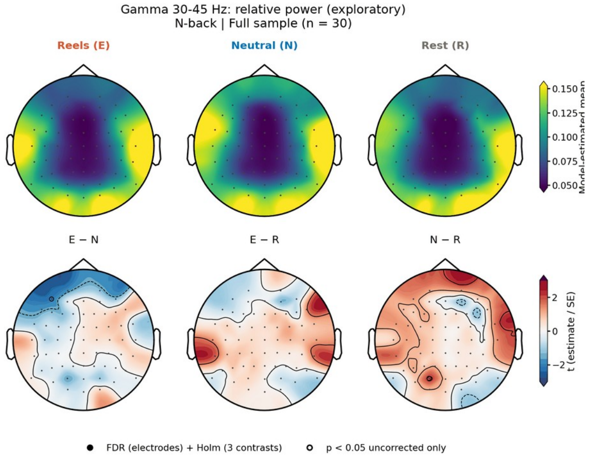
Gamma 30-45 Hz: relative power (exploratory)
N-back | Full sample (n = 30)
Reels (E)
Neutral (N)
Rest (R)
Model-estimated mean
E − N
E − R
N − R
t (estimate / SE)
FDR (electrodes) + Holm (3 contrasts)
p < 0.05 uncorrected only

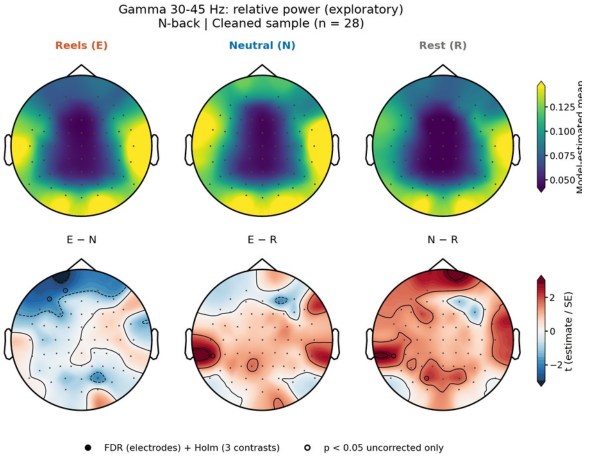
Gamma 30-45 Hz: relative power (exploratory)
N-back | Cleaned sample (n = 28)
Reels (E)
Neutral (N)
Rest (R)
Model-estimated mean
E − N
E − R
N − R
t (estimate / SE)
FDR (electrodes) + Holm (3 contrasts)
p < 0.05 uncorrected only

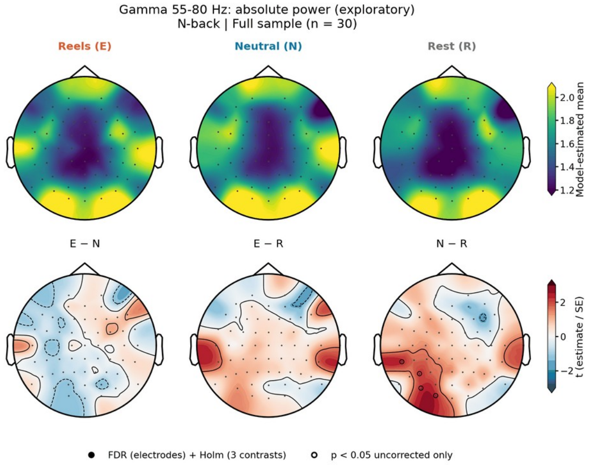
Gamma 55-80 Hz: absolute power (exploratory)
N-back | Full sample (n = 30)
Reels (E)
Neutral (N)
Rest (R)
Model-estimated mean
E − N
E − R
N − R
t (estimate / SE)
FDR (electrodes) + Holm (3 contrasts)
p < 0.05 uncorrected only

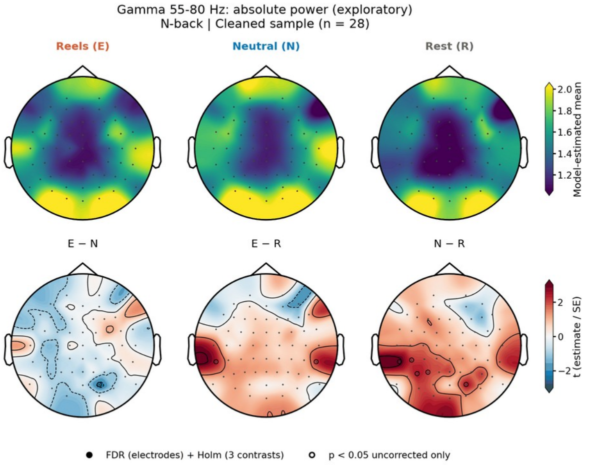
Gamma 55-80 Hz: absolute power (exploratory)
N-back | Cleaned sample (n = 28)
Reels (E)
Neutral (N)
Rest (R)
Model-estimated mean
E − N
E − R
N − R
t (estimate / SE)
FDR (electrodes) + Holm (3 contrasts)
p < 0.05 uncorrected only

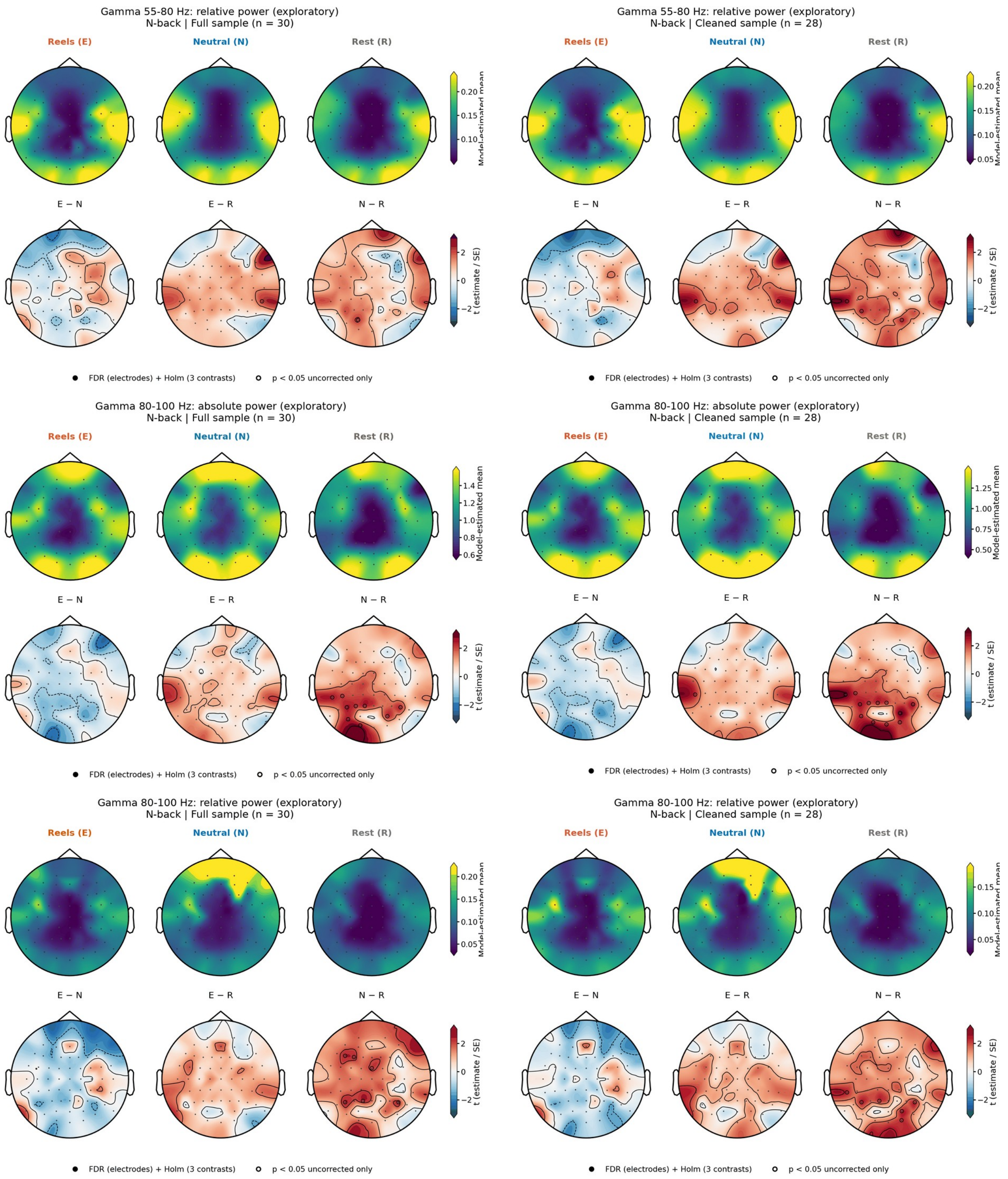


**Supplementary Fig. S18. Task EEG, N-back: electrode topographies for all frequency bands.** For each band and measure (absolute power, ln μV²; relative power, fraction of 1-45 Hz power), the left panel shows the full sample (N = 30) and the right panel the cleaned sample (N = 28). Upper row of each panel: model-estimated means after Reels (E), neutral images (N), and rest (R); lower row: *t* statistics (estimate/SE) of the E − N, E − R, and N − R contrasts. Filled circles, electrodes surviving FDR correction across electrodes and Holm correction across the three contrasts; open circles, uncorrected *p* < .05 only. Bands marked exploratory were not part of the primary analyses.

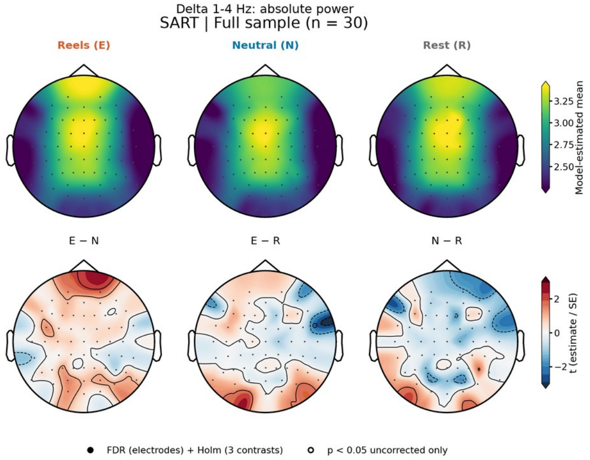
Delta 1-4 Hz: absolute power
SART | Full sample (n = 30)
Reels (E)
Neutral (N)
Rest (R)
Model-estimated mean
E − N
E − R
N − R
t (estimate / SE)
FDR (electrodes) + Holm (3 contrasts)
p < 0.05 uncorrected only

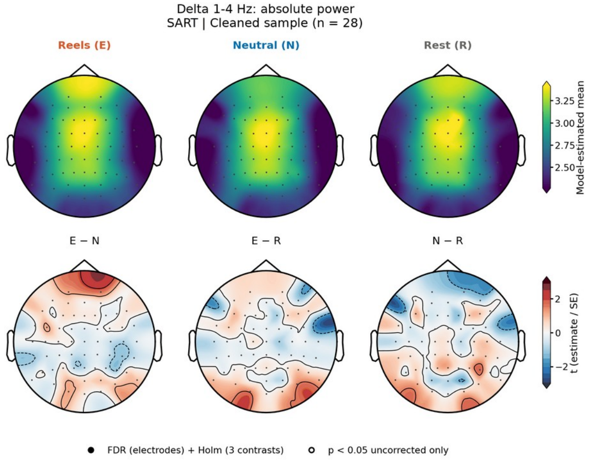
Delta 1-4 Hz: absolute power
SART | Cleaned sample (n = 28)
Reels (E)
Neutral (N)
Rest (R)
Model-estimated mean
E − N
E − R
N − R
t (estimate / SE)
FDR (electrodes) + Holm (3 contrasts)
p < 0.05 uncorrected only

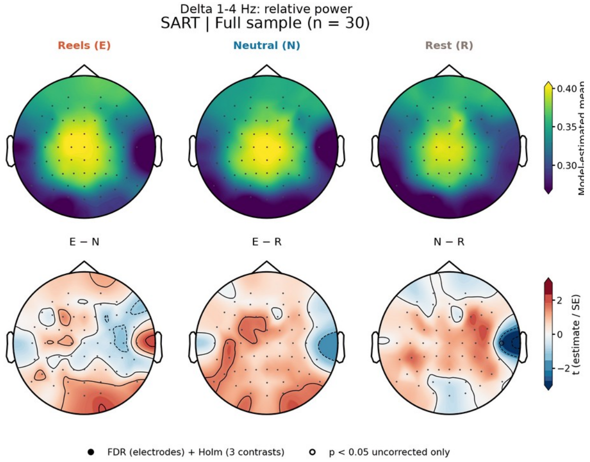
Delta 1-4 Hz: relative power
SART | Full sample (n = 30)
Reels (E)
Neutral (N)
Rest (R)
E − N
E − R
N − R
t (estimate / SE)
FDR (electrodes) + Holm (3 contrasts)
p < 0.05 uncorrected only

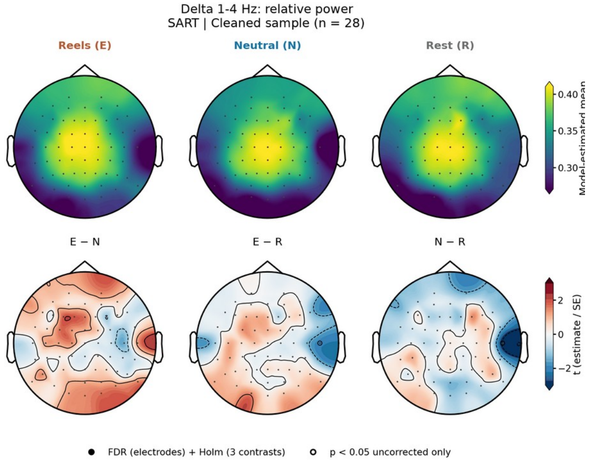
Delta 1-4 Hz: relative power
SART | Cleaned sample (n = 28)
Reels (E)
Neutral (N)
Rest (R)
E − N
E − R
N − R
t (estimate / SE)
FDR (electrodes) + Holm (3 contrasts)
p < 0.05 uncorrected only

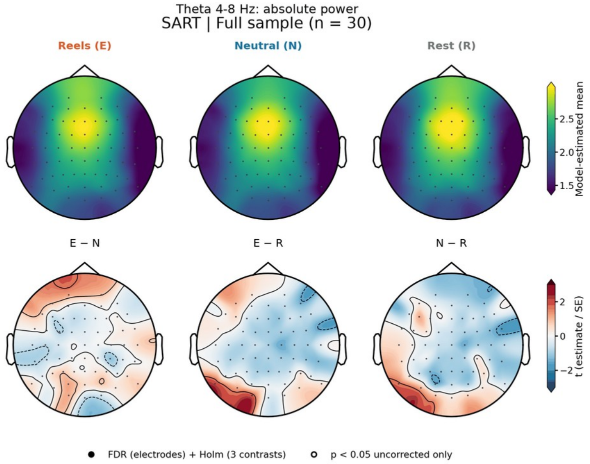
Theta 4-8 Hz: absolute power
SART | Full sample (n = 30)
Reels (E)
Neutral (N)
Rest (R)
Model-estimated mean
E − N
E − R
N − R
t (estimate / SE)
FDR (electrodes) + Holm (3 contrasts)
p < 0.05 uncorrected only

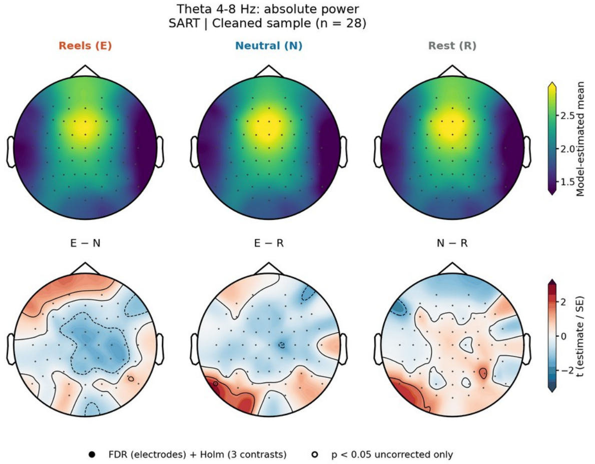
Theta 4-8 Hz: absolute power
SART | Cleaned sample (n = 28)
Reels (E)
Neutral (N)
Rest (R)
Model-estimated mean
E − N
E − R
N − R
t (estimate / SE)
FDR (electrodes) + Holm (3 contrasts)
p < 0.05 uncorrected only

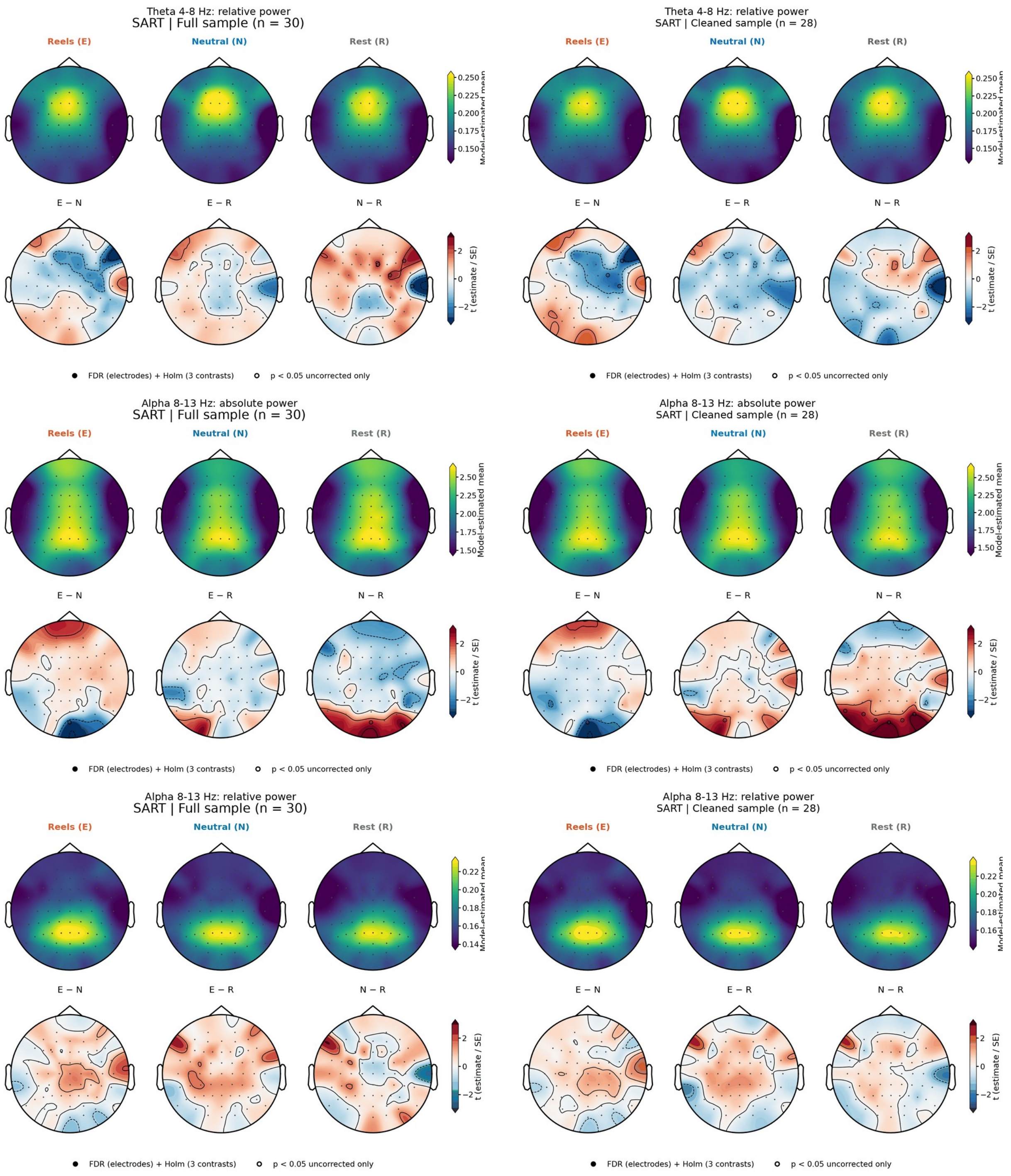
Theta 4-8 Hz: relative power
SART | Full sample (n = 30)
Reels (E)
Neutral (N)
Rest (R)
0.250
0.225
0.200
0.175
0.150
Model-estimated mean
E − N
E − R
N − R
2
0
−2
t (estimate / SE)
FDR (electrodes) + Holm (3 contrasts)
p < 0.05 uncorrected only
Theta 4-8 Hz: relative power
SART | Cleaned sample (n = 28)
Reels (E)
Neutral (N)
Rest (R)
0.250
0.225
0.200
0.175
0.150
Model-estimated mean
E − N
E − R
N − R
2
0
−2
t (estimate / SE)
FDR (electrodes) + Holm (3 contrasts)
p < 0.05 uncorrected only
Alpha 8-13 Hz: absolute power
SART | Full sample (n = 30)
Reels (E)
Neutral (N)
Rest (R)
2.50
2.25
2.00
1.75
1.50
Model-estimated mean
E − N
E − R
N − R
2
0
−2
t (estimate / SE)
FDR (electrodes) + Holm (3 contrasts)
p < 0.05 uncorrected only
Alpha 8-13 Hz: absolute power
SART | Cleaned sample (n = 28)
Reels (E)
Neutral (N)
Rest (R)
2.50
2.25
2.00
1.75
1.50
Model-estimated mean
E − N
E − R
N − R
2
0
−2
t (estimate / SE)
FDR (electrodes) + Holm (3 contrasts)
p < 0.05 uncorrected only
Alpha 8-13 Hz: relative power
SART | Full sample (n = 30)
Reels (E)
Neutral (N)
Rest (R)
0.22
0.20
0.18
0.16
0.14
Model-estimated mean
E − N
E − R
N − R
2
0
−2
t (estimate / SE)
FDR (electrodes) + Holm (3 contrasts)
p < 0.05 uncorrected only
Alpha 8-13 Hz: relative power
SART | Cleaned sample (n = 28)
Reels (E)
Neutral (N)
Rest (R)
0.22
0.20
0.18
0.16
Model-estimated mean
E − N
E − R
N − R
2
0
−2
t (estimate / SE)
FDR (electrodes) + Holm (3 contrasts)
p < 0.05 uncorrected only

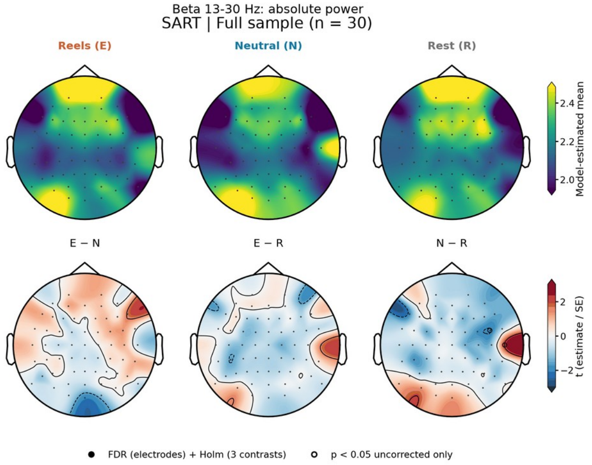
Beta 13-30 Hz: absolute power
SART | Full sample (n = 30)
Reels (E)
Neutral (N)
Rest (R)
Model-estimated mean
E − N
E − R
N − R
t (estimate / SE)
FDR (electrodes) + Holm (3 contrasts)
p < 0.05 uncorrected only

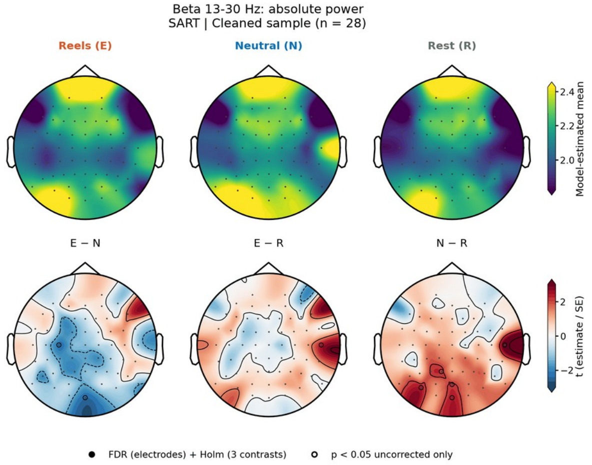
Beta 13-30 Hz: absolute power
SART | Cleaned sample (n = 28)
Reels (E)
Neutral (N)
Rest (R)
Model-estimated mean
E − N
E − R
N − R
t (estimate / SE)
FDR (electrodes) + Holm (3 contrasts)
p < 0.05 uncorrected only

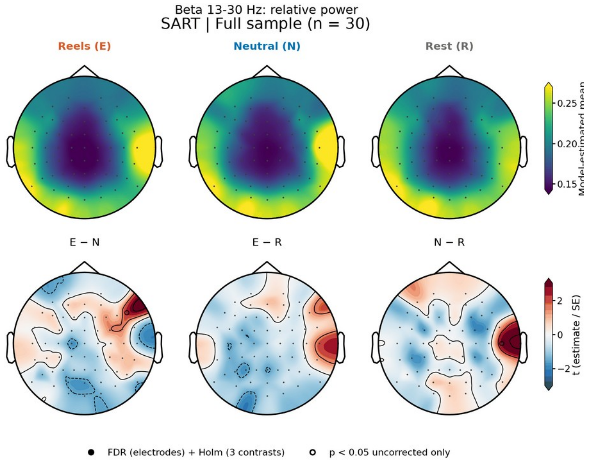
Beta 13-30 Hz: relative power
SART | Full sample (n = 30)
Reels (E)
Neutral (N)
Rest (R)
E − N
E − R
N − R
t (estimate / SE)
FDR (electrodes) + Holm (3 contrasts)
p < 0.05 uncorrected only

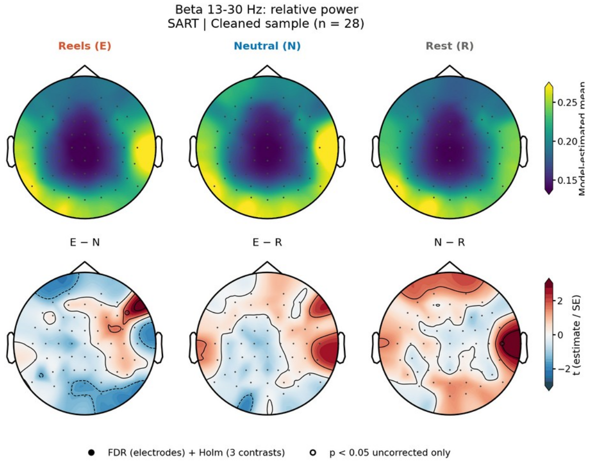
Beta 13-30 Hz: relative power
SART | Cleaned sample (n = 28)
Reels (E)
Neutral (N)
Rest (R)
E − N
E − R
N − R
t (estimate / SE)
FDR (electrodes) + Holm (3 contrasts)
p < 0.05 uncorrected only

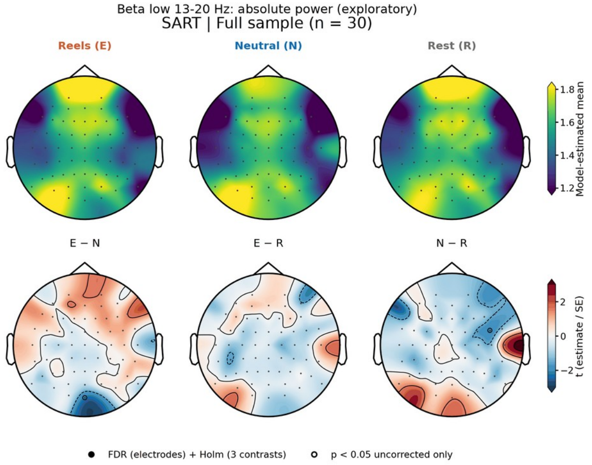
Beta low 13-20 Hz: absolute power (exploratory)
SART | Full sample (n = 30)
Reels (E)
Neutral (N)
Rest (R)
Model-estimated mean
E − N
E − R
N − R
t (estimate / SE)
FDR (electrodes) + Holm (3 contrasts)
p < 0.05 uncorrected only

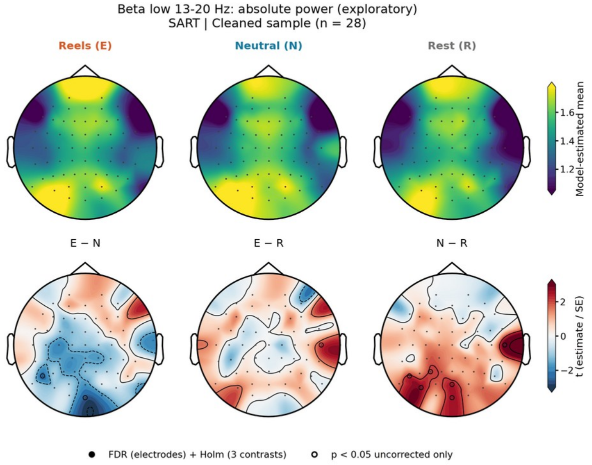
Beta low 13-20 Hz: absolute power (exploratory)
SART | Cleaned sample (n = 28)
Reels (E)
Neutral (N)
Rest (R)
Model-estimated mean
E − N
E − R
N − R
t (estimate / SE)
FDR (electrodes) + Holm (3 contrasts)
p < 0.05 uncorrected only

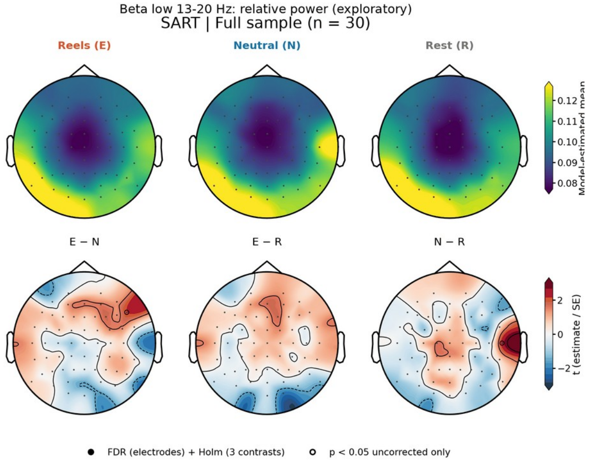
Beta low 13-20 Hz: relative power (exploratory)
SART | Full sample (n = 30)
Reels (E)
Neutral (N)
Rest (R)
Model-estimated mean
E − N
E − R
N − R
t (estimate / SE)
FDR (electrodes) + Holm (3 contrasts)
p < 0.05 uncorrected only

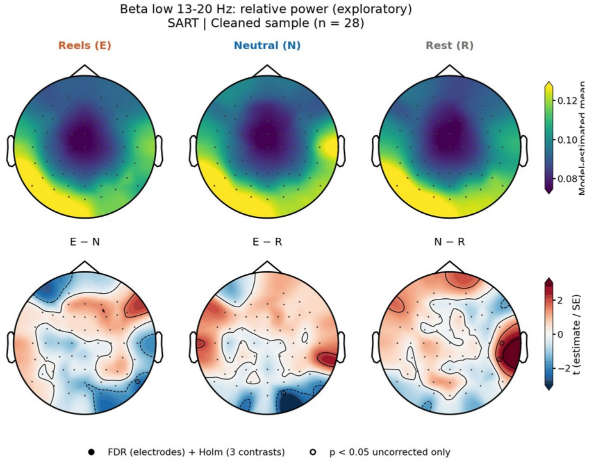
Beta low 13-20 Hz: relative power (exploratory)
SART | Cleaned sample (n = 28)
Reels (E)
Neutral (N)
Rest (R)
Model-estimated mean
E − N
E − R
N − R
t (estimate / SE)
FDR (electrodes) + Holm (3 contrasts)
p < 0.05 uncorrected only

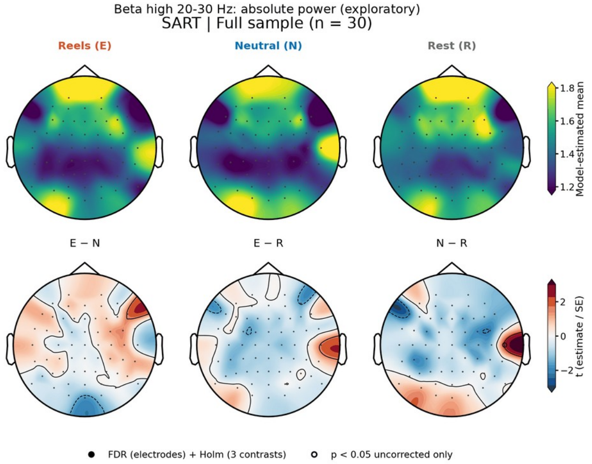
Beta high 20-30 Hz: absolute power (exploratory)
SART | Full sample (n = 30)
Reels (E)
Neutral (N)
Rest (R)
Model-estimated mean
E − N
E − R
N − R
t (estimate / SE)
FDR (electrodes) + Holm (3 contrasts)
p < 0.05 uncorrected only

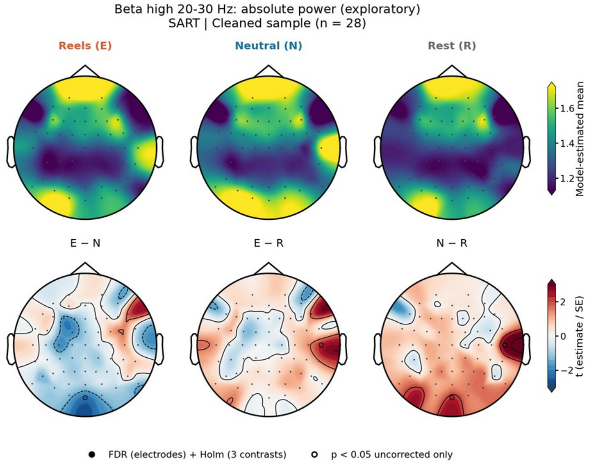
Beta high 20-30 Hz: absolute power (exploratory)
SART | Cleaned sample (n = 28)
Reels (E)
Neutral (N)
Rest (R)
Model-estimated mean
E − N
E − R
N − R
t (estimate / SE)
FDR (electrodes) + Holm (3 contrasts)
p < 0.05 uncorrected only

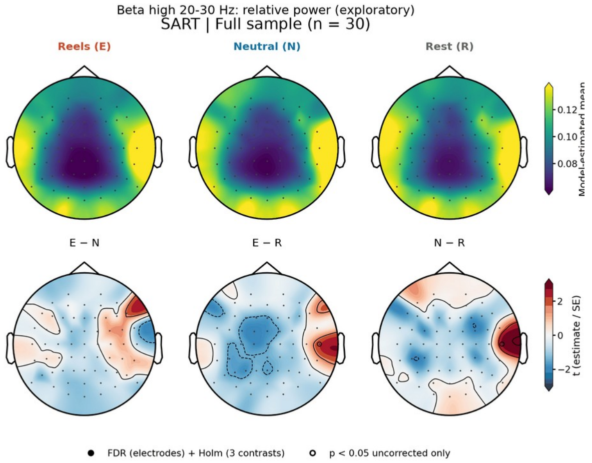
Beta high 20-30 Hz: relative power (exploratory)
SART | Full sample (n = 30)
Reels (E)
Neutral (N)
Rest (R)
Model-estimated mean
E − N
E − R
N − R
t (estimate / SE)
FDR (electrodes) + Holm (3 contrasts)
p < 0.05 uncorrected only

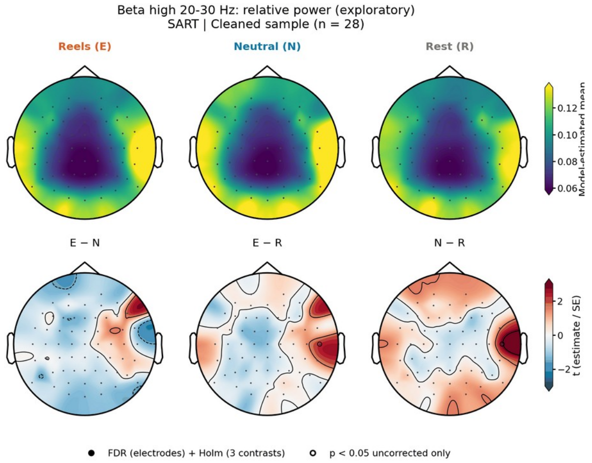
Beta high 20-30 Hz: relative power (exploratory)
SART | Cleaned sample (n = 28)
Reels (E)
Neutral (N)
Rest (R)
Model-estimated mean
E − N
E − R
N − R
t (estimate / SE)
FDR (electrodes) + Holm (3 contrasts)
p < 0.05 uncorrected only

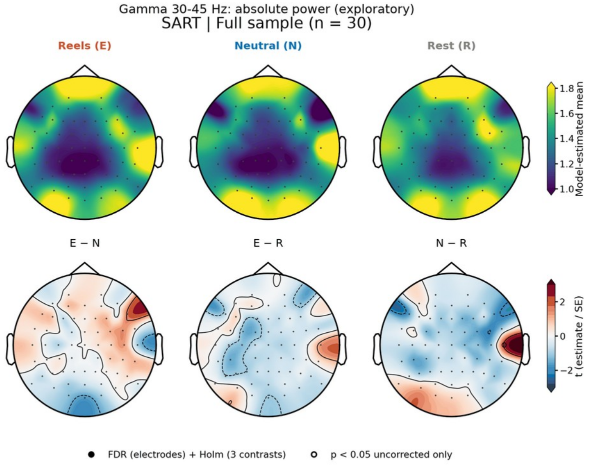
Gamma 30-45 Hz: absolute power (exploratory)
SART | Full sample (n = 30)
Reels (E)
Neutral (N)
Rest (R)
Model-estimated mean
E − N
E − R
N − R
t (estimate / SE)
FDR (electrodes) + Holm (3 contrasts)
p < 0.05 uncorrected only

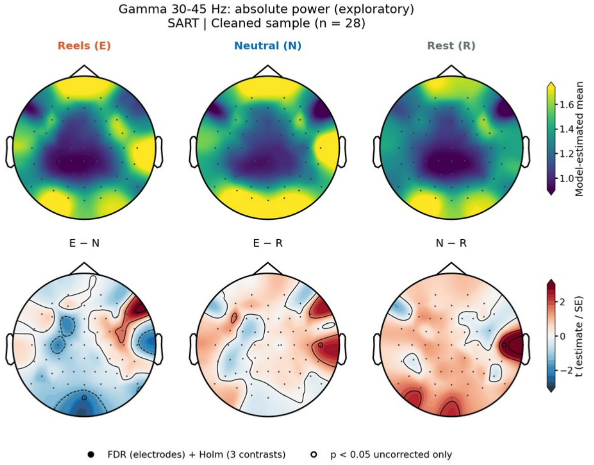
Gamma 30-45 Hz: absolute power (exploratory)
SART | Cleaned sample (n = 28)
Reels (E)
Neutral (N)
Rest (R)
Model-estimated mean
E − N
E − R
N − R
t (estimate / SE)
FDR (electrodes) + Holm (3 contrasts)
p < 0.05 uncorrected only

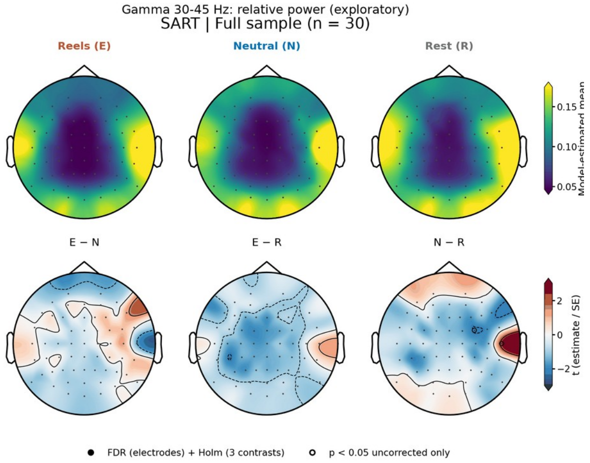
Gamma 30-45 Hz: relative power (exploratory)
SART | Full sample (n = 30)
Reels (E)
Neutral (N)
Rest (R)
E − N
E − R
N − R
t (estimate / SE)
FDR (electrodes) + Holm (3 contrasts)
p < 0.05 uncorrected only

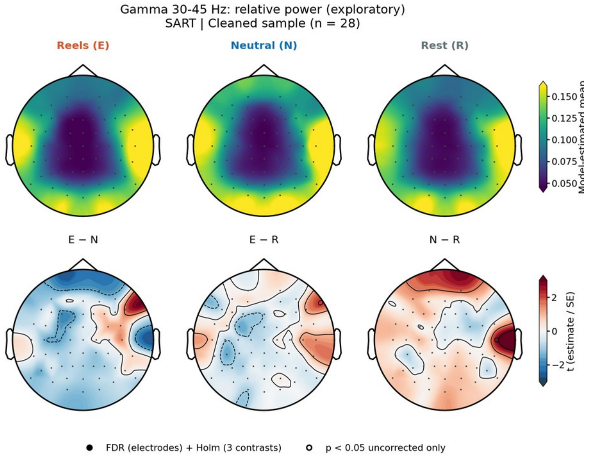
Gamma 30-45 Hz: relative power (exploratory)
SART | Cleaned sample (n = 28)
Reels (E)
Neutral (N)
Rest (R)
E − N
E − R
N − R
t (estimate / SE)
FDR (electrodes) + Holm (3 contrasts)
p < 0.05 uncorrected only

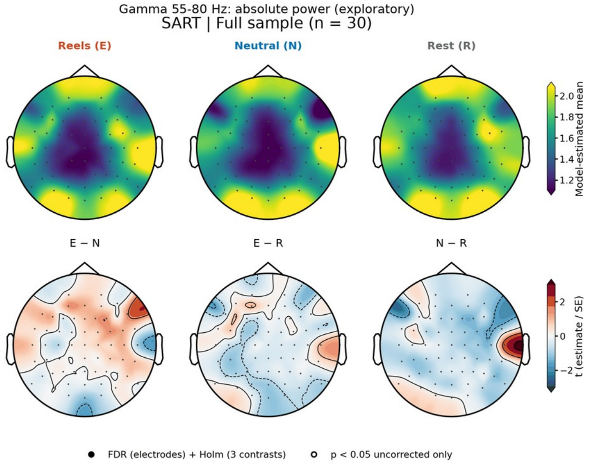
Gamma 55-80 Hz: absolute power (exploratory)
SART | Full sample (n = 30)
Reels (E)
Neutral (N)
Rest (R)
Model-estimated mean
E − N
E − R
N − R
t (estimate / SE)
FDR (electrodes) + Holm (3 contrasts)
p < 0.05 uncorrected only

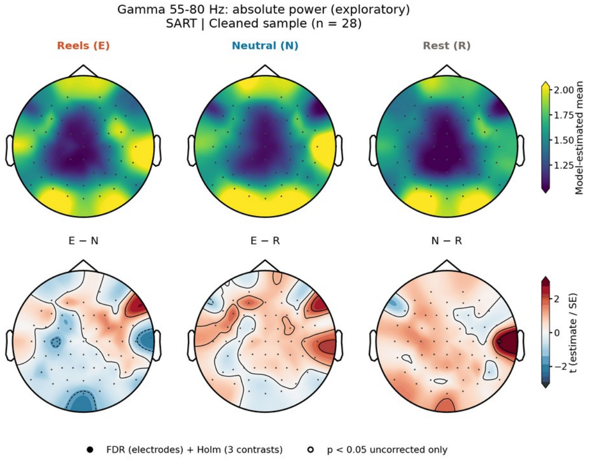
Gamma 55-80 Hz: absolute power (exploratory)
SART | Cleaned sample (n = 28)
Reels (E)
Neutral (N)
Rest (R)
Model-estimated mean
E − N
E − R
N − R
t (estimate / SE)
FDR (electrodes) + Holm (3 contrasts)
p < 0.05 uncorrected only

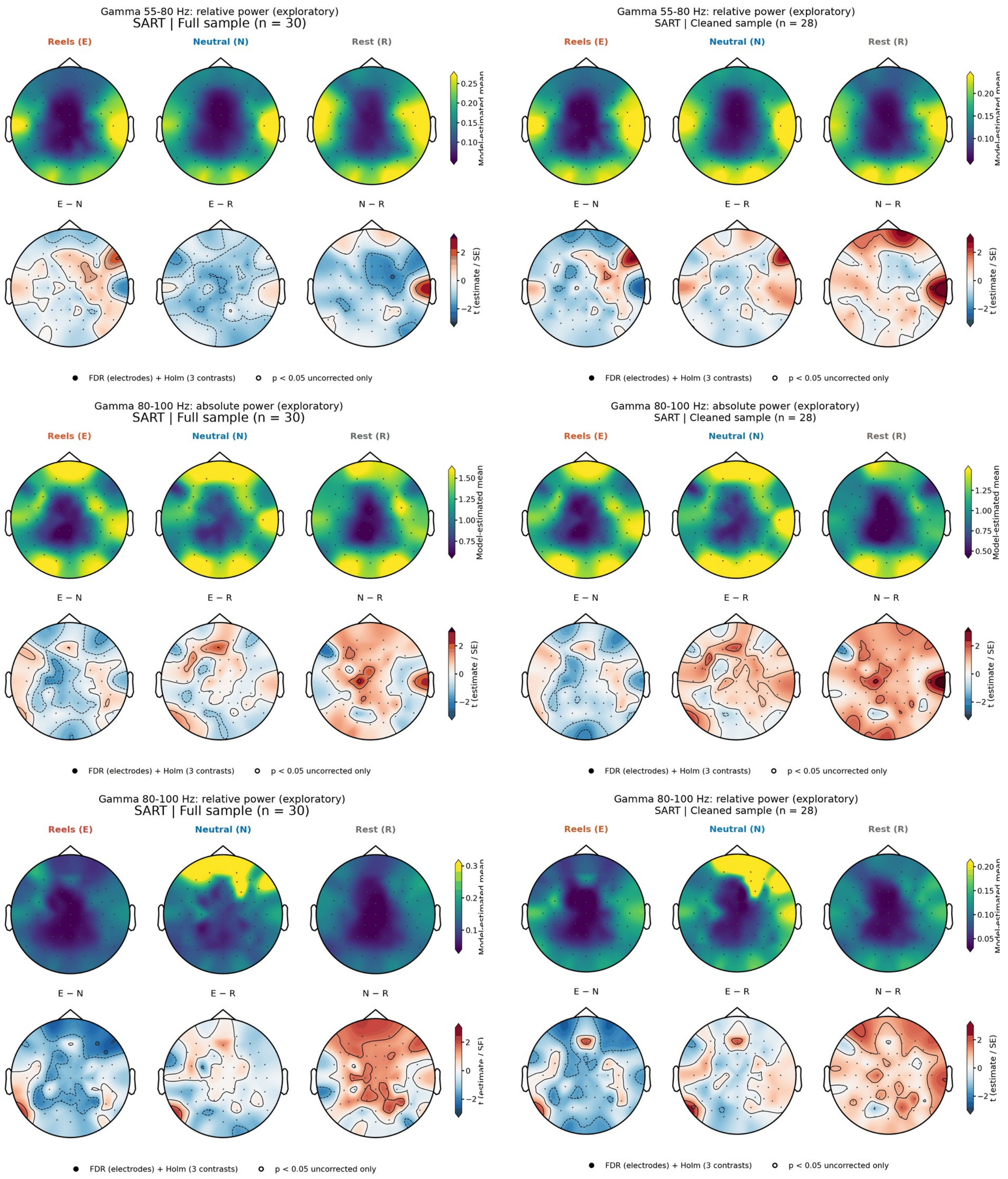


**Supplementary Fig. S19. Task EEG, SART: electrode topographies for all frequency bands.** For each band and measure (absolute power, ln μV²; relative power, fraction of 1-45 Hz power), the left panel shows the full sample (N = 30) and the right panel the cleaned sample (N = 28). Upper row of each panel: model-estimated means after Reels (E), neutral images (N), and rest (R); lower row: *t* statistics (estimate/SE) of the E − N, E − R, and N − R contrasts. Filled circles, electrodes surviving FDR correction across electrodes and Holm correction across the three contrasts; open circles, uncorrected *p* < .05 only. Bands marked exploratory were not part of the primary analyses.

**a**

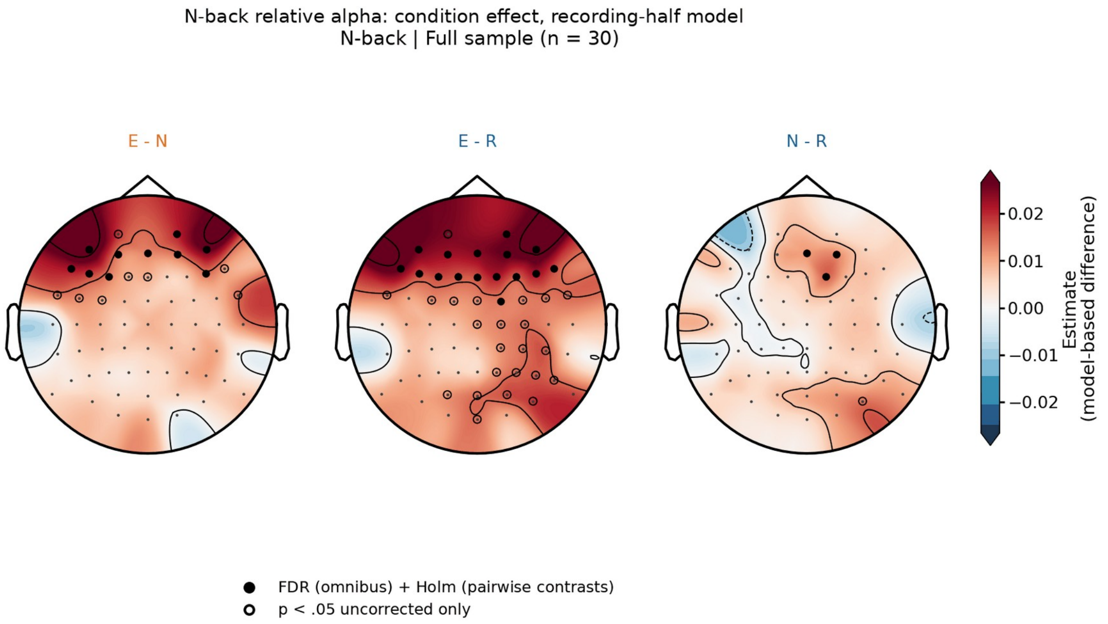


**b**

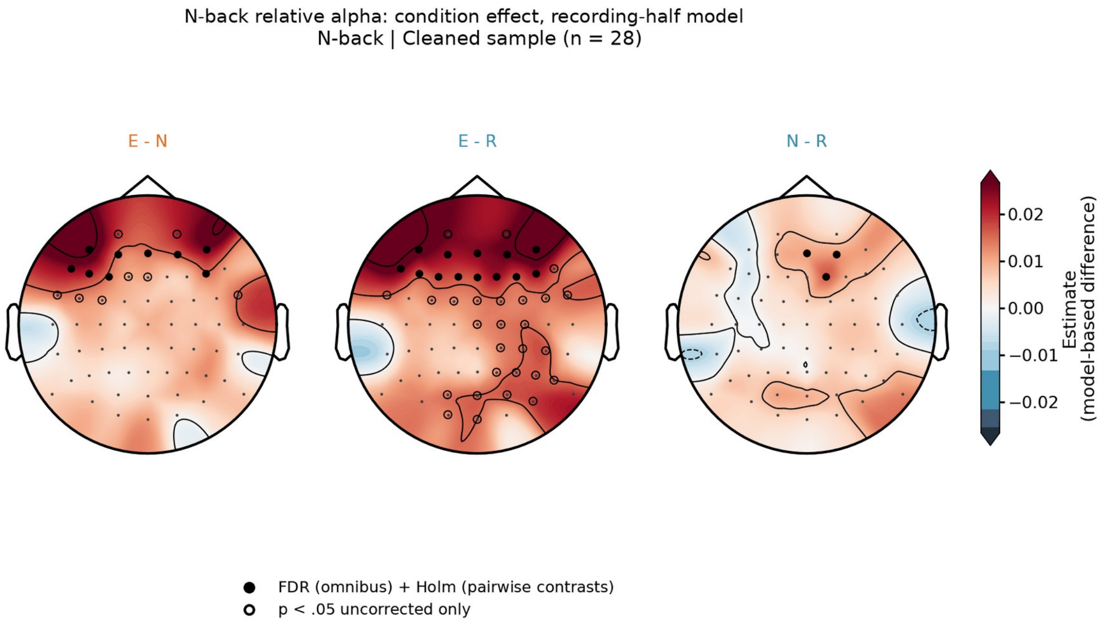


**Supplementary Fig. S20. N-back relative alpha in the condition × recording-half model.** Electrode-wise E − N, E − R, and N − R contrasts in the full (a) and cleaned (b) samples. Filled circles, electrodes whose omnibus test survived FDR correction and whose contrast survived Holm correction; open circles, uncorrected $p < .05$ only. Reels exceeded neutral images at 10 (full sample) and 9 (cleaned sample) frontal electrodes after correction. Because this model enters two observations per recording without modelling their dependence, it is shown as supportive evidence only.

**a**

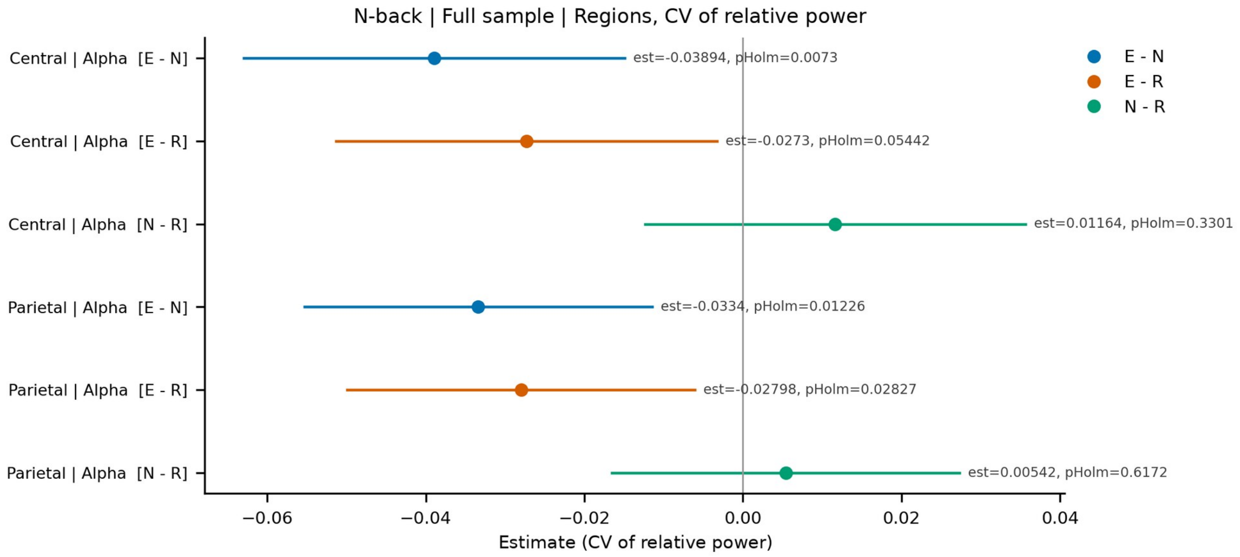


**b**

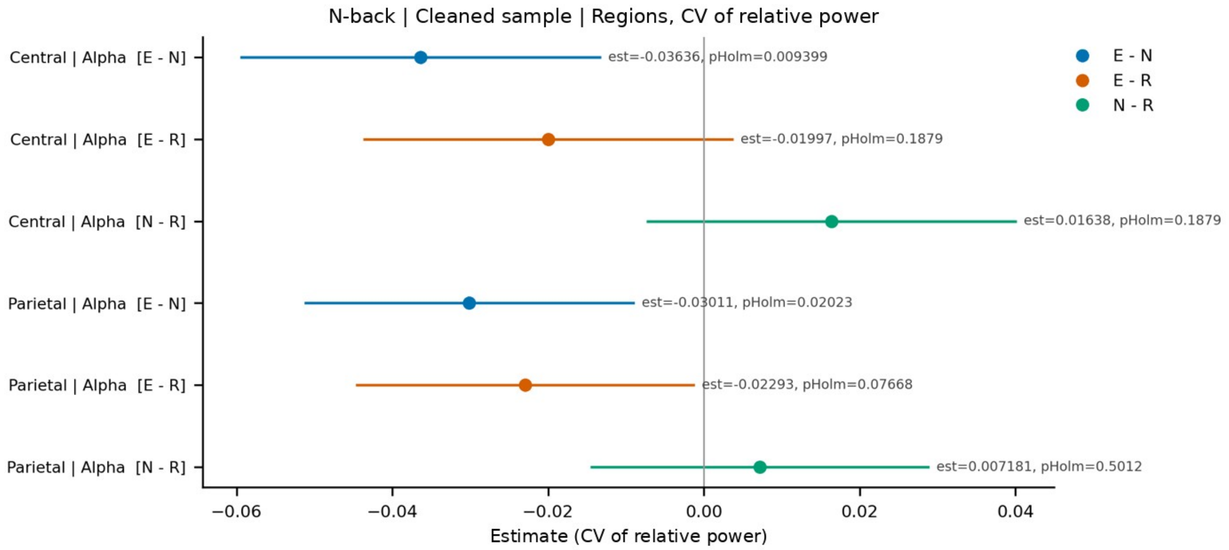


**Supplementary Fig. S21. Moment-to-moment variability of N-back relative alpha.** Coefficient of variation (CV) of relative alpha across 4-s windows in the central and parietal regions: E − N, E − R, and N − R contrasts with 95% CIs and Holm-corrected *p* values in the full (a) and cleaned (b) samples. Variability was lower after Reels than after neutral images in both regions and both samples.

**a**

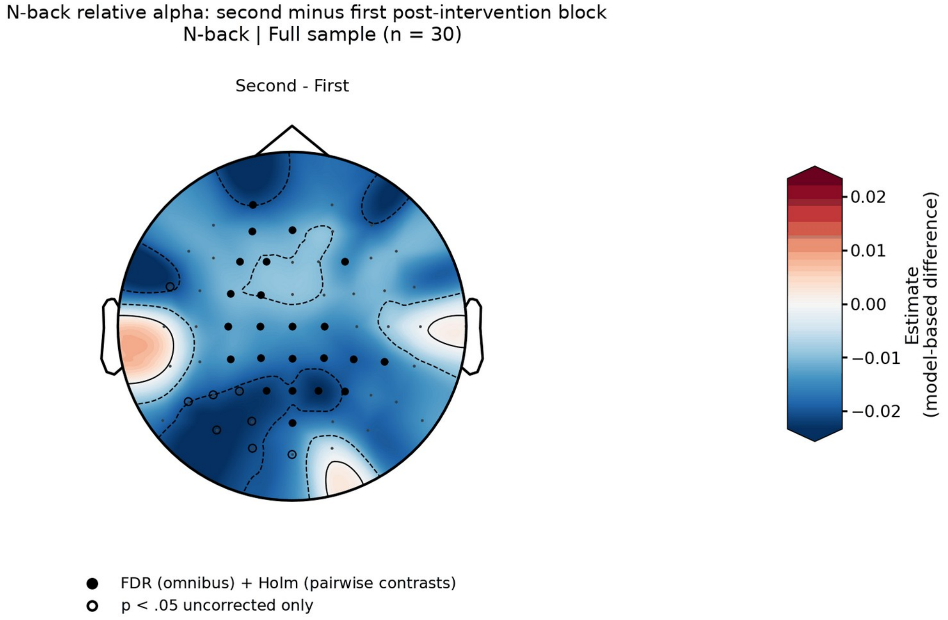


**b**

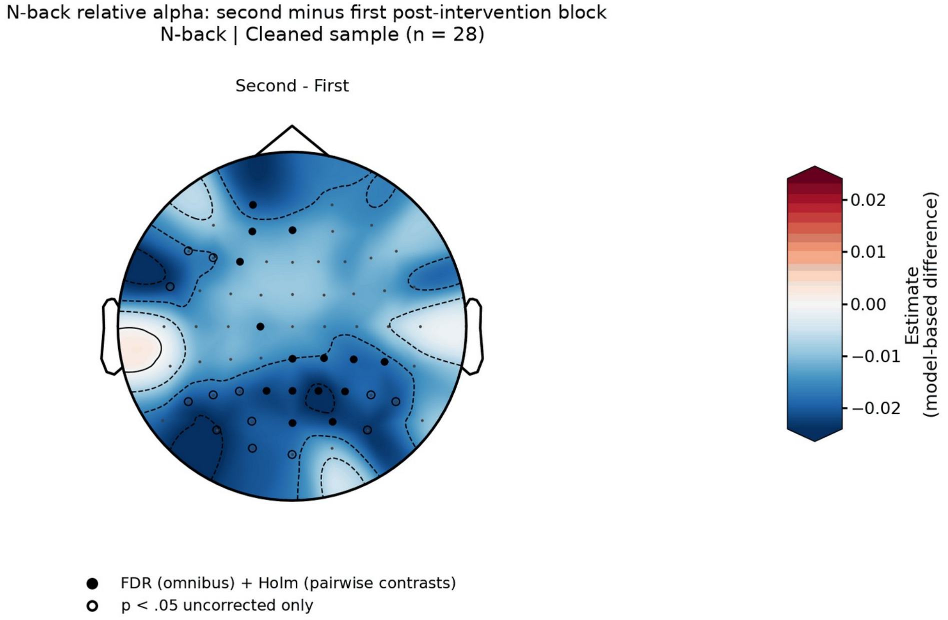


**Supplementary Fig. S22. Change in N-back relative alpha between the post-intervention blocks.** Electrode-wise difference between the second and the first post-intervention block (intervention-position effect), averaged over conditions, in the full (a) and cleaned (b) samples. Filled circles, electrodes surviving correction; open circles, uncorrected $p < .05$ only. Relative alpha decreased at 33 electrodes and globally in both samples, with no condition × position interaction.

**a**

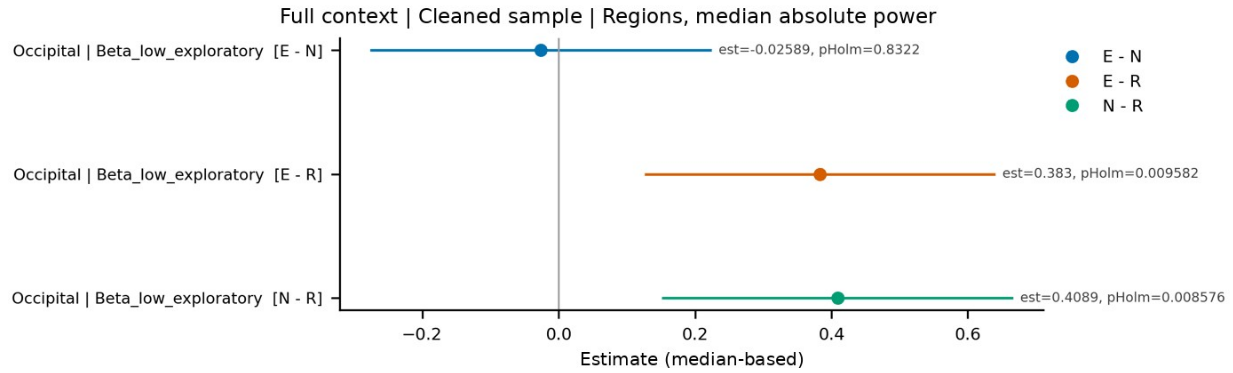
Full context | Cleaned sample | Regions, median absolute power
Occipital | Beta_low_exploratory [E - N]
Occipital | Beta_low_exploratory [E - R]
Occipital | Beta_low_exploratory [N - R]
est=-0.02589, pHolm=0.8322
est=0.383, pHolm=0.009582
est=0.4089, pHolm=0.008576
E - N
E - R
N - R
−0.2
0.0
0.2
0.4
0.6
Estimate (median-based)

**b**

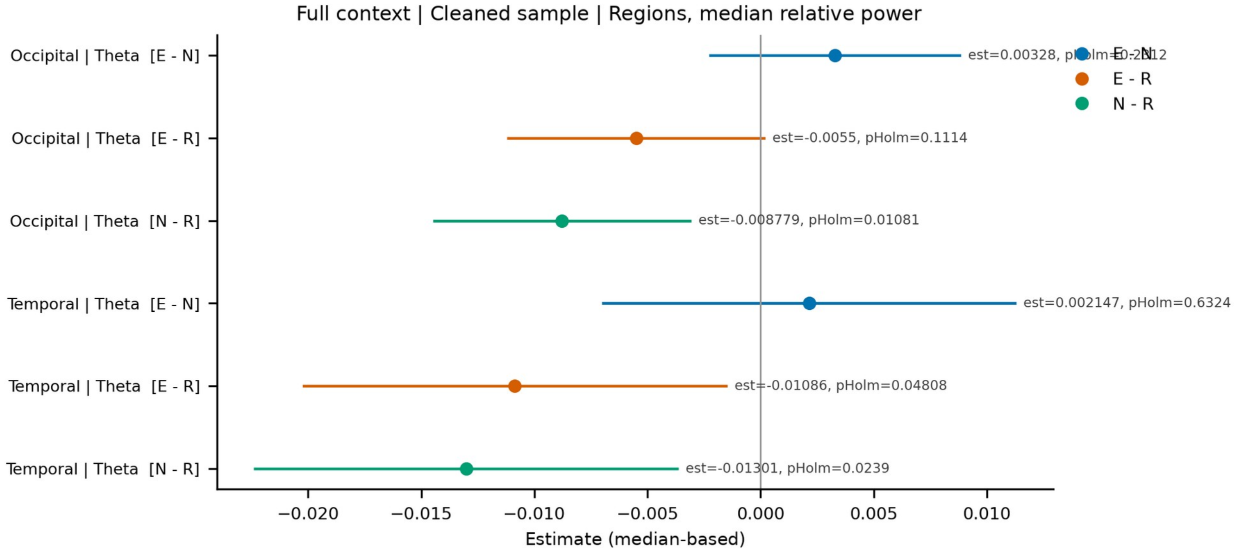
Full context | Cleaned sample | Regions, median relative power
Occipital | Theta [E - N]
Occipital | Theta [E - R]
Occipital | Theta [N - R]
Temporal | Theta [E - N]
Temporal | Theta [E - R]
Temporal | Theta [N - R]
est=-0.0055, pHolm=0.1114
est=-0.008779, pHolm=0.01081
est=0.002147, pHolm=0.6324
est=-0.01086, pHolm=0.04808
est=-0.01301, pHolm=0.0239
E - N
E - R
N - R
−0.020
−0.015
−0.010
−0.005
0.000
0.005
0.010
Estimate (median-based)

**c**

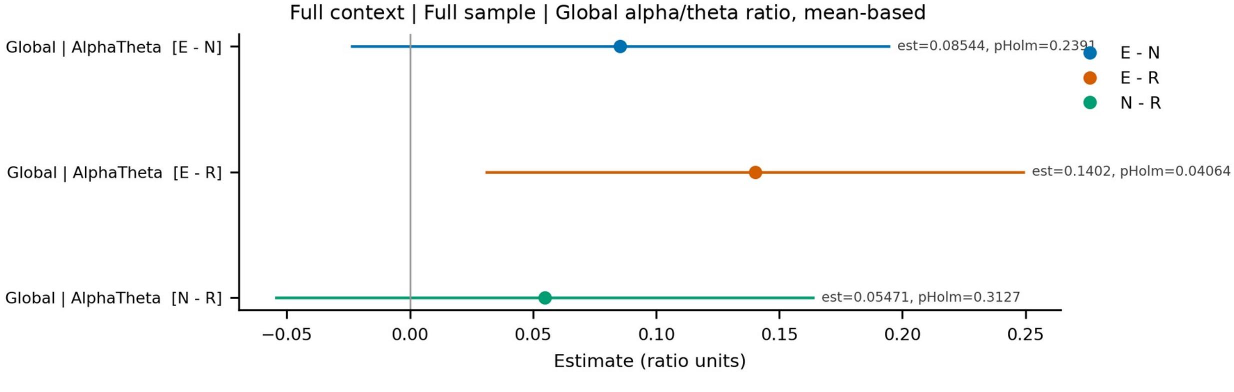
Full context | Full sample | Global alpha/theta ratio, mean-based
Global | AlphaTheta [E - N]
Global | AlphaTheta [E - R]
Global | AlphaTheta [N - R]
est=0.1402, pHolm=0.04064
est=0.05471, pHolm=0.3127
E - N
E - R
N - R
−0.05
0.00
0.05
0.10
0.15
0.20
0.25
Estimate (ratio units)

**d**

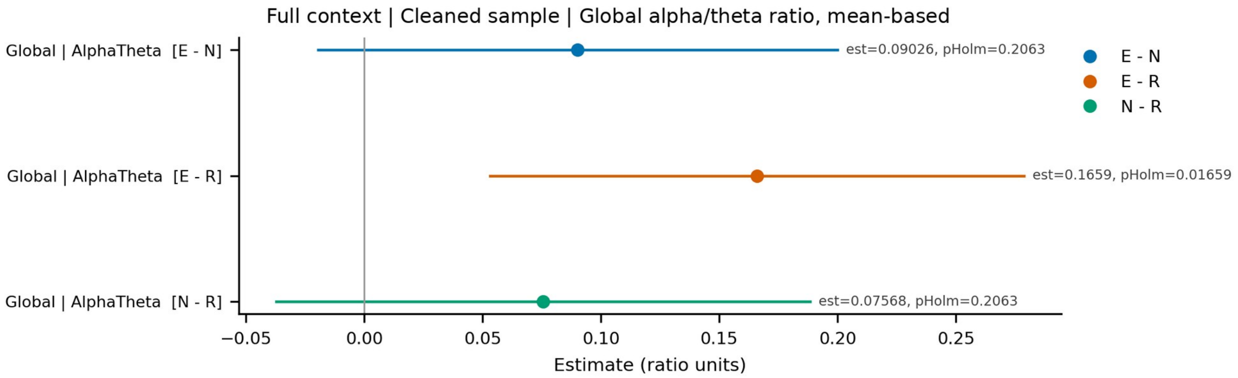


**e**

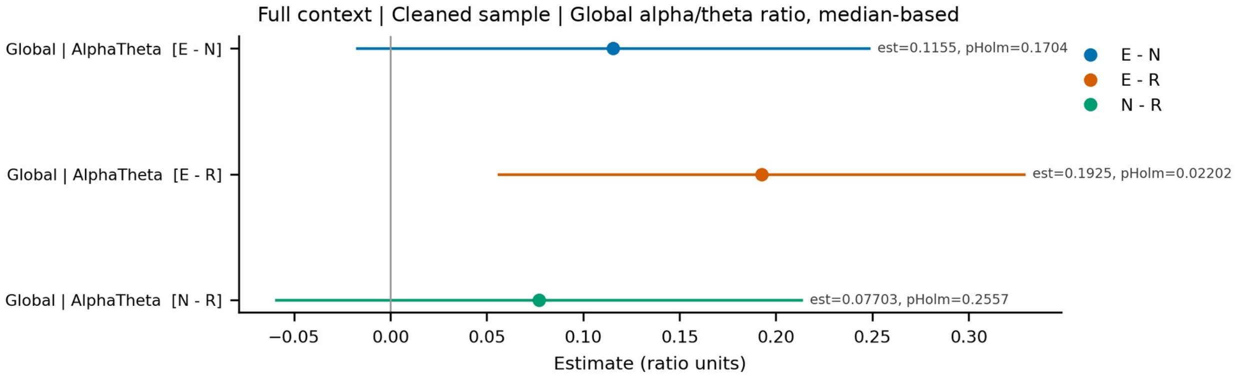


**Supplementary Fig. S23. Follow-up models, Full context.** Median-based regional power in the cleaned sample (a, b) and the global alpha/theta ratio, mean-based in the full (c) and cleaned (d) samples and median-based in the cleaned sample (e). Each row shows a contrast (E − N, E − R, or N − R; for position effects, second minus first post-intervention block) with its 95% CI; labels give the estimate and the Holm-corrected *p*. Only units whose omnibus test survived FDR correction are shown. Beta_low, Beta_high, Gamma_low, and Gamma_high1 denote the exploratory 13-20, 20-30, 30-45, and 55-80 Hz bands.

**a**

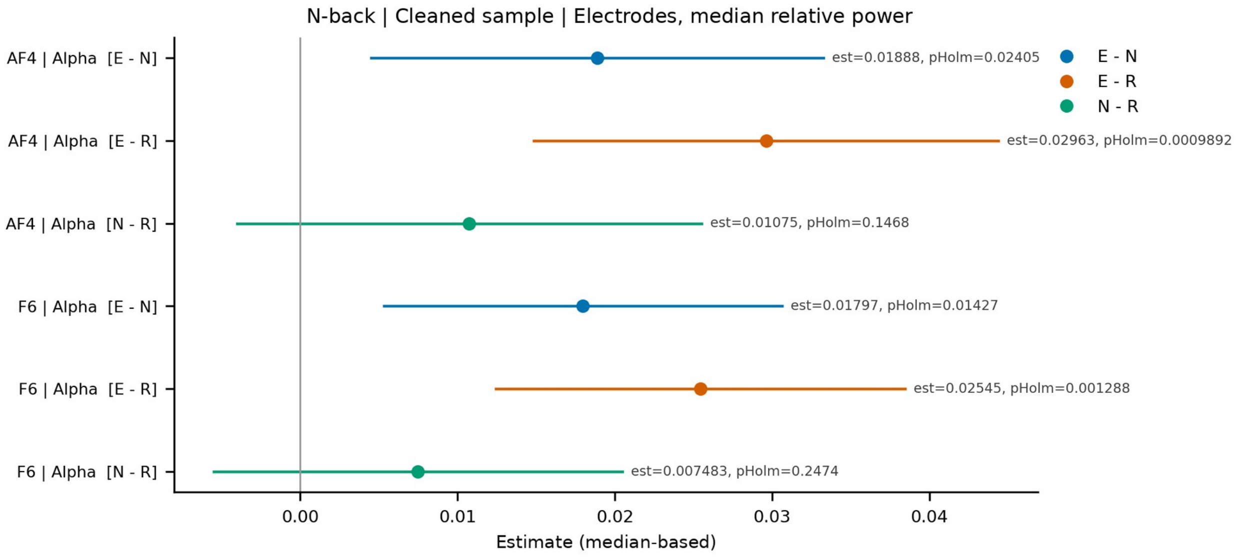
N-back | Cleaned sample | Electrodes, median relative power
AF4 | Alpha [E - N]
AF4 | Alpha [E - R]
AF4 | Alpha [N - R]
F6 | Alpha [E - N]
F6 | Alpha [E - R]
F6 | Alpha [N - R]
est=0.01888, pHolm=0.02405
est=0.02963, pHolm=0.0009892
est=0.01075, pHolm=0.1468
est=0.01797, pHolm=0.01427
est=0.02545, pHolm=0.001288
est=0.007483, pHolm=0.2474
E - N
E - R
N - R
0.00
0.01
0.02
0.03
0.04
Estimate (median-based)

**b**

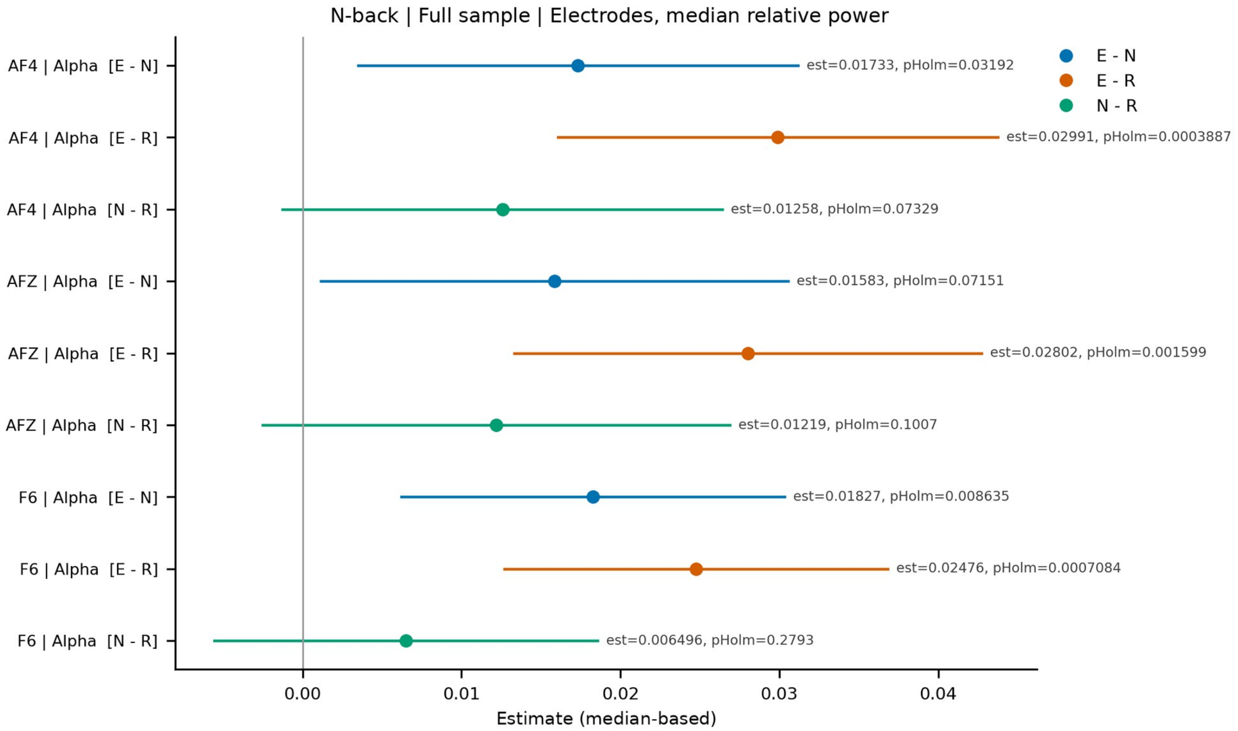
N-back | Full sample | Electrodes, median relative power
AF4 | Alpha [E - N]
AF4 | Alpha [E - R]
AF4 | Alpha [N - R]
AFZ | Alpha [E - N]
AFZ | Alpha [E - R]
AFZ | Alpha [N - R]
F6 | Alpha [E - N]
F6 | Alpha [E - R]
F6 | Alpha [N - R]
est=0.01733, pHolm=0.03192
est=0.02991, pHolm=0.0003887
est=0.01258, pHolm=0.07329
est=0.01583, pHolm=0.07151
est=0.02802, pHolm=0.001599
est=0.01219, pHolm=0.1007
est=0.01827, pHolm=0.008635
est=0.02476, pHolm=0.0007084
est=0.006496, pHolm=0.2793
E - N
E - R
N - R
0.00
0.01
0.02
0.03
0.04
Estimate (median-based)

**c**

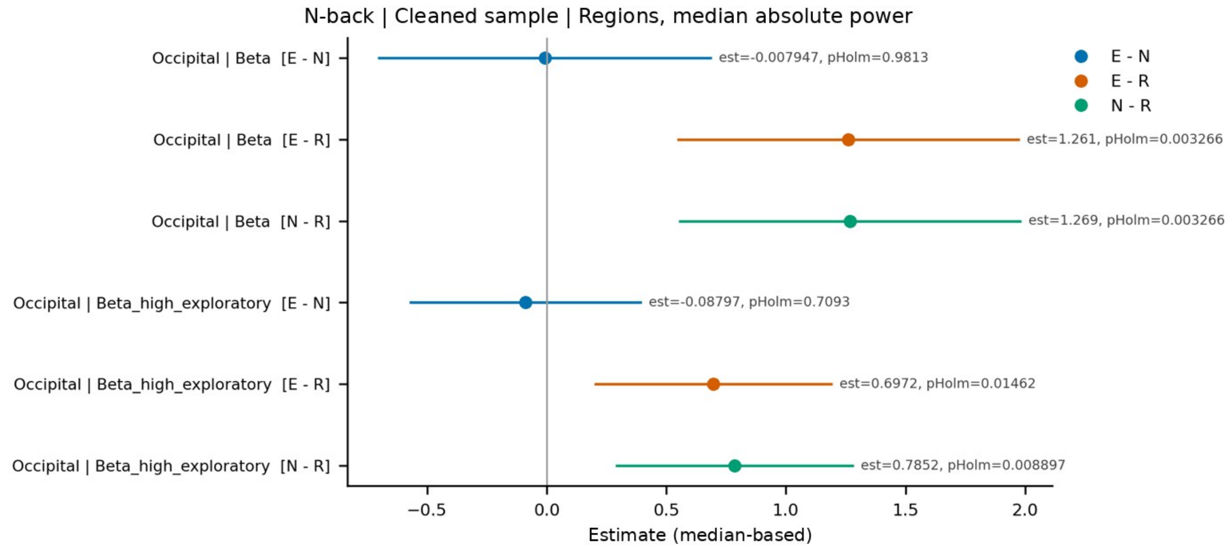
N-back | Cleaned sample | Regions, median absolute power
Occipital | Beta [E - N]
Occipital | Beta [E - R]
Occipital | Beta [N - R]
Occipital | Beta_high_exploratory [E - N]
Occipital | Beta_high_exploratory [E - R]
Occipital | Beta_high_exploratory [N - R]
est=-0.007947, pHolm=0.9813
est=1.261, pHolm=0.003266
est=1.269, pHolm=0.003266
est=-0.08797, pHolm=0.7093
est=0.6972, pHolm=0.01462
est=0.7852, pHolm=0.008897
E - N
E - R
N - R
−0.5
0.0
0.5
1.0
1.5
2.0
Estimate (median-based)

**d**

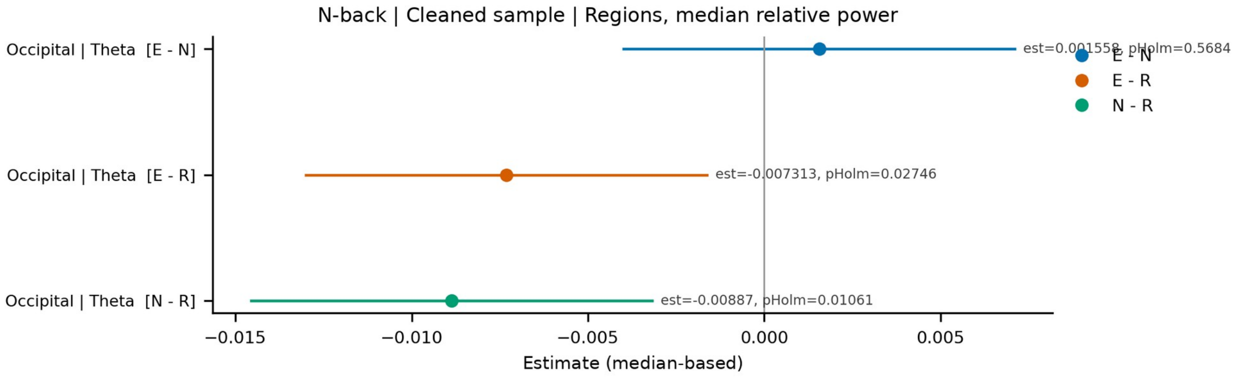
N-back | Cleaned sample | Regions, median relative power
Occipital | Theta [E - N]
Occipital | Theta [E - R]
Occipital | Theta [N - R]
est=0.001558, pHolm=0.5684
est=-0.007313, pHolm=0.02746
est=-0.00887, pHolm=0.01061
E - N
E - R
N - R
−0.015
−0.010
−0.005
0.000
0.005
Estimate (median-based)

**e**

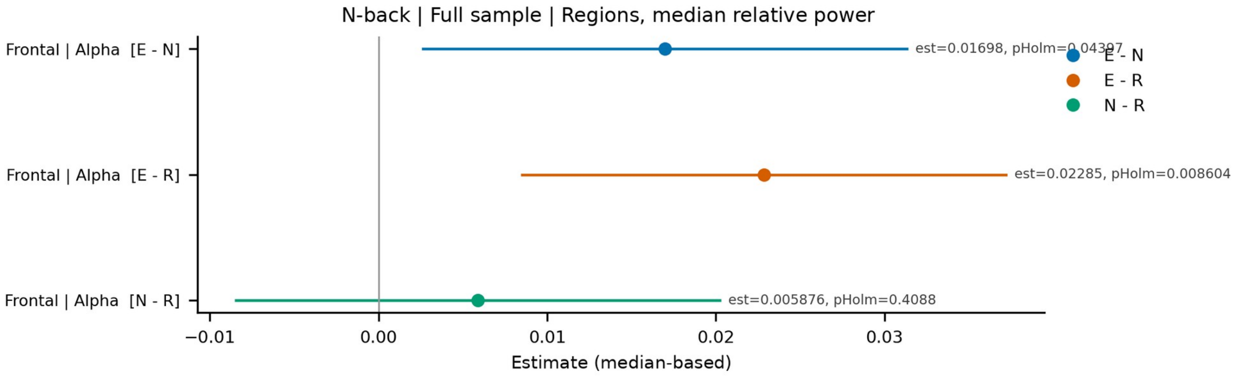
N-back | Full sample | Regions, median relative power
Frontal | Alpha [E - N]
Frontal | Alpha [E - R]
Frontal | Alpha [N - R]
est=0.01698, pHolm=0.04397
est=0.02285, pHolm=0.008604
est=0.005876, pHolm=0.4088
E - N
E - R
N - R
−0.01
0.00
0.01
0.02
0.03
Estimate (median-based)

**f**

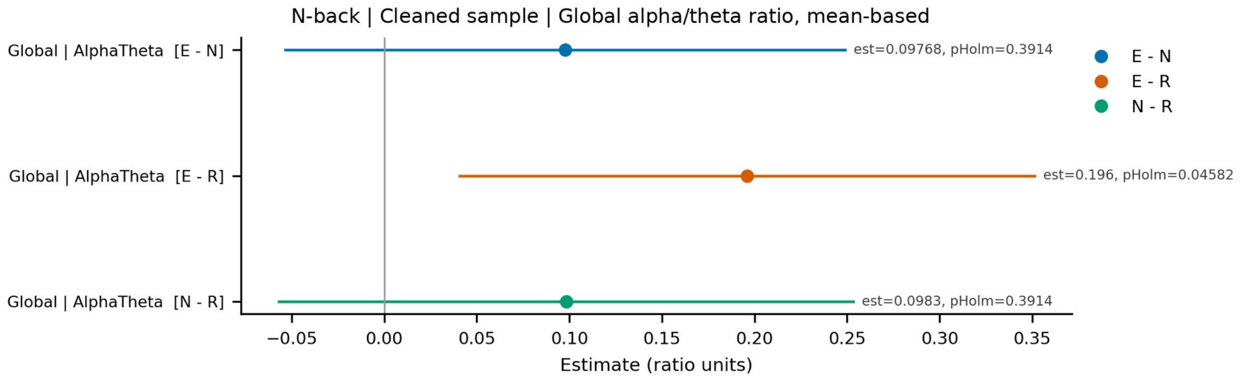
N-back | Cleaned sample | Global alpha/theta ratio, mean-based
Global | AlphaTheta [E - N]
Global | AlphaTheta [E - R]
Global | AlphaTheta [N - R]
est=0.09768, pHolm=0.3914
est=0.196, pHolm=0.04582
est=0.0983, pHolm=0.3914
E - N
E - R
N - R
−0.05
0.00
0.05
0.10
0.15
0.20
0.25
0.30
0.35
Estimate (ratio units)

**g**

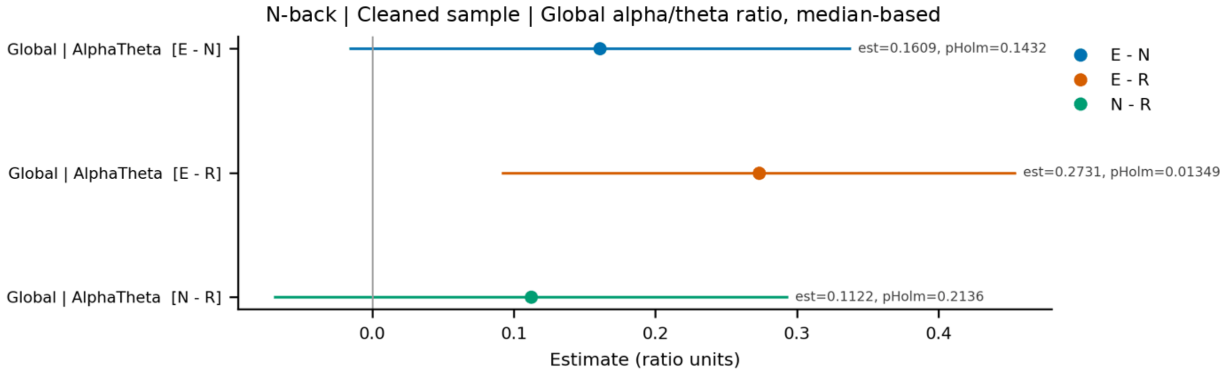
N-back | Cleaned sample | Global alpha/theta ratio, median-based
Global | AlphaTheta [E - N]
Global | AlphaTheta [E - R]
Global | AlphaTheta [N - R]
est=0.1609, pHolm=0.1432
est=0.2731, pHolm=0.01349
est=0.1122, pHolm=0.2136
E - N
E - R
N - R
0.0
0.1
0.2
0.3
0.4
Estimate (ratio units)

**h**

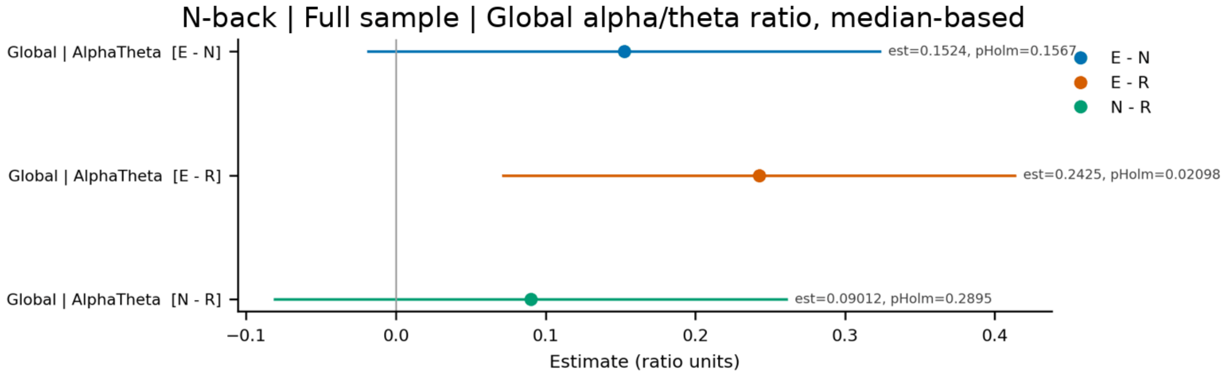
N-back | Full sample | Global alpha/theta ratio, median-based
Global | AlphaTheta [E - N]
Global | AlphaTheta [E - R]
Global | AlphaTheta [N - R]
est=0.1524, pHolm=0.1567
est=0.2425, pHolm=0.02098
est=0.09012, pHolm=0.2895
E - N
E - R
N - R
−0.1
0.0
0.1
0.2
0.3
0.4
Estimate (ratio units)

**i**

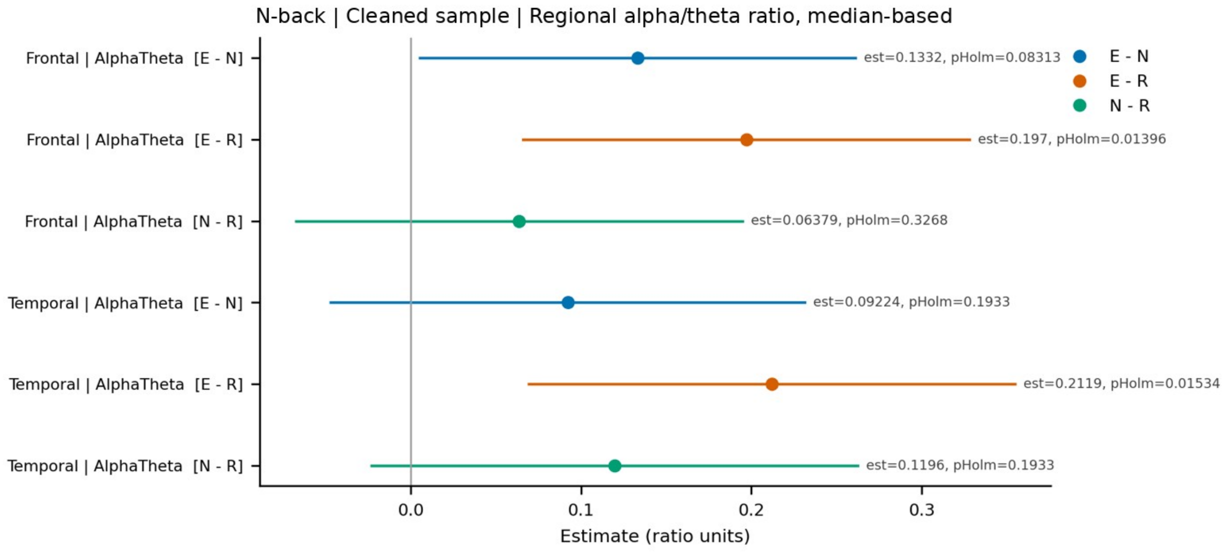

N-back | Cleaned sample | Regional alpha/theta ratio, median-based
Frontal | AlphaTheta [E - N]
Frontal | AlphaTheta [E - R]
Frontal | AlphaTheta [N - R]
Temporal | AlphaTheta [E - N]
Temporal | AlphaTheta [E - R]
Temporal | AlphaTheta [N - R]
est=0.1332, pHolm=0.08313
est=0.197, pHolm=0.01396
est=0.06379, pHolm=0.3268
est=0.09224, pHolm=0.1933
est=0.2119, pHolm=0.01534
est=0.1196, pHolm=0.1933
E - N
E - R
N - R
0.0
0.1
0.2
0.3
Estimate (ratio units)


**j**

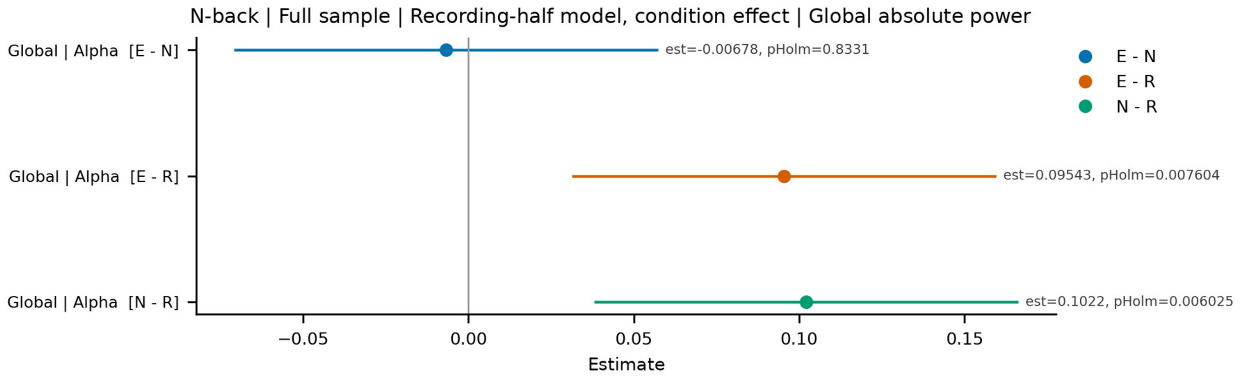

N-back | Full sample | Recording-half model, condition effect | Global absolute power
Global | Alpha [E - N]
Global | Alpha [E - R]
Global | Alpha [N - R]
est=-0.00678, pHolm=0.8331
est=0.09543, pHolm=0.007604
est=0.1022, pHolm=0.006025
E - N
E - R
N - R
−0.05
0.00
0.05
0.10
0.15
Estimate


**k**

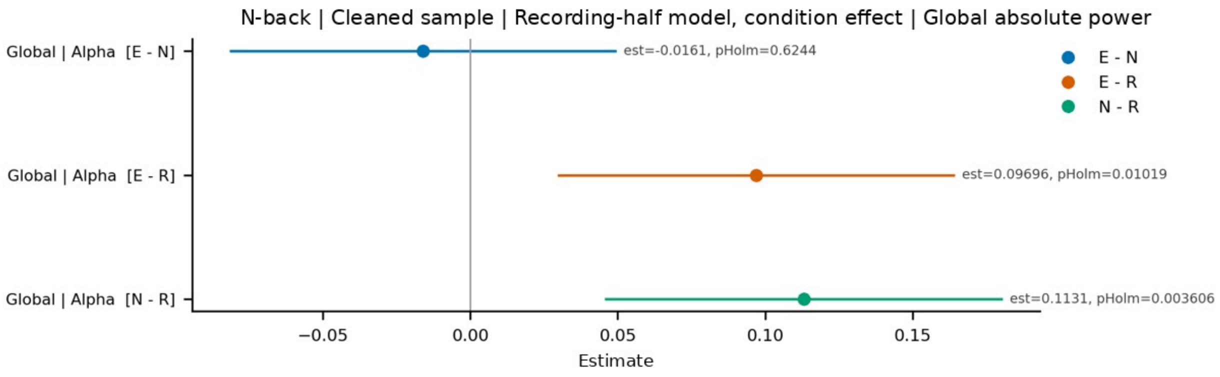

N-back | Cleaned sample | Recording-half model, condition effect | Global absolute power
Global | Alpha [E - N]
Global | Alpha [E - R]
Global | Alpha [N - R]
est=-0.0161, pHolm=0.6244
est=0.09696, pHolm=0.01019
est=0.1131, pHolm=0.003606
E - N
E - R
N - R
−0.05
0.00
0.05
0.10
0.15
Estimate

**l**

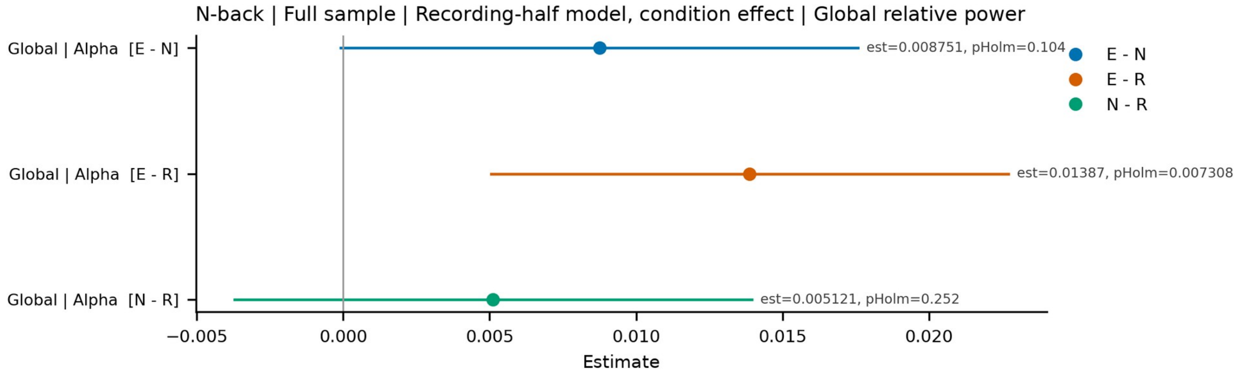
N-back | Full sample | Recording-half model, condition effect | Global relative power
Global | Alpha [E - N]
Global | Alpha [E - R]
Global | Alpha [N - R]
est=0.008751, pHolm=0.104
est=0.01387, pHolm=0.007308
est=0.005121, pHolm=0.252
E - N
E - R
N - R
−0.005
0.000
0.005
0.010
0.015
0.020
Estimate

**m**

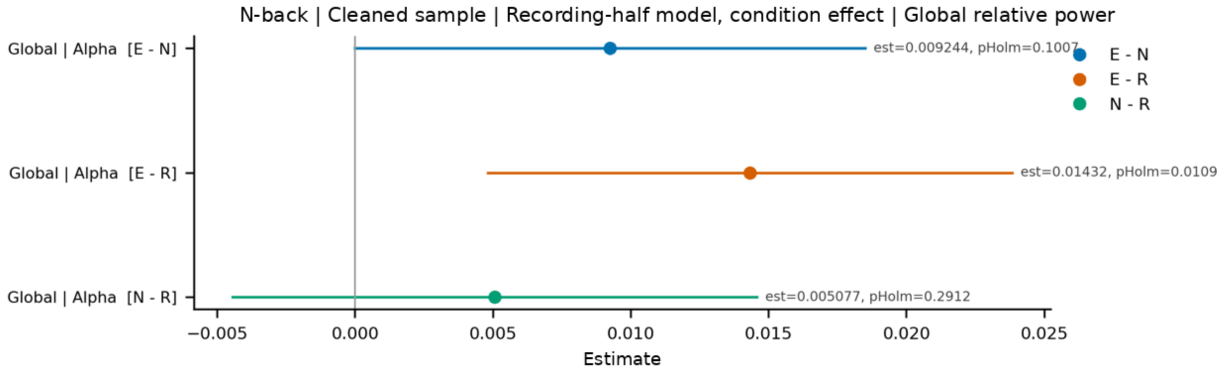
N-back | Cleaned sample | Recording-half model, condition effect | Global relative power
Global | Alpha [E - N]
Global | Alpha [E - R]
Global | Alpha [N - R]
est=0.009244, pHolm=0.1007
est=0.01432, pHolm=0.0109
est=0.005077, pHolm=0.2912
E - N
E - R
N - R
−0.005
0.000
0.005
0.010
0.015
0.020
0.025
Estimate

**n**

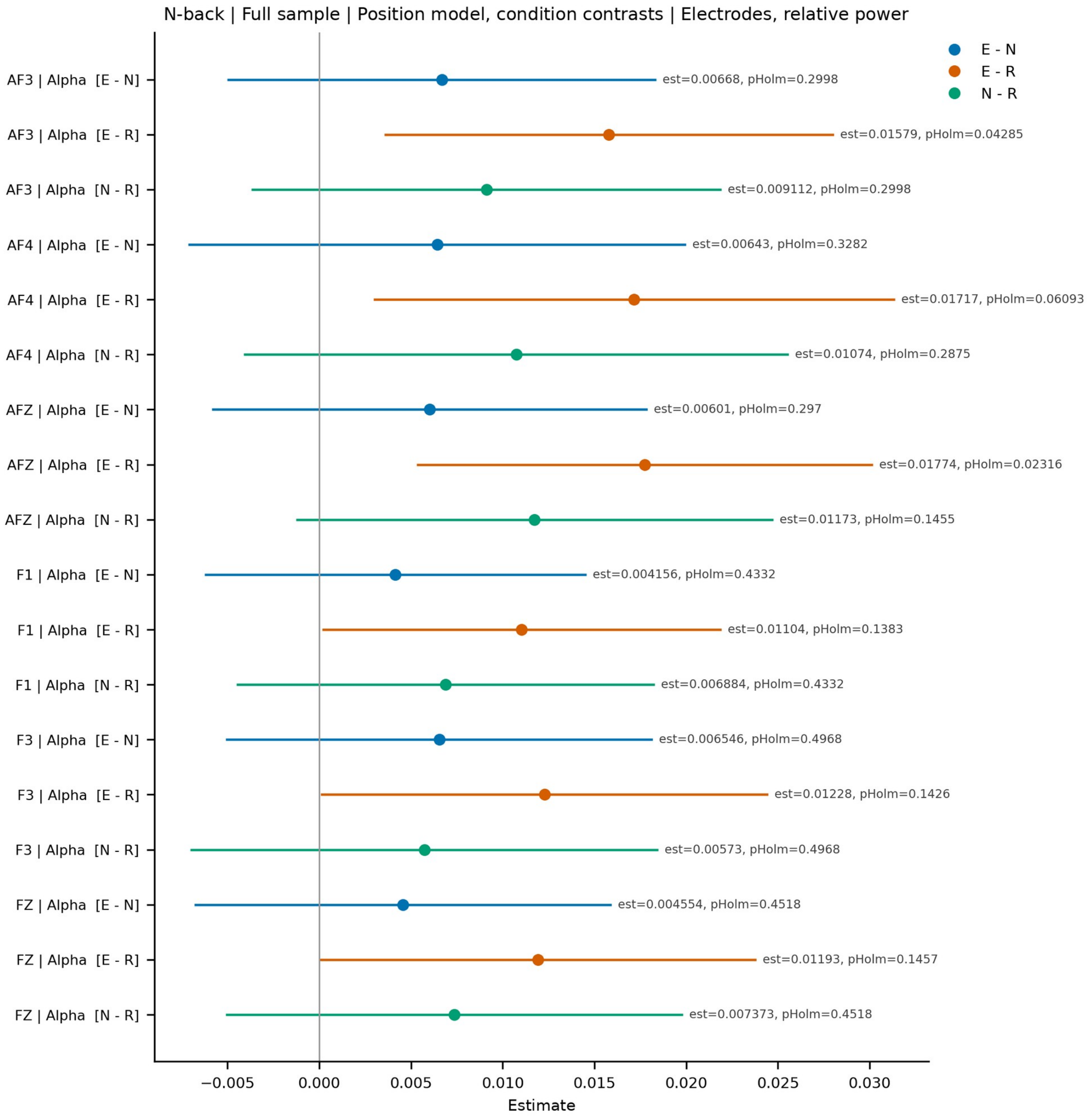
N-back | Full sample | Position model, condition contrasts | Electrodes, relative power
E - N
E - R
N - R
AF3 | Alpha [E - N]
AF3 | Alpha [E - R]
AF3 | Alpha [N - R]
AF4 | Alpha [E - N]
AF4 | Alpha [E - R]
AF4 | Alpha [N - R]
AFZ | Alpha [E - N]
AFZ | Alpha [E - R]
AFZ | Alpha [N - R]
F1 | Alpha [E - N]
F1 | Alpha [E - R]
F1 | Alpha [N - R]
F3 | Alpha [E - N]
F3 | Alpha [E - R]
F3 | Alpha [N - R]
FZ | Alpha [E - N]
FZ | Alpha [E - R]
FZ | Alpha [N - R]
est=0.00668, pHolm=0.2998
est=0.01579, pHolm=0.04285
est=0.009112, pHolm=0.2998
est=0.00643, pHolm=0.3282
est=0.01717, pHolm=0.06093
est=0.01074, pHolm=0.2875
est=0.00601, pHolm=0.297
est=0.01774, pHolm=0.02316
est=0.01173, pHolm=0.1455
est=0.004156, pHolm=0.4332
est=0.01104, pHolm=0.1383
est=0.006884, pHolm=0.4332
est=0.006546, pHolm=0.4968
est=0.01228, pHolm=0.1426
est=0.00573, pHolm=0.4968
est=0.004554, pHolm=0.4518
est=0.01193, pHolm=0.1457
est=0.007373, pHolm=0.4518
−0.005
0.000
0.005
0.010
0.015
0.020
0.025
0.030
Estimate

**o**

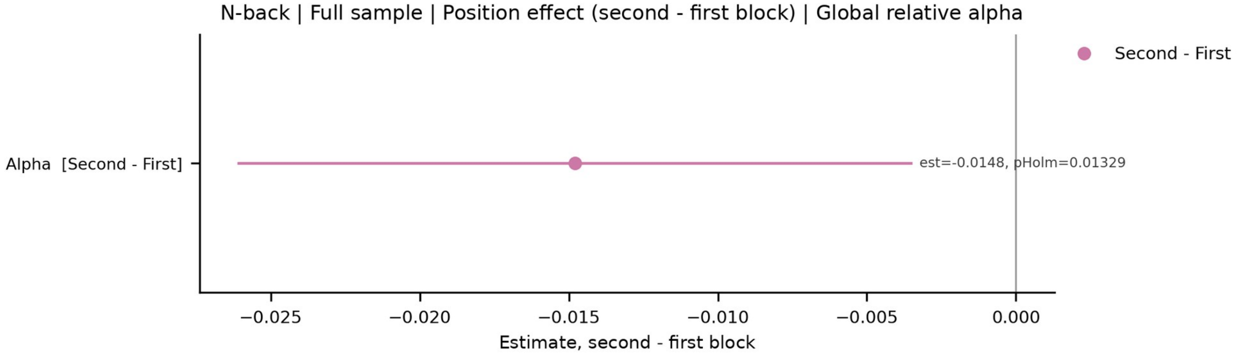
N-back | Full sample | Position effect (second - first block) | Global relative alpha
Second - First
Alpha [Second - First]
est=-0.0148, pHolm=0.01329
−0.025
−0.020
−0.015
−0.010
−0.005
0.000
Estimate, second - first block

**p**

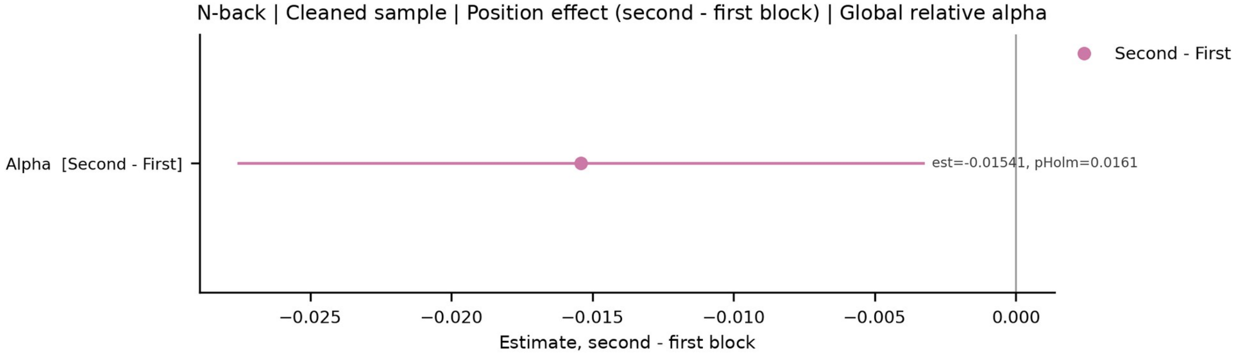
N-back | Cleaned sample | Position effect (second - first block) | Global relative alpha
Second - First
Alpha [Second - First]
est=-0.01541, pHolm=0.0161
−0.025
−0.020
−0.015
−0.010
−0.005
0.000
Estimate, second - first block

**q**

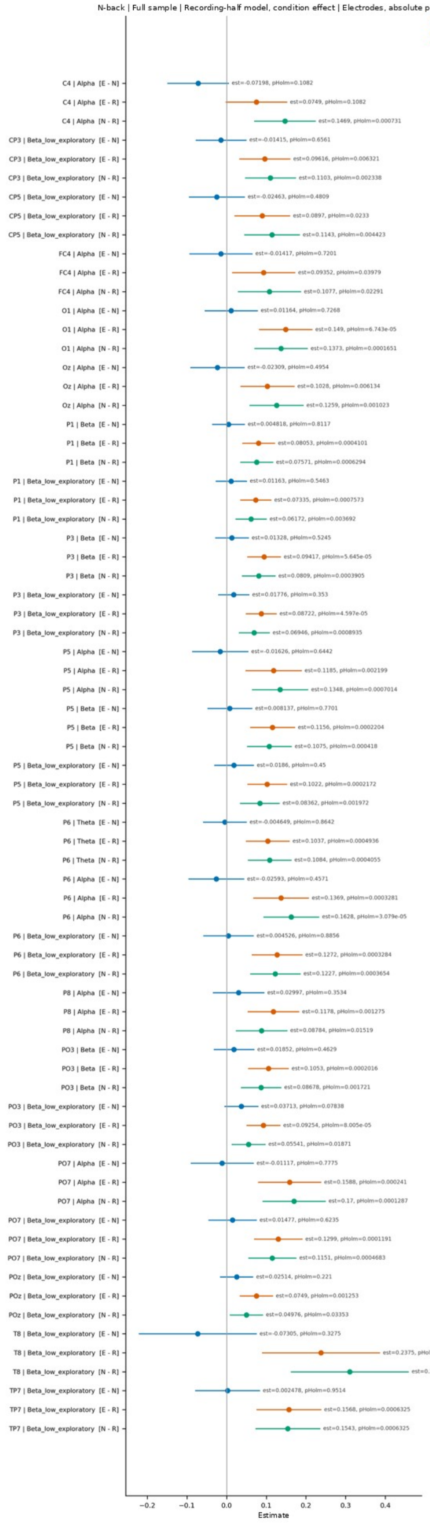
N-back | Full sample | Recording-half model, condition effect | Electrodes, absolute power
Estimate

**r**

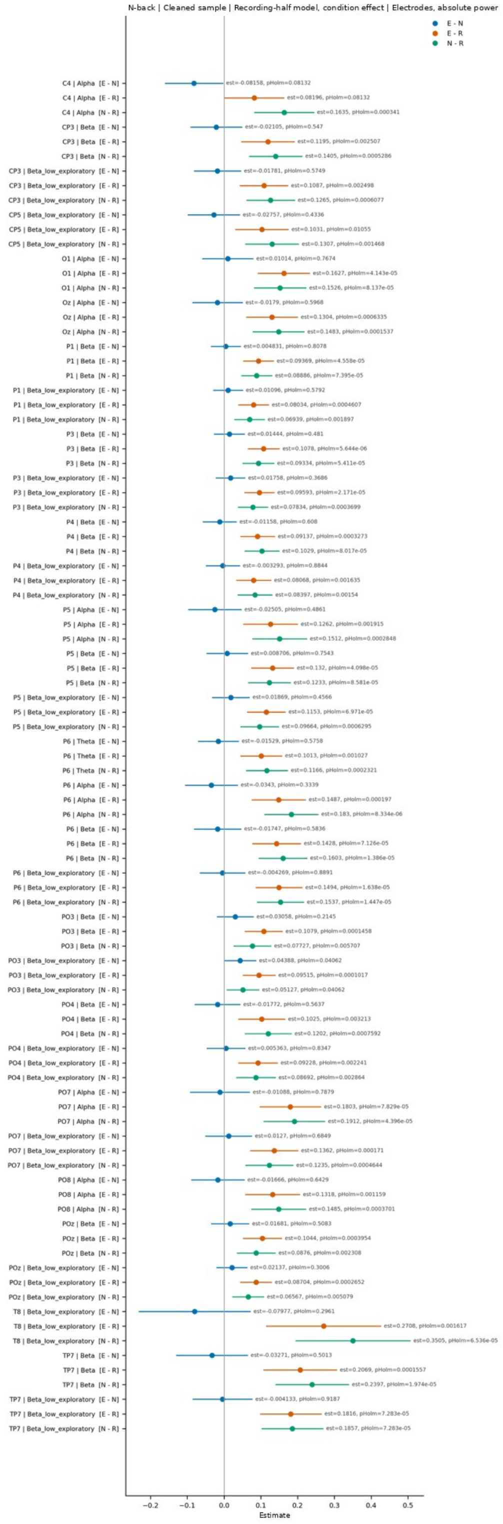
N-back | Cleaned sample | Recording-half model, condition effect | Electrodes, absolute power
Estimate

**s**

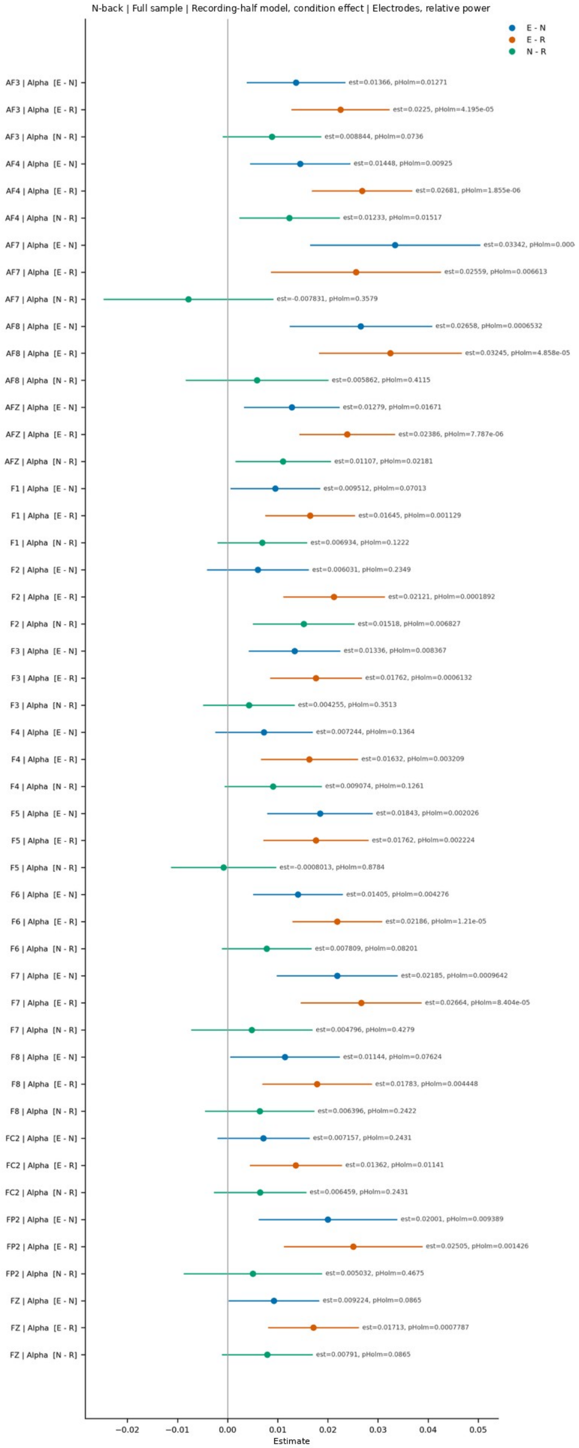
N-back | Full sample | Recording-half model, condition effect | Electrodes, relative power
E - N
E - R
N - R
Estimate

**t**

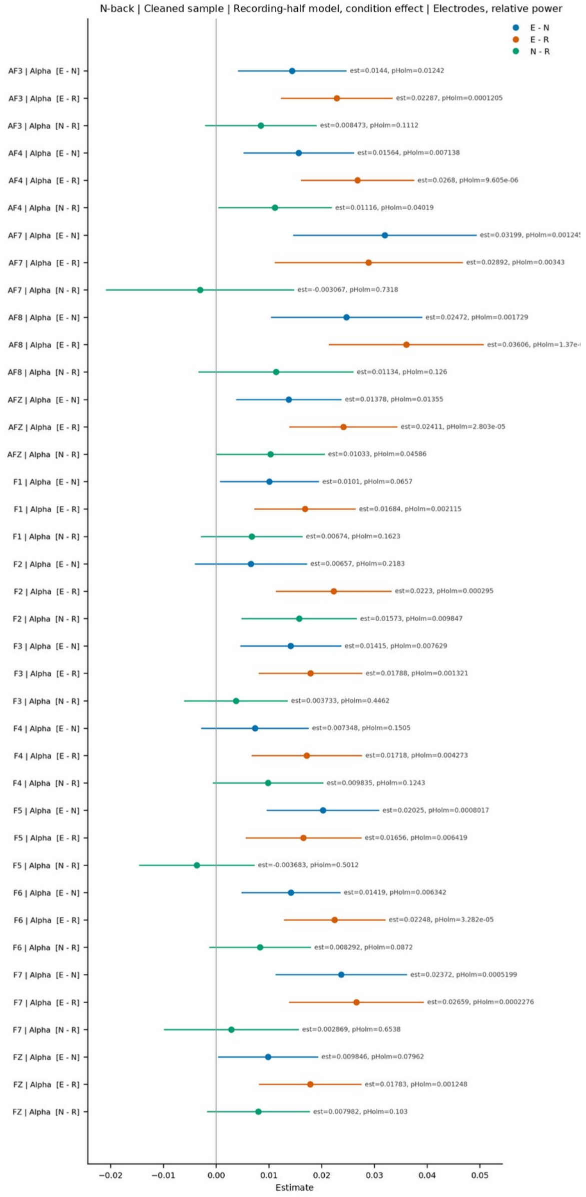
N-back | Cleaned sample | Recording-half model, condition effect | Electrodes, relative power
E - N
E - R
N - R
Estimate

**u**

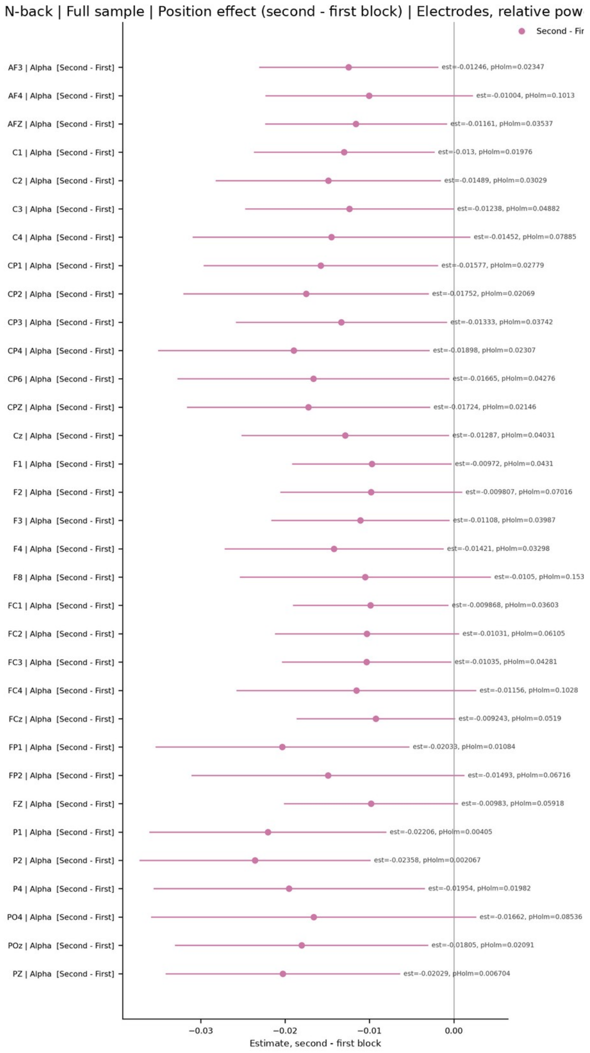


**v**

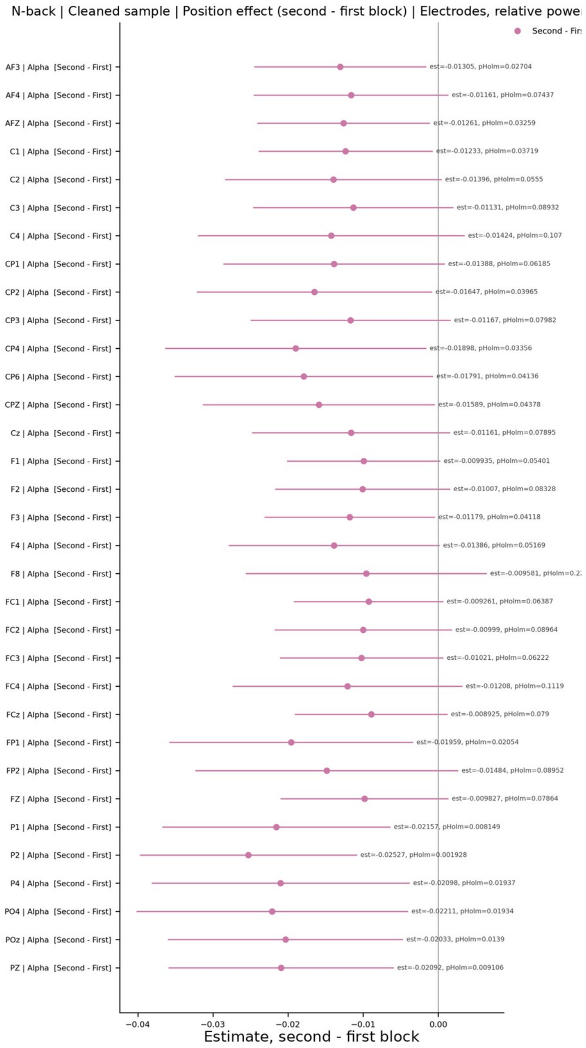


**Supplementary Fig. S24. Follow-up models, N-back.** Median-based power at the electrode and region levels (a-e), alpha/theta ratios (f-i), condition effects in the condition × recording-half model at the global level (j-m), condition contrasts in the intervention-position model (n), the position effect on global relative alpha (o, p), electrode-level condition effects in the recording-half model for absolute (q, r) and relative power (s, t), and the electrode-level position effect on relative power (u, v). Panel titles give the task, sample, model, and measure. Each row shows a contrast (E − N, E − R, or N − R; for position effects, second minus first post-intervention block) with its 95% CI; labels give the estimate and the Holm-corrected *p*. Only units whose omnibus test survived FDR correction are shown. Beta_low, Beta_high, Gamma_low, and Gamma_high1 denote the exploratory 13-20, 20-30, 30-45, and 55-80 Hz bands.

**a**

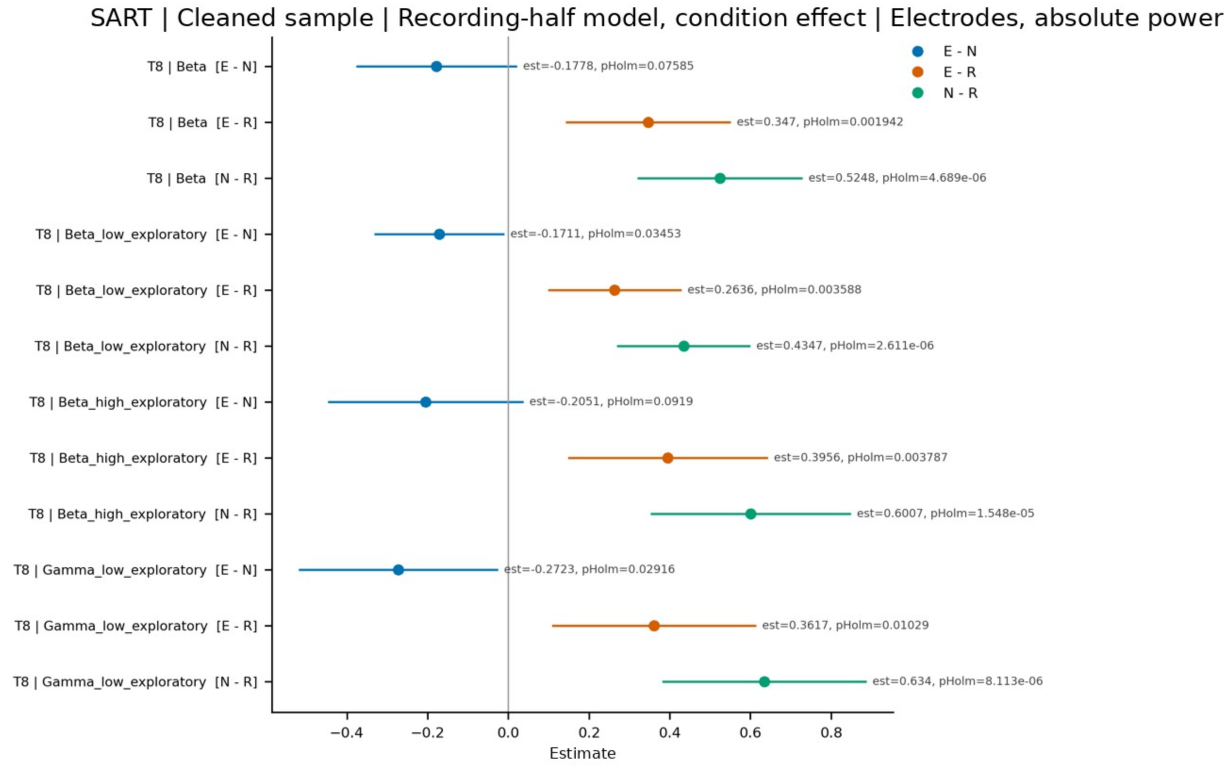
SART | Cleaned sample | Recording-half model, condition effect | Electrodes, absolute power
T8 | Beta [E - N]
T8 | Beta [E - R]
T8 | Beta [N - R]
T8 | Beta_low_exploratory [E - N]
T8 | Beta_low_exploratory [E - R]
T8 | Beta_low_exploratory [N - R]
T8 | Beta_high_exploratory [E - N]
T8 | Beta_high_exploratory [E - R]
T8 | Beta_high_exploratory [N - R]
T8 | Gamma_low_exploratory [E - N]
T8 | Gamma_low_exploratory [E - R]
T8 | Gamma_low_exploratory [N - R]
est=-0.1778, pHolm=0.07585
est=0.347, pHolm=0.001942
est=0.5248, pHolm=4.689e-06
est=-0.1711, pHolm=0.03453
est=0.2636, pHolm=0.003588
est=0.4347, pHolm=2.611e-06
est=-0.2051, pHolm=0.0919
est=0.3956, pHolm=0.003787
est=0.6007, pHolm=1.548e-05
est=-0.2723, pHolm=0.02916
est=0.3617, pHolm=0.01029
est=0.634, pHolm=8.113e-06
E - N
E - R
N - R
−0.4
−0.2
0.0
0.2
0.4
0.6
0.8
Estimate

**b**

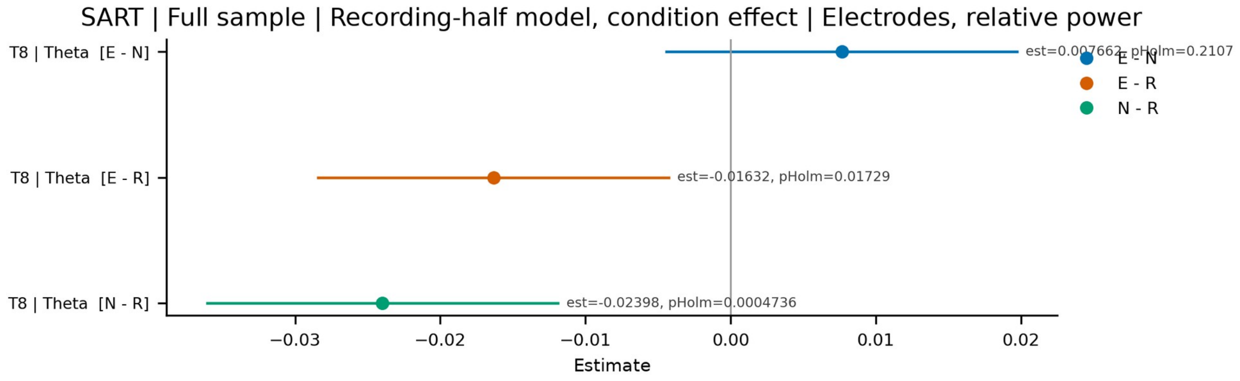
SART | Full sample | Recording-half model, condition effect | Electrodes, relative power
T8 | Theta [E - N]
T8 | Theta [E - R]
T8 | Theta [N - R]
est=0.007662, pHolm=0.2107
est=-0.01632, pHolm=0.01729
est=-0.02398, pHolm=0.0004736
E - N
E - R
N - R
−0.03
−0.02
−0.01
0.00
0.01
0.02
Estimate

**c**

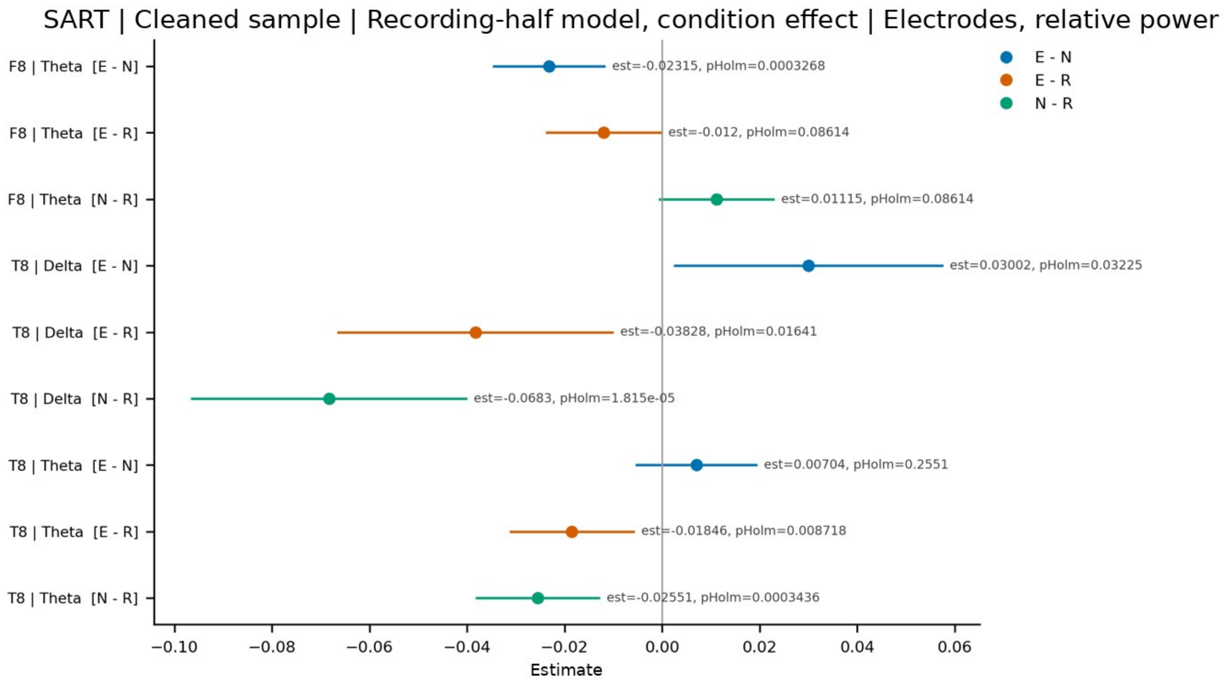


**d**

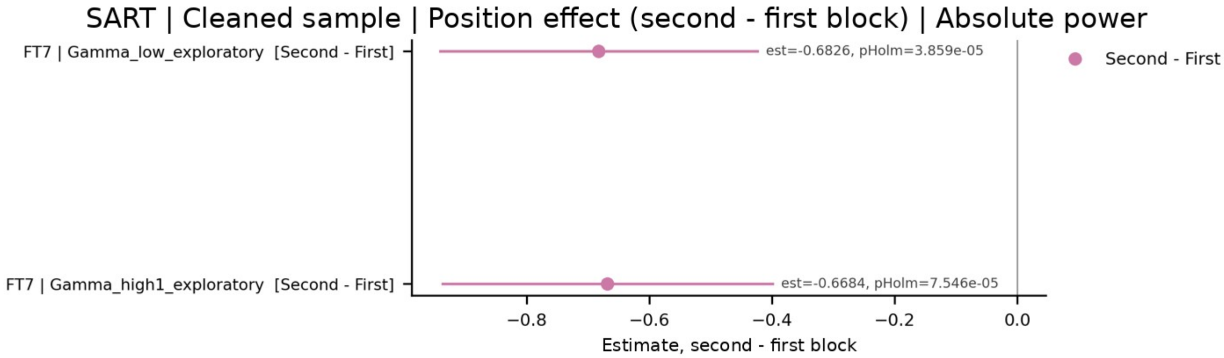


**Supplementary Fig. S25. Follow-up models, SART.** Condition effects in the condition × recording-half model at the electrode level for absolute (a) and relative power (b, c), and the position effect on absolute power (d). Panel titles give the sample, model, and measure. Each row shows a contrast (E − N, E − R, or N − R; for position effects, second minus first post-intervention block) with its 95% CI; labels give the estimate and the Holm-corrected *p*. Only units whose omnibus test survived FDR correction are shown. Beta_low, Beta_high, Gamma_low, and Gamma_high1 denote the exploratory 13-20, 20-30, 30-45, and 55-80 Hz bands.
